# Diet induces reproducible alterations in the mouse and human gut microbiome

**DOI:** 10.1101/541797

**Authors:** Jordan E. Bisanz, Vaibhav Upadhyay, Jessie A. Turnbaugh, Kimberly Ly, Peter J. Turnbaugh

## Abstract

The degree to which diet reproducibly alters the human and mouse gut microbiota remains unclear. Here, we focus on the consumption of a high-fat diet (HFD), one of the most frequently studied dietary interventions in mice. We employed a subject-level meta-analysis framework for unbiased collection and analysis of publicly available 16S rRNA gene and metagenomic sequencing data from studies examining HFD in rodent models. In total, we re-analyzed 27 studies, 1101 samples, and 106 million reads mapping to 16S rRNA gene sequences. We report reproducible changes in gut microbial community structure both within and between studies, including a significant increase in the Firmicutes phylum and decrease in the Bacteroidetes phylum; however, reduced alpha diversity is not a consistent feature of HFD. Finer taxonomic analysis revealed that the strongest signal of HFD on microbiota species composition is *Lactococcus* spp., which we demonstrate is a common dietary contaminant through the molecular testing of dietary ingredients, culturing, microscopy, and germ-free mouse experiments. After *in silico* removal of *Lactococcus* spp., we employed machine learning to define a unique operational taxonomic unit (OTU)-based signature capable of predicting the dietary intake of mice and demonstrate that phylogenetic and gene-family transformations of this model are capable of accurately predicting human samples in controlled feeding settings (area under the receiver operator curve = 0.75 and 0.88 respectively). Together, these results demonstrate the utility of microbiome meta-analyses in identifying robust bacterial signals for mechanistic studies and creates a framework for the routine meta-analysis of microbiome studies in preclinical models.

## Introduction

Numerous studies have evaluated the impact of macronutrient intake on community composition of the distal gut microbiota (Turnbaugh, 2017). Perhaps, the most often-studied dietary intervention is the consumption of a high-fat diet (HFD) given evidence for a causal role of HFD-induced shifts in the gut microbiota in multiple disease models (Turnbaugh et al., 2008; Upadhyay et al., 2012). Despite the use of murine hosts which might be expected to be have a more homogeneous gut microbiota than human populations, many reports are qualitative in nature, and a lack of quantitative definition directly limits understanding of what signals are reproducible across studies.

This lack of definition is emblematic of the broader reproducibility crisis in scientific literature (Baker, 2016), wherein technical and/or biological inconsistencies between studies can complicate interpretation of the effect, or lack thereof, of diet on a given phenotype of interest (in this particular case, gut microbial community structure). On shallow examination, many studies may appear to examine the same experimental variables and outputs; however, this is often not the case. Differences between studies likely reflect a complex interaction between the specific diet formulations used, the host species (e.g., mouse vs. rat), colonization status (conventional vs. gnotobiotic), animal vendor, and technical differences in DNA extraction, sample handling, sequencing, and/or computational methods (Sinha et al., 2017). Scientific meta-analysis can help to address these discrepancies in an unbiased manner, providing a stronger foundation for follow-on studies (Gurevitch et al., 2018).

Here, we present the results of a large-scale meta-analysis of amplicon- and metagenomic-sequencing based studies investigating the effect of HFD on the gut microbiome in both conventional and humanized murine models. We also re-analyze data from two human dietary intervention studies (Wu et al., 2011; David et al., 2013). Despite major differences in experimental design between studies, we are able to identify microbial signatures that are consistent and predictive of HFD intake. Utilizing consistent computational tools across multiple data sets, this analysis partially accounts for inter-study differences described above and also employs the statistical power of the collected studies en masse to define a reproducible molecular signal indicative of the rodent response to HFD feeding and demonstrates translatability to humans.

## Results

### Study selection and characteristics

A total of 427 unique studies were retrieved by our search methodology (**Figure 1A**). 11 reviewers working in the microbiome field, and who were familiar with the terminology and methodology of the microbiome field, redundantly reviewed the studies to determine the relevance of study design and methodology for inclusion (*see Methods*). Of these 427 studies, 79 studies were chosen through a web-based crowd-sourced consensus to be eligible for this meta-analysis. Of these 79 studies, 44 lacked clear information regarding a specific, public location of sequencing data. Of the remaining 35 studies, 10 lacked complete metadata sufficient for pairing sequencing data to sample information. This left 25 murine studies for inclusion in our meta-analysis. Two additional human studies were identified to examine translatability of mouse studies to humans (Wu et al., 2011; David et al., 2013). Relevant per-study descriptions and metadata are listed in **Table S1**. These studies encompassed 1073 murine samples (477 HFD, 596 LFD), and 29 human samples (14 HFD, 15 LFD). Per sample metadata is provided in **Table S2**.

**Figure 1.**
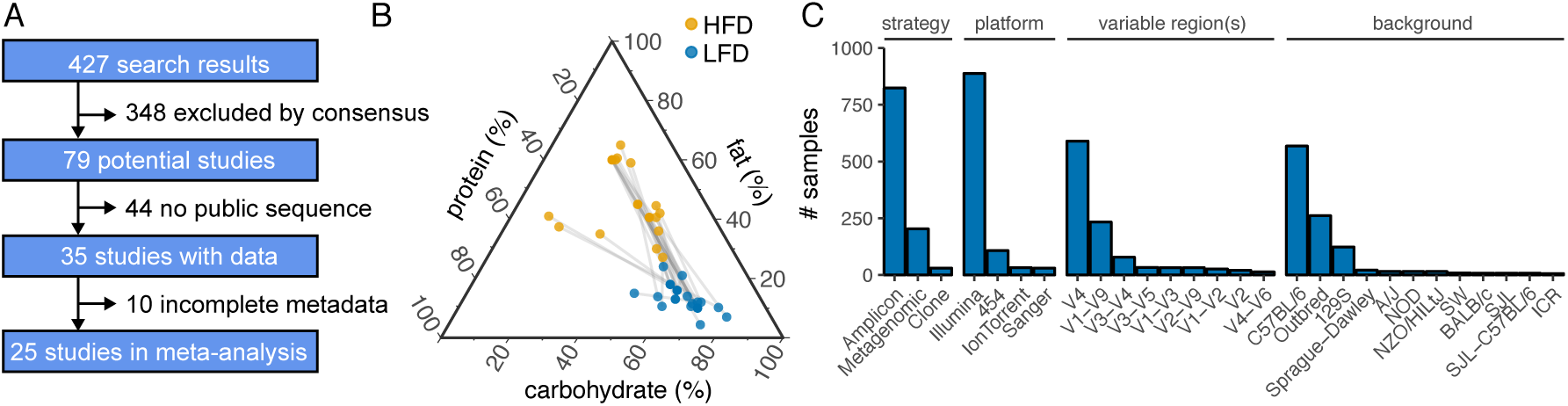
Study characteristics. **(A)** A total of 427 studies were identified for potential inclusion of which 25 murine studies were ultimately included. **(B)** Macronutrient composition (by % kCal) of studied HFD and LFD diets demonstrates diets are generally isocaloric in protein content while there is great variability in what fat content is considered high (linked diets represent those used in the same study). **(C)** Sample breakdown by sequencing strategy, platform, variable region and host strain demonstrate that Illumina sequencing of the V4 hypervariable region of the 16S rRNA gene in C57BL/6 hosts are most common.

Studies varied significantly by dietary fat content, immediately suggesting that a portion of variability is likely due to lack of specificity surrounding terminology. The range of reported dietary fat across all groups was 4.4% kCal to 65% kCal (**Figure 1B**). Furthermore, the range of fat composition that constituted a LFD or a control diet and a HFD varied significantly. The highest fat content of LFD fed animals was 24% of kCal from fat (range 4.4%-24%) but this nearly overlapped the lower bound of the range of HFD which was 27.1% (range 27.1%-65%). The majority of studies varied dietary fat at the expense of carbohydrate content (range 11-80%). On the other hand, protein content was more consistent across most studies with few outliers (range 13%-48%, **Figure 1B**).

Studies also varied considerably in technical considerations including: sequencing strategy, platform, variable regions, and host strain (**Figure 1C**, Figure S1A), in addition to study size and sequencing depth (**Figure S1B**). The majority of samples were derived from outbred and C57BL/6 mice profiled via V4 16S rRNA amplicon sequencing on an Illumina-platform sequencer. Raw reads were obtained on a per sample basis for 1102 samples (**Table S2**) from the NCBI Sequence Read Archive (SRA) and/or MGRAST where they were consistently processed for closed-reference OUT picking against the 13-8 Greengenes release. A total of 29,937 OTUs were observed across all samples before any form of filtering or quality control.

### Effect of HFD on microbial diversity

We calculated commonly used metrics for alpha diversity: Chao1 richness, Shannon’s diversity, and Faith’s phylogenetic diversity, as well as beta diversity: Bray-Curtis dissimilarity, weighted/unweighted UniFrac, Jensen-Shannon divergence, PhILR Euclidean distance (Silverman et al., 2017), and CLR Euclidean distance (i.e. Aitchison distance, Gloor et al. (2017)). The ratio of the phyla Firmicutes and Bacteroidetes was also calculated due to its frequent use in the literature. To visualize the data and account for varied baseline states, all values were scaled to the geometric mean of LFD samples on a per-study basis. In visualizing these metrics, it is apparent that altered alpha diversity is not a consistent feature of HFD across all studies with considerable heterogeneity in the direction of effect observed **(Figure 2A-C)**. Considering all studies together, there is a modest significant decrease in Chao1 richness (−0.215 [-0.270 to −0.159], P=8.73e-14), Shannon’s diversity (−0.048 [-0.084 to −0.013], P=7.83e-3), and Faith’s phylogenetic diversity (−0.122 [-0.157 to −0.088], P=7.26e-12) (log2(fold change) [95% CI]). To consider dietary fat content as a continuous variable, we also examined the correlation between fat content and diversity (**Figure S2**), finding no meaningful relationship (P 0.05 Spearman’s Correlation).

**Figure 2.**
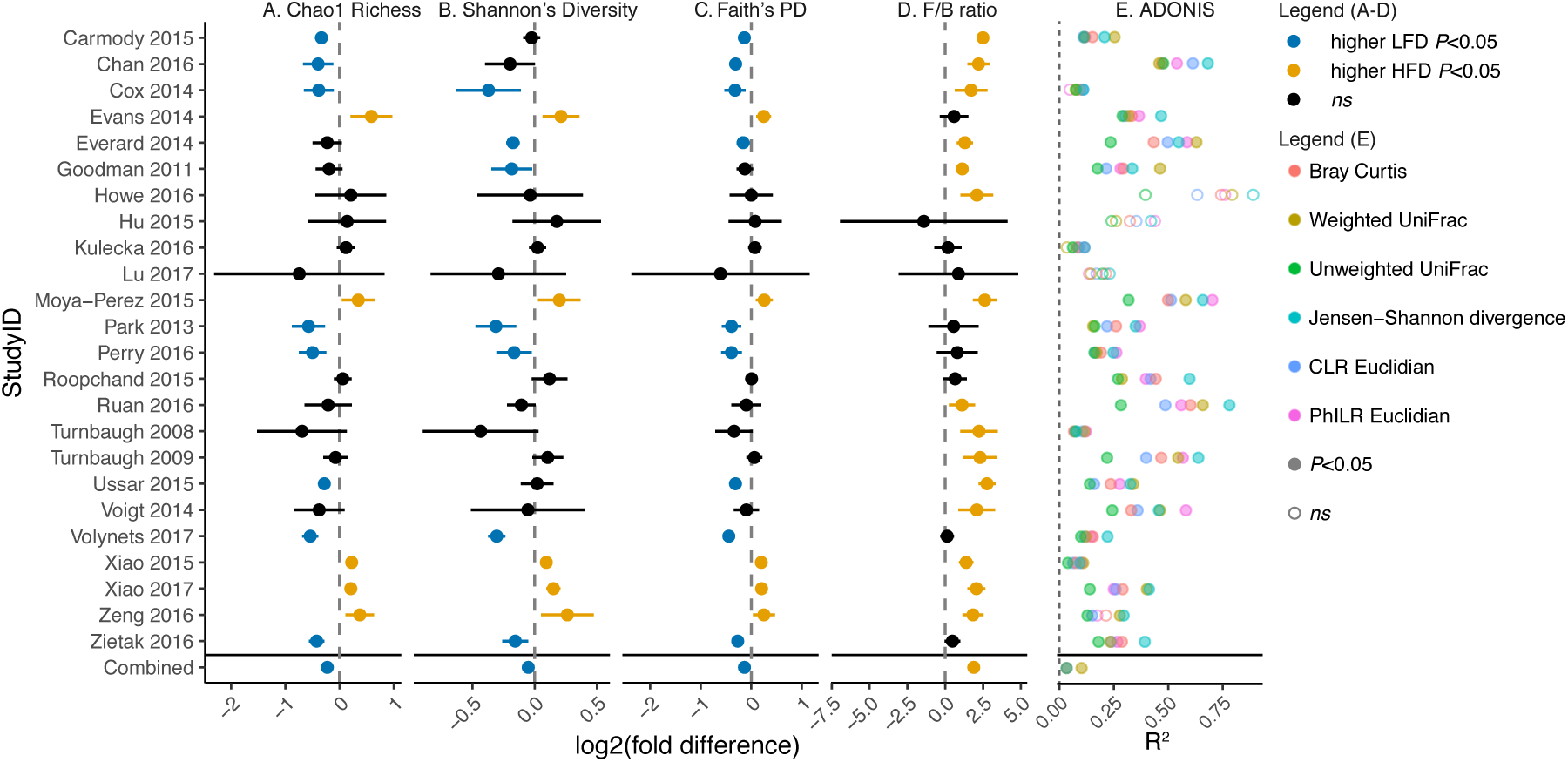
Forest plot of commonly described community metrics in murine studies. Measures of alpha diversity including **(A)** Chao1 richness, **(B)** Shannon’s diversity, and **(C)** Faith’s phylogenetic diversity demonstrate inconsistent effects of diet between studies. **(D)** The ratio of the Firmicutes phylum to the Bacteroidetes phylum is a consistent feature of the HFD. Points represent mean and 95% confidence interval (panels A-D, individual studies computed by Welch’s t-test and combined by linear mixed effects model with study as random effect). **(E)** ADONIS tests of various distance metrics demonstrates consistent within-study effects on community composition as measured by the % variation explained (*R*^2^) by diet classification (Combined withstudy as stratum).

Next, the ratio of Firmicutes to Bacteroidetes was calculated and found to be consistently increased in 15 of 25 murine studies, which was supported statistically when all studies were considered in aggregate (*log*_2_*FC*=1.84 [1.65 to 2.03], P=3.4e-69). This trend is even apparent when visualized in the commonly reported phylum-level bar plot (**Figure S3**).

### HFD reproducibly alters community composition

We next employed visualization strategies and statistically tested the effect of HFD on community composition as a whole using principal coordinates analysis of multiple distance metrics with statistical testing via ADONIS (analysis of variance using distance matrices, (**Figure S4**, **Figure 2E**). With the exception of Howe 2016, Hu 2015, and Lu 2017 which suffer from a lack of power (n=3/group each), all remaining studies demonstrated a significant effect of diet on community composition (P*<*0.05, ADONIS), albeit with variable variance explained ranging from 0.035 to 0.891 (R^2^). All studies were aggregated and shallowly subsampled (112 reads) to overcome extreme discrepancies in read depth (**Figure** S1). Due to matrix sparsity (96.7% zero-observations, **Figure S5A-B**), significant distance saturation was observed (**Figure S6**) so only phylogeny-aware metrics were employed: weighted and unweighted UniFrac, and PhILR Euclidean (**Figure 3**, associated scree plot **Figure S7A**). Clear visual clustering independent of study was observed; which was supported by ADONIS (P*<*0.001) for weighted and unweighted UniFrac, and PhILR Euclidean data types (R^2^=0.102, 0.033, 0.0335 respectively). The interstudy-variation outweighed the effect of diet (R^2^=0.198 to 0.487, **Table S3**). Given the clear evidence for an underlying HFD-signal from multivariate analyses, we sought to further understand which specific features of the microbiota are responsive to diet intervention.

**Figure 3.**
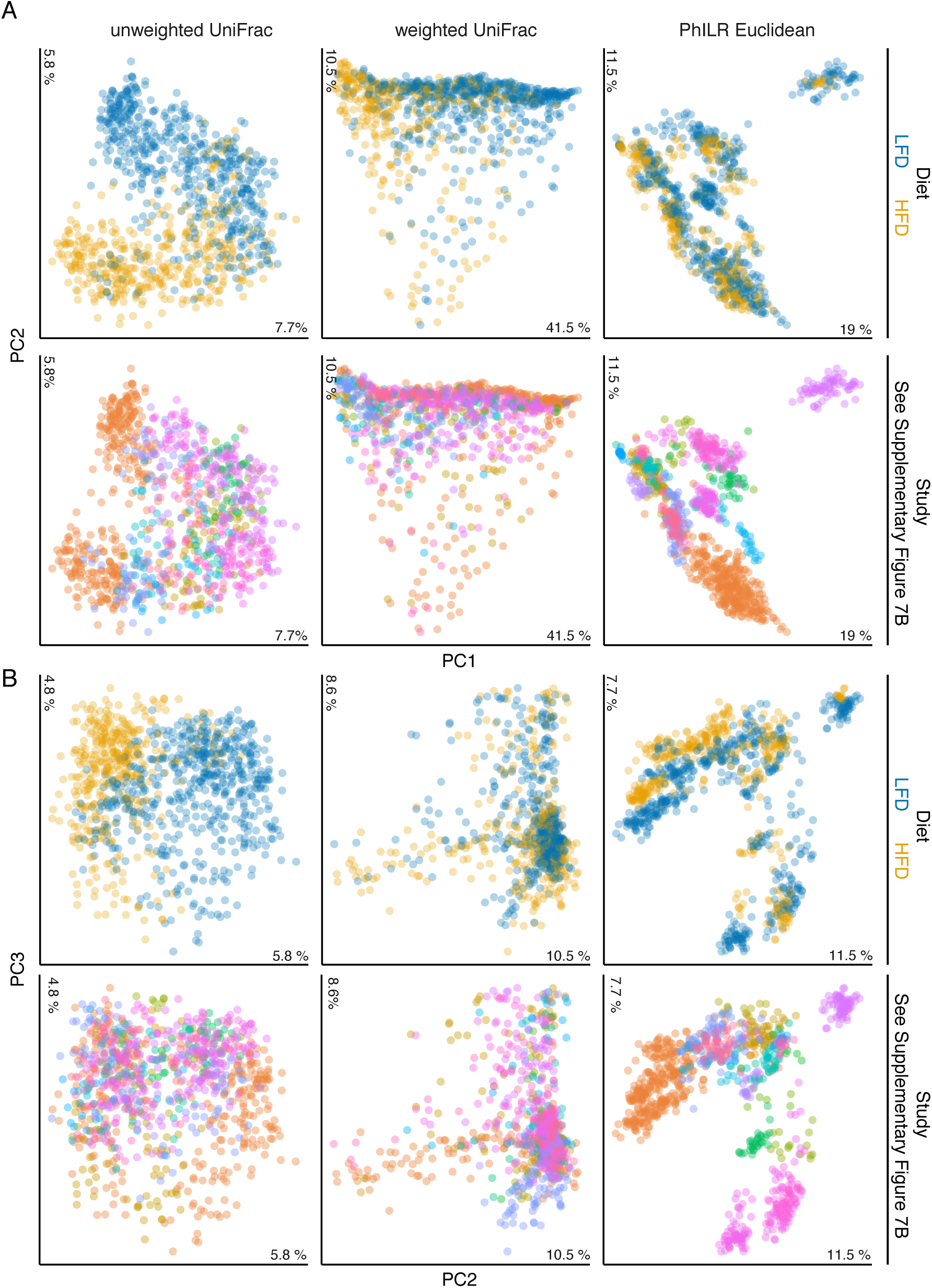
Principal coordinate analysis of murine samples by diet and study. **(A)** Axis 1 versus 2 and **(B)** axis 2 versus 3 provide clear visual evidence for a significant effect of study as well as an effect of diet composition (n=978 samples, P*<*0.001 ADONIS; **Table S3**).

### Predictive microbial responses to HFD

We used a random forest classifier to define reliable biomarkers of the gut microbial response to HFD. To avoid issues related to overfitting, we established 5 groups for training and sequential validation: a stepwise approach was used wherein murine samples were randomly selected with two thirds of the resulting set used for training (Murine Training Set n=569) and one third for validation (Murine Test Set n=284). Next the 3 randomly selected external validation sets were predicted whose samples were not used to inform the initial model (Everard 2014, Xiao 2015, and Evans 2014, External Murine Sample n=173) followed by Humanized Mice (n=46), and Human (n=29) samples. A summary of these datasets is provided in **Table S4**.

Using only the Murine Training Set, 10-fold cross validation was applied to determine the optimal number of features included in the model required to minimize error rates in the training set. We noted that with even with as few as 4 OTUs (**Figure 4A**), classification error rates of *<*15% could be obtained, emphasizing the high predictive power of the top features. To visualize these, a phylogenetic tree of the 229 most-informative OTUs was created (**Figure 4B**). The most predictive OTUs belonged to the genus *Lactococcus* based on mean decrease in GINI coefficient (**Figure 5A**) which was significantly elevated in 14 of the examined murine studies (**Figure 5B**). Given that the particular OTUs in question (97% OTUs 716006, 571744, and 4468805) mapped to *Lactococcus lactis*, we reasoned that this signal may represent latent contamination in the food rather than a response of the microbiota itself. While we and others have previously reported this possibility (Carmody et al., 2015; Dalby et al., 2017; Dollive et al., 2013), this highly reproducible finding across multiple labs and studies drove us to definitively test this hypothesis.

**Figure 4.**
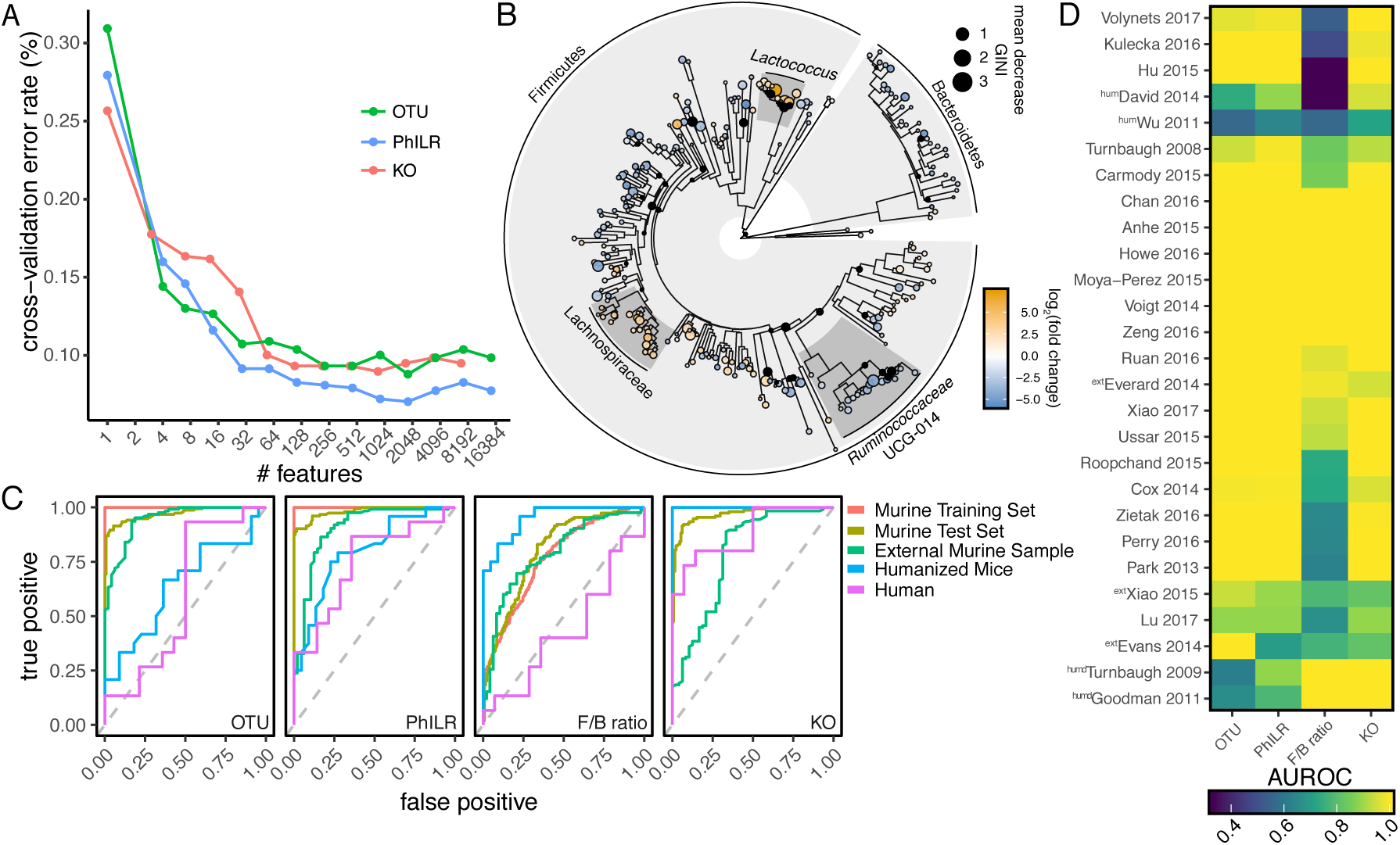
Identification of reproducible microbial signatures of the HFD-associated gut micro-biota. **(A)** 10-fold cross validation for feature selection identifies a minimum core set of informative features. **(B)** Phylogenetic tree of informative OTUs (n=229) demonstrates highly informative clades of *Lactococcus, Ruminococcaceae* UCG-014 and *Lachnospiraceae*. Size of circle correlates with mean decrease GINI coefficient and data are colored by log2(fold change). **(C)** Receiver operator curves for *Lactococcus*-depleted OTUs, phylogenetic node balances (PhILR), and KEGG orthologies (KO). Area under the receiver operator curve (AUROC) are provided in **Table S6**. **(D)** Area under the receiver operator curve (AUROC) by study demonstrates that PhILR and KO models are generalizable to most studies (abbreviations: ext - external murine sample, hum-human sample, humd-humanized murine sample). Rows are ordered by UPGMA clustering.

**Figure 5.**
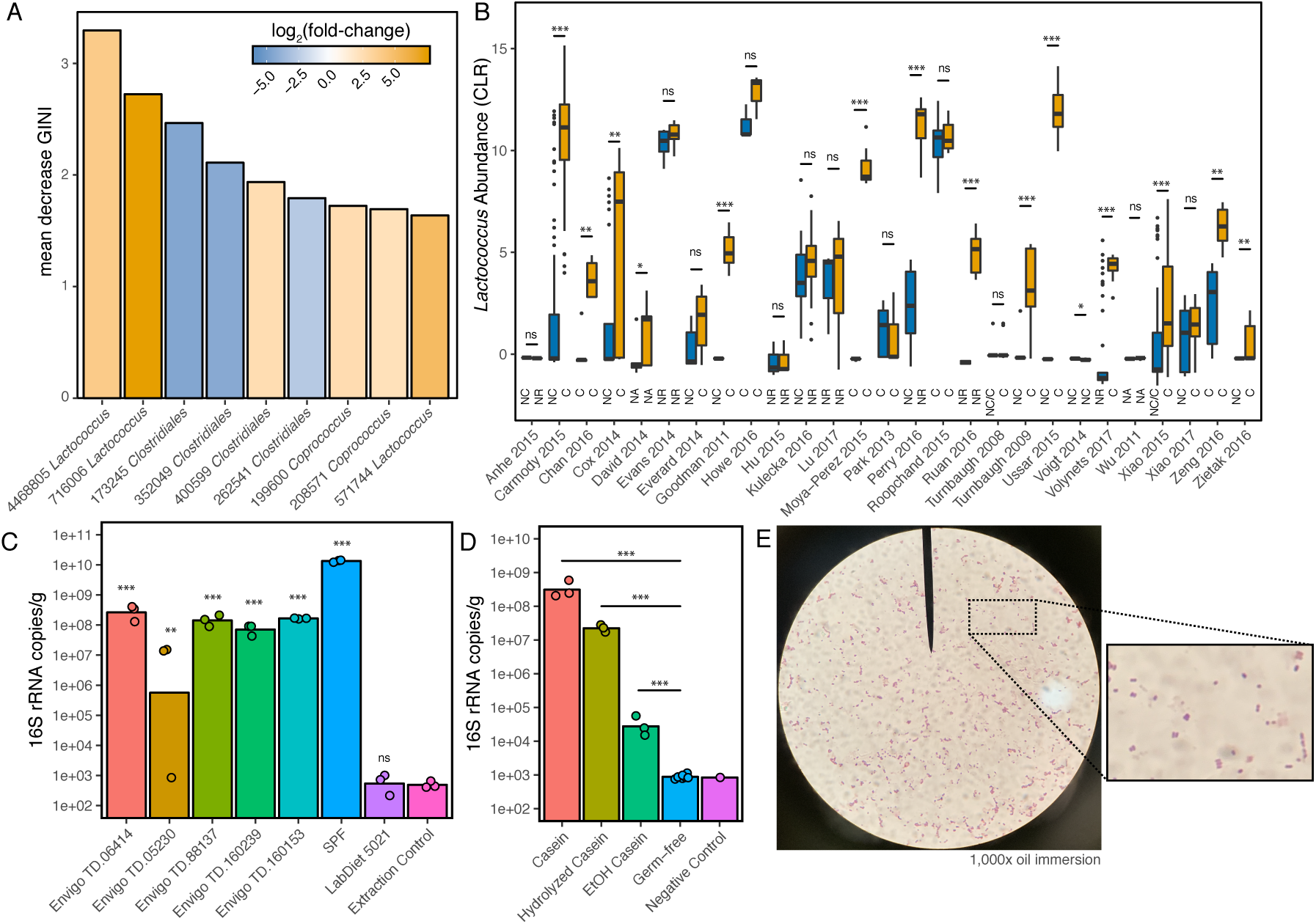
Widespread *Lactococcus* contamination in high-fat diets. **(A)** The most important features (as ranked by mean decrease in GINI coefficient) from a random forest classifier demonstrates an over-representation of *Lactococcus* spp. **(B)** Differential *Lactococcus* genus abundance on a per-study basis. Abundance is calculated as centered log_2_ ratio and diet composition is denoted as follows: C casein, NC no casein reported, NR diet content not explicitly reported or available, NA not applicable. **(C)** Fecal bacterial DNA content (expressed as 16S rRNA gene copies/gram wet weight) in culture-negative germ-free mice on various diets (n=1 mouse per diet analyzed in triplicate). **(D)** Bacterial DNA content in various casein preparations. *P*<*0.05, **P*<*0.01, ***P*<*0.001 Welch’s t-test (panel B), ANOVA with TukeyHSD (panels C and D). **(E)** Microscopy reveals Gram-positive cocci within stool sample of culture-negative germ-free mice fed a high-fat diet (TD.88137).

We began by feeding 6 semi-purified diets, and our standard chow diet, to 7 germ-free adult mice housed in individual Techniplast gnotobiotic isolators. In contrast to our standard chow controls (LabDiet 5021) which does not contain detectable levels of background DNA, the feces of mice fed all tested semi-purified diets contained detectable microbial DNA content orders of magnitude higher than negative controls (**Figure 5C**). To attempt to identify the nature of the *Lactococcus* signal, we sequenced two near-full length 16S amplicons from the semi-purified diets TD.88137 and TD.05230, which were found to match the most informative *L. lactis* OTU we identified (*>*99.2% nucleotide identity to OTU 716006, Genbank Accessions MK248688 and MK248689). Reasoning that casein may be the source of contamination due to the use of *Lactococcus* in dairy processes, we analyzed regular, hydrolyzed, and ethanol washed casein finding microbial DNA content in all 3 preparations. A 1,000-fold reduction in ethanol washed casein (**Figure 5D**) was observed, suggesting ethanol washed casein may be appropriate as the basis for formulating diets for future studies as it may be easily substituted in custom diet formulation. Attempts to culture bacteria using multiple rich medias under aerobic and anaerobic conditions, as well as M17 *Lactococcus* selective media, failed despite the presence of cell morphologies consistent with intact *Lactococcus* cells in the feces of culture-negative germ-free mice (**Figure 5E**, TD.88137). Together, these results suggest that high levels of intact but dead bacterial cells are present in multiple commonly used HFDs.

To prevent *Lactococcus* from creating a false signal of HFD in the entirety of our meta-analysis, we stripped all OTUs descendent from the most recent common ancestor of named *Lactococcus* OTUs in Green Genes 13-8 tree (64 OTUs). We then replicated all aspects of the analysis to this point observing that our findings have been robust to the removal of *Lactococcus* **(Extended Data)**.

The classifier was retrained as before finding that 228 OTUs, 108 KOs, and 456 Phylogenetic Nodes could predict the training set with a 9.7%, 8.1%, and 9.26% error rate respectively **(Table S5)**. These OTUs were heavily enriched in 3 major clades of *Lachnospiraceae, Ruminococcaceae* UCG-014, and *S24-7* (*Muribaculaceae*) OTUs within the Bacteroidetes (**Figure 4B**). Receiver operator curves were contrasted against a simple logistic regression model using the F/B ratio (**Figure 4C**). The calculated area under the curves (AUROCs) are shown in **Table S6** and on a per-study basis in **Figure 4D**. Likely due to data sparsity and interspecies variation in OTU content (**Figure** S5A-B), the model trained on OTUs was capable of predicting murine samples (AUROC*>*0.94), but failed to translate to humanized mice and humans (AUROC*<*0.64). By contrast, PhILR and KO-transformed data, which reduce dimensionality but preserve phylogenetic signals and functional information, provided considerably improved performance for both humanized gnotobiotic mice and humans (AUROC*>*0.75, **Table S6**). In all cases, these data outperformed the ratio of Firmicutes to Bacteroidetes demonstrating that this reproducible feature of murine HFD is not necessarily translatable across species. Humanized mice were not better predictors of human diet than conventional mice; however, this may be a function of data set size and limited diversity of studies as opposed to a reflection on underlying physiology **(Extended Data)**.

In considering the predictive performance of these models across studies (**Figure 4D**), it is apparent that these models are not heavily biased in favor of the prediction of only a particular subset of studies, but are generalizable across studies with the exception of the logistic model for F/B ratio whose predictive performance mirrors the data presented in **Figure 2D**.

## Discussion

We report the findings of a meta-analysis of murine-based microbiota-sequencing studies. Our results establish the effect of HFD on the gut microbiome in an effort to address reproducibility within the microbiome-diet field, and identify novel targets around which to build future experimental work. Although we only examined 25 studies in detail, it is notable that this group of studies has a wide range of diets, size, sequencing technologies employed, targets of sequencing, and rodent models examined. Overall, we demonstrated that a minimal set of OTUs reproducibly found across many laboratories can be used to successfully predict changes in microbial physiology in response to diet. This finding is notable in that this number of OTUs is relatively small and may be used to customize future experiments using gnotobiotic animals. Furthermore, we demonstrate that *Lactococcus* spp. produce a very strong and reproducible signal at the sequencing level across multiple studies and identified both computational and experimental strategies to address and explore this phenomenon further.

Interestingly, the F/B ratio is reproducibly increased following HFD feeding. This ratio was originally reported to be significantly increased in a rodent and human studies of obesity (Ley et al., 2005, 2006), but is not a reproducible marker across human cohorts examining BMI (Finucane et al., 2014; Sze and Schloss, 2016). It is important to point out that BMI and dietary feeding are two separate phenomena, and the implications of differing responses of the F/B ratio change in mice fed a HFD and obese humans is not clear. Recent work has indicated that this ratio is in part driven by the use of refined HFD relative to the more complex plant-polysaccharide-rich chow diets (Dalby et al., 2017). This particular signal either reflects a bloom of Firmicutes, an extinction of Bacteroidetes or some combination thereof which cannot be determined from sequencing-based approaches and whose mechanism is not fully appreciated. This could be direct, as a cause of nutrient consumption, or indirect as a result of diets’ effects on host-physiology.

Our findings also highly the considerable technical and experimental variation across studies. While many early projects employed shallow-sequencing depth and higher read length pyrosequencing across multiple 16S rRNA variable regions, most current studies use high-depth shorter-read amplicon or metagenomic sequencing. These differences may impact the taxonomic resolution of sequencing and associated error profiles with the potential to alter compositional profiles (Luo et al., 2012). There are also features by which studies vary that are not consistently reported (i.e. use of antibiotics or acidification in vivaria drinking water, use of specific type of nesting, number of animals housed per cage etc.), which likely make a significant impact on whether studies are reproducible. The lack of reporting of such features could simply be a lack of documentation or could be due to the fact that investigators may be separated from the care of animals within facilities and effectively blinded from these conditions. These collective variables contribute to what we term a study effect which has a major impact on microbial community composition **(Supplementary Table 3)** but whose individual effects are difficult to estimate due to potential issues of multicollinearity (**Figure** S1A).

Given multivariate evidence of consistent microbiota features of HFD-response, we defined a core set of features for follow up experimental studies capable of predicting diet both internally and externally. We were surprised that given the very high number of OTUs identified across our meta-analysis (n=29,937), merely 228 OTUs readily discriminated between HFD and LFD fed states (0.76%). This relatively small subset provides unique experimental opportunities which can be further prioritized based on their relative contributions to the models (**Table S5**). A major next step would be to isolate representatives of each OTU or to identify isolates (Lagkouvardos et al., 2016) that have similar functional profiles to colonize gnotobiotic animals and explore their impact on growth under high fat diet fed conditions in a gnotobiotic model.

Finally, we reproducibly detected a strong signal generated by *Lactococcus* spp. Future studies should account for this to prevent an artificial microbial signature of diet and consider avoiding formulations with a high *Lactococcus*-content, or ensuring both diets have consistent content. This is especially important as recent evidence suggests that non-viable *Lactococcus* cells can impact colonic inflammation (Ballal et al., 2015). Additional experiments are warranted to determine whether or not latent *Lactococcus* contamination in food is similarly able to alter host physiology.

There are several caveats worth noting. To render all datasets directly comparable, a closed reference OTU picking approach was applied which misses much of the resolution possible from denoised exact sequence variant-based approaches and may compress the true sample diversity (Callahan et al., 2017; Amir et al., 2017; Edgar, 2016). This strategy also restricts the number of possible observed OTUs to only those within the 13-8 version of the Green Genes database (DeSantis et al., 2006), and furthermore, taxonomic assignment within such datasets has been demonstrated to vary by length of sequencing product and variable region which inherently introduces and magnifies noise already present within these data sets (Edgar, 2017). A second problem worth noting is commonly referred to as the ‘file drawer problem’ wherein, published research is biased towards those studies with positive results (Gurevitch et al., 2018). This is likely why 22 out of our 25 murine studies demonstrate some effect with respect to influence of HFD on beta-diversity.

Despite clear inter-study variation, and issues in reproducing simple ecological metrics, the intention of this analysis was to examine the consistent features of the murine gut microbiota’s response to HFD to yield tangible targets for mechanistic studies. These results were generated from collective data across multiple laboratories and may be a robust foundation for future work geared on dissecting the links between shifts in microbial ecology, dietary intake, and the downstream consequences for host health and disease.

## Acknowledgments

We thank Robert Edgar for pivotal discussions and members of the Turnbaugh, Pollard, Spitzer, and Koliwad labs for their contributions of materials, critical feedback, and participation in abstract review. We also thank Katherine Pollard, Nirav Bhakta, Raman Khanna, and Jean Macklaim for critical feedback on the manuscript. This project was supported by the National Institutes of Health (R01HL122593; R21CA227232). PJT is a Chan Zuckerberg Biohub investigator and a Nadias Gift Foundation Innovator supported, in part, by the Damon Runyon Cancer Research Foundation (DRR-42-16) and the Searle Scholars Program (SSP-2016-1352). JEB was the recipient of a postdoctoral fellowship from Natural Science and Engineering Research Council of Canada.

## Author Contributions

Conceptualization, JEB, VU, and PJT; Methodology, JEB, JAT; Investigation, JEB, VU, KL, JAT, and PJT; Writing Original Draft, VU and JEB; Writing - Review and Editing, JEB, VU, and PJT; Funding Acquisition, PJT; Supervision, PJT.

## STAR METHODS

### CONTACT FOR REAGENT AND RESOURCE SHARING

Further information and requests for resources and reagents should be directed to and will be fulfilled by the Corresponding author, Peter Turnbaugh, or alternatively Jordan Bisanz.

## METHOD DETAILS

### Study Selection

The following all encompassing search term was entered into PubMed and the NCBI Sequence Read Archive (SRA) in July 2017 to generate an unbiased representation of studies studying the effect of diet composition on the murine gut microbiome: “high fat diet”[All Fields]AND “microbiome”[All Fields] OR “high fat diet”[All Fields] AND “microbiota”[All Fields] OR “diet induced obesity”[All Fields] AND “microbiome”[All Fields] OR “diet induced obesity”[All Fields] AND “microbiota”[All Fields] OR “ketogenic diet”[All Fields] AND “microbiome”[All Fields] OR “ketogenic diet”[All Fields] AND “microbiota”[All Fields] OR “western diet”[All Fields] AND “microbiome”[All Fields] OR “western diet”[All Fields] AND “microbiota”[All Fields] OR (high-fat[All Fields] AND high-sugar[All Fields] AND (”diet”[MeSH Terms] OR “diet”[All Fields])) AND “microbiome”[All Fields] OR (high-fat[All Fields] AND high-sugar[All Fields] AND (”diet”[MeSH Terms] OR “diet”[All Fields])) AND “microbiota”[All Fields] OR ((”obesity”[MeSH Terms] OR “obesity”[All Fields]) AND promoting[All Fields] AND conditions[All Fields]) AND (”microbiota”[MeSH Terms] OR “microbiota”[All Fields]) OR ((”obesity”[MeSH Terms] OR “obesity”[All Fields]) AND promoting[All Fields] AND conditions[All Fields]) AND (”microbiota”[MeSH Terms] OR “microbiota”[All Fields] OR “microbiome”[All Fields]) OR ((microbiome[All Fields] OR microbiota[All Fields] OR microflora[All Fields] OR “microbial ecology”[All Fields]) AND (mouse[All Fields] OR murine[All Fields] OR “mus musculus”[All Fields]) AND (diet[All Fields]) AND (sugar[All Fields] OR high-sugar[All Fields] OR low-sugar[All Fields] OR fat[All Fields] OR high-fat[All Fields] OR low-fat[All Fields] OR ketogenic[All Fields]) AND (sequenc*[All Fields] OR 16S[All Fields] OR metagenom*[All Fields]) AND (fecal[All Fields] OR feces[All Fields] OR digest*[All Fields] OR gut[All Fields] OR intestin*[All Fields])).

The search yielded 427 potential studies for inclusion. To filter these studies on relevance, we employed a crowdsourcing approach wherein the studies were randomly distributed to 11 laboratory-based volunteers via a purpose-built web application AbstractReviewR (source code and reviewer instructions available: https://www.github.com/jbisanz/AbstractReviewR). Reviewers were provided the title, abstract, year, journal, and author list (as automatically retrieved from NCBI). Each study was reviewed by 2 adjudicators who were blinded to each others decisions. One author (VU) reviewed all abstracts and this vote was used to resolve a split decision. Consensus conclusions were accepted with the exception of 5 studies that were included post-review due to clear relevance and available metadata and sequencing data.

Studies that fulfilled criteria for the meta-analysis were then evaluated for sample type. In the event that studies varied another variable besides diet (i.e. genetic manipulation, use of a probiotic, addition of a non-dietary based supplement etc.), samples were selected where dietary fat content was the principal variable modulated and controlled for, and all other samples were discarded. In the case of cross-sectional longitudinal sample collection, only end point samples were analyzed. In the case of interventional time-course, only baseline and endpoint samples were analyzed. Categorization of HFD or LFD was done on a per study basis; some studies had three diets that were studied, and the diet highest in fat or animal fat content was assigned the designation of HFD.

### Data Retrieval and OTU picking

Code to recapitulate analysis is available at https://www.github.com/jbisanz/MetaDiet and in **Extended Data**. Sequence data was downloaded directly from the SRA and MGRAST by listed accessions **(Table S2)**. When only multiplexed runs were available, they were demultiplexed using either usearch-fastx demux or custom code in R using the shortRead package. Other studies were retrieved from their respective lab repositories (https://gordonlab.wustl.edu/TurnbaughSE_10_09/STM_2009.html) but have been redeposited to the Sequence Read Archive under BioProject PRJNA482456.

Where paired reads were available, reads were overlapped using vsearch 2.4.4 (Rognes et al., 2016). If *>*50% of reads could be overlapped, the merger was carried forward for analysis; otherwise, only the forward reads were considered. Reads were filtered on quality (where available) again using vsearch with the following parameters –fastq trunqqual 20, –fastq maxns 0, –fastq minlen 60, –fastq maxee 2. Next reads were prefiltered using SortMeRNA (Kopylova et al., 2012) using the SILVA-Bac-16S-id90 database. Finally, OTUs were picked against the 13-8 Green Genes release clustered at 97% identity using usearch 10.0.240 (Edgar, 2010)-closed ref with the following parameters:-strand both, -id 0.97. The resulting table was converted to a biom file for processing with PICRUSt (Langille et al., 2013) with the following scripts: normalize by copy number.py and predict metagenomes.py.

### Diversity analysis

For analysis on a per study basis, samples were rarefied (Subsample. Table, MicrobeR 0.31[https://www.github.com/jbisanz/MicrobeR]) to the lowest depth sample within the study for generating alpha diversity metrics. The diversity and estimateR functions of Vegan (Dixon, 2003) were used to generate Shannon’s diversity index (log base e) and Chao1 estimates respectively and Picante (Kembel et al., 2010) was used to generate Faith’s phylogenetic distance. To generate the Firmicutes to Bacteroidetes ratio, the OTU table was summarized to phylum level (Summarize.Taxa, MicrobeR) and the log_2_ of the ratio of proportional abundances was calculated with a prior count of 0.1%. UniFrac and Jensen-Shannon divergence were calculated using the parallel-enabled distance function of Phyloseq (McMurdie and Holmes, 2013) on subsampled proportional abundances. Bray-Curtis dissimilarity was also calculated (vegdist, Vegan) on subsampled proportional abundances. The CLR Euclidean distance was calculated by carrying out a centered log2-ratio transformation (Make.CLR, MicrobeR) with count zero multiplicative replacement (zCompositions, (Martín-Fernández et al., 2014)) followed by calculating the Euclidean distance (dist, base R 3.5.0). The PhILR Euclidian distance was calculated by first carrying out the phylogenetic isometric log ratio transformation (philr, PhILR, (Silverman et al., 2017)) and calculating the distance matrix as before. Principal coordinates analysis was carried out using the pcoa function of APE (Paradis et al., 2004). ADONIS calculations were carried out (adonis, Vegan) with 999 replications on each distance/dissimilarity metric. All studies were internally normalized against the geometric mean of the LFD group and statistical analysis was determined using Welch’s t-test (t.test, base MRO 3.4.3) to determine significance and the 95% confidence interval. Anhe 2015 was not included in Figure 2 due to n=2 per group. The combined analysis was conducted using a linear mixed effects model (Kuznetsova et al., 2017) with the formula *log*_2_(*difference*)∼*Diet* + (1| *Study*), significance was determined with Satterthwaite’s method using the anova function and 95% confidence interval using the confint function.

### Random Forest Classifiers

3 murine studies were randomly sampled to be used as validation studies and the remaining samples were randomly separated into a training set and validation set (2/3 and 1/3 respectively). CLR normalized abundances, PhILR abundances, and CLR normalized PICRUSt abundances were used as predictor variables for the HFD/LFD cases. 10-fold cross validation (rfcv, RandomForest, (Breiman, 2001)) was carried out to determine the optimal number of features for classifier accuracy. The number of predictor variables was determined by selecting the point of saturation in minimizing error rate and selecting the features based on ranked MeanDecreaseGINI, all other features were excluded from the model. *M*_*try*_ and *N*_*tree*_ were left as default values 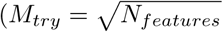 and *N*_M*tree*_ = 500). The final model was trained (randomForest, Random-Forest) and then applied to all data sets. Predictions and performance metrics were generated using the predict, prediction, and performance functions of ROCR. The entirety of the analysis was repeated after removing *Lactococcus* spp., by finding the most recent common ancestor in the reference tree to all named *Lactococcus* spp. (getMRCA, APE), finding its constituent tips (extract.clade, APE) and removing these OTUs from the OTU table and tree (drop.tip, APE).

### *Lactococcus* content of diets and casein

To examine the effect of semipurified diets on fecal microbial DNA content, C57BL/6J and BALB/c mice were singly housed in Techniplast rack-mounted individual isolators. Fecal contents were collected after approximately 1 week. These experiments were conducted under protocol AN170098 approved by the UCSF Institutional Animal Care and Use Committee. Animals were fed the following diets: LabDiet 5021 and Envigo TD.05230, TD.88137, TD.160239, and TD.160153, and TD.064114. To examine microbial DNA content of casein, we were able to obtain complementary samples from Envigo of the following varieties: standard casein (cat num. 032.0024, lot 18322), ethanol washed vitamin-free casein (0.32.0352 lot 2458429), and hydrolyzed casein (0.33.2599 lot 7333). To confirm the lack of viable cultures, feces was used to inoculate LB, YPD, and sheep’s blood agar (aerobically) and BHI and sheep’s blood agar (anaerobically, COY anaerobic system) and incubated for 48h at 37C. Selective culture of *Lactococcus* was carried out using MRS and M17 media (Oxoid) supplemented with 1% lactose and incubation at 30C aerobically for 48h. DNA was extracted from fecal pellets and casein using the Zymbiomics 96 MagBead DNA kit following the manufacturers instructions with an additional 10 minute incubation at 65C after cell disruption. Quantification against a standard curve of purified gDNA from *Eggerthella lenta* was performed by qPCR of the V6 region using BioRad iTaq Universal Probes Supermix in 10 *µ*L reactions using 200 nM of the following oligonucleotides: 5’-TGGAGCATGTGGTTTAATTCGA-3’, 5’-TGCGGGACTTAACCCAACA-3’, 5’-[Cy5]CACGAGCTGACGACARCCATGCA[BHQ3]-3’. Reactions were conducted in a BioRad CFX384 with the following cycle parameters: 95C 5min followed by 40 cycles of 95C for 5 sec, 60C for 15 sec.

## QUANTIFICATION AND STATISTICAL ANALYSIS

Unless otherwise specified, statistical analysis was carried out in R 3.5.0 using the appropriate base function. Individual data points have been shown where possible but are otherwise represented as the mean*±*standard error. Significance was determined as P*<*0.05 unless otherwise stated. Randomization was carried out using the sample function of R 3.5.0 with a fixed seed for reproducible sampling as identified in **Extended Data**.

## DATA AND SOFTWARE AVAILABILITY

All datasets analyzed in this study are available from public sources as identified in **Table S2** and the STAR Key Resources Table. Precomputed feature tables are available for download in the github repository available at https://www.github.com/jbisanz/MetaDiet.

## KEY RESOURCES TABLE

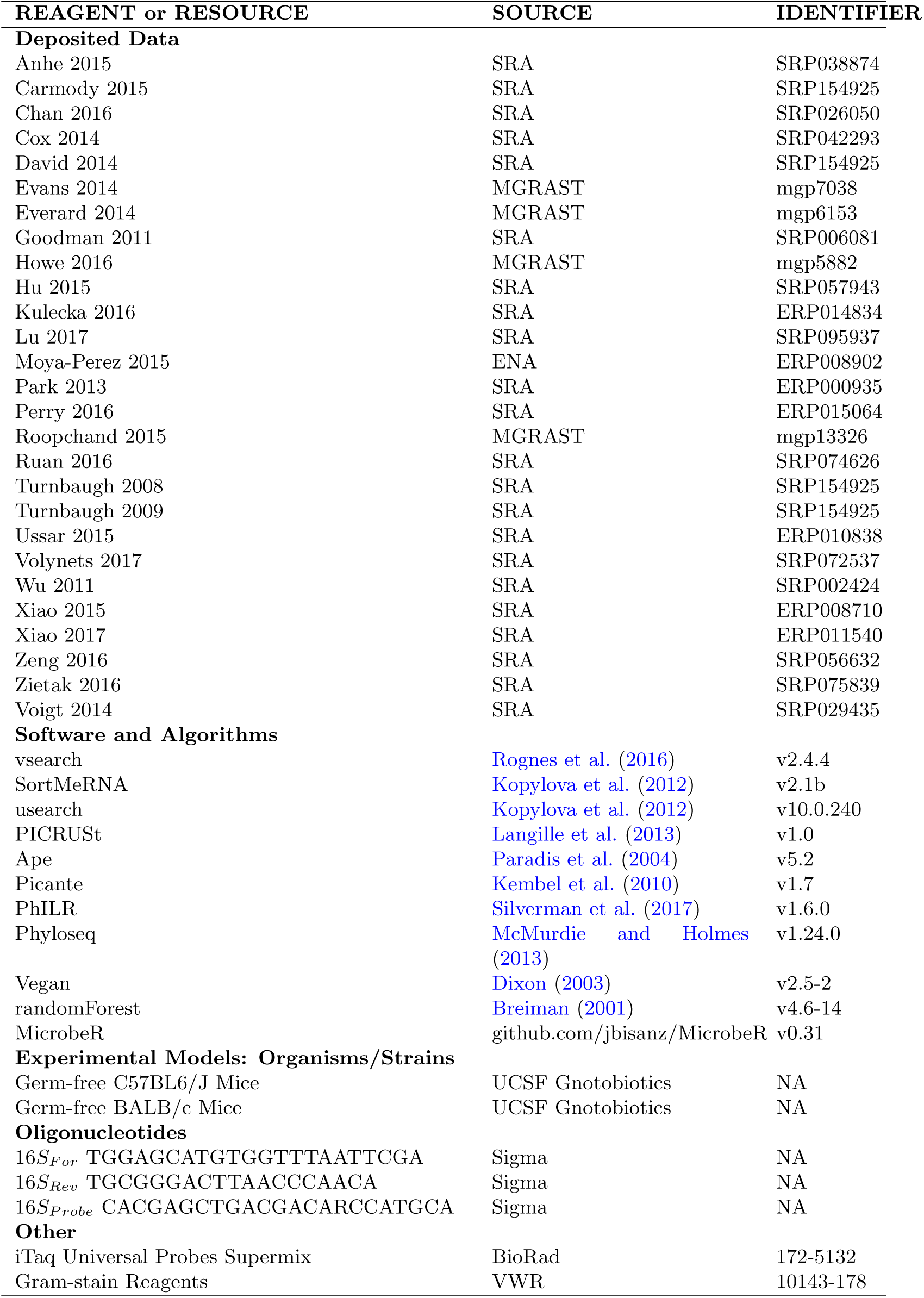

## Supplemental Figures

**Figure S1. relating to Figure 1.**
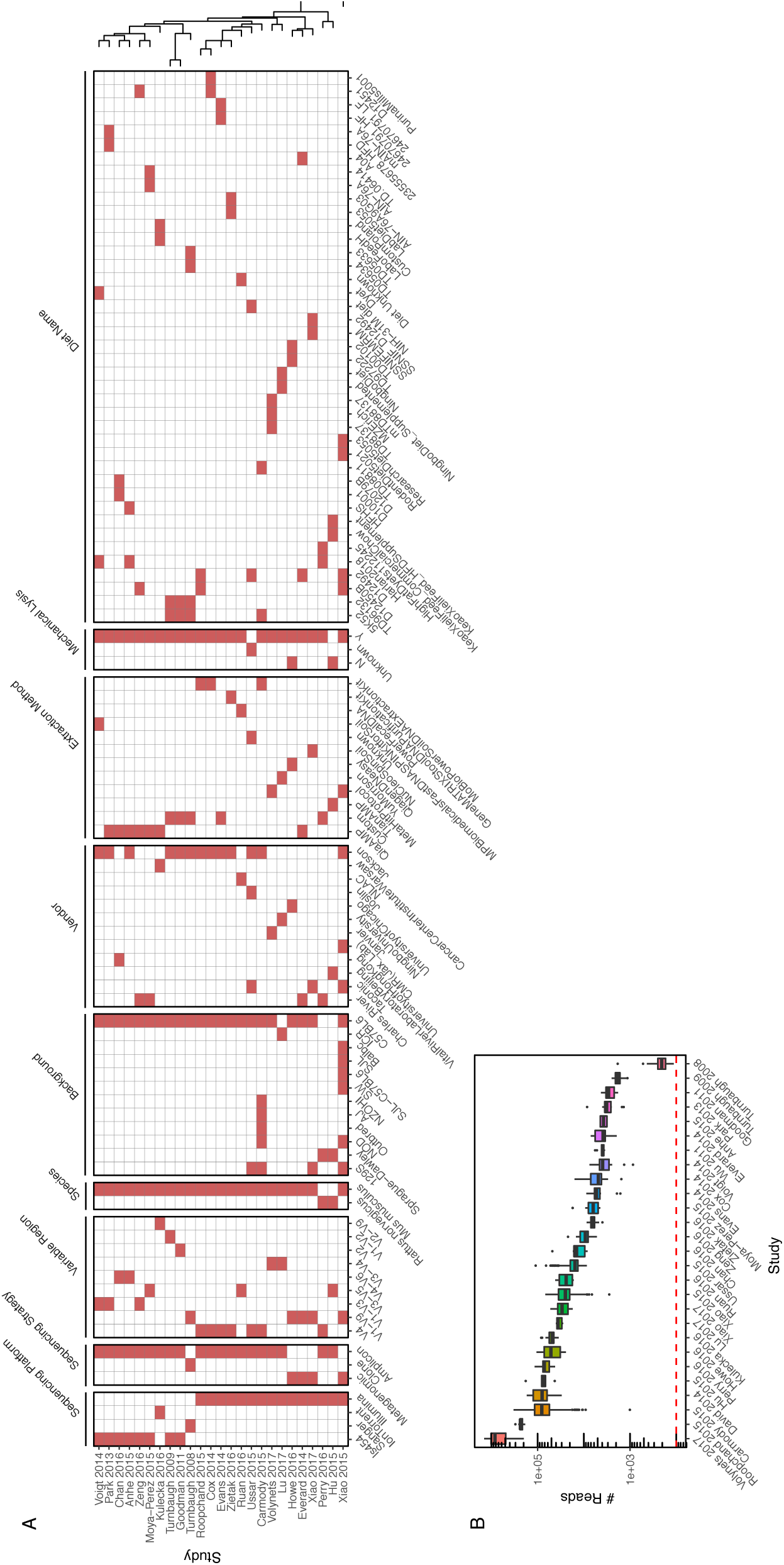
**(A)** Study metadata with studies ordered by UPGMA clustering. **(B)** Read depth distribution among studies. The dashed red line denotes the subsampling depth for pooled analysis.

**Figure S2.**
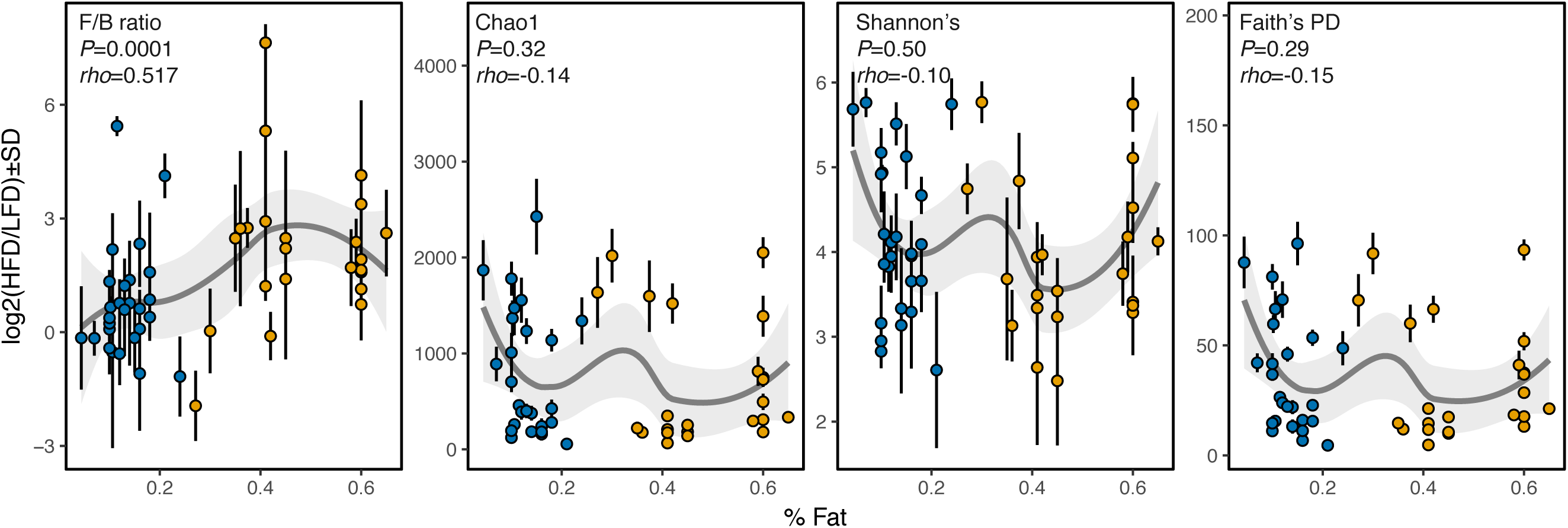
Correlations of diversity and dietary fat content relating to Figure 2. Each point represents a study’s HFD or LFD group (as colored) with the x axis representing the fat content. The trend line represents a LOESS regression *±*standard error. Dietary fat is positively correlated with the ratio of Firmicutes to Bacteroidetes (P=0.0001, *rho*=0.517).

**Figure S3.**
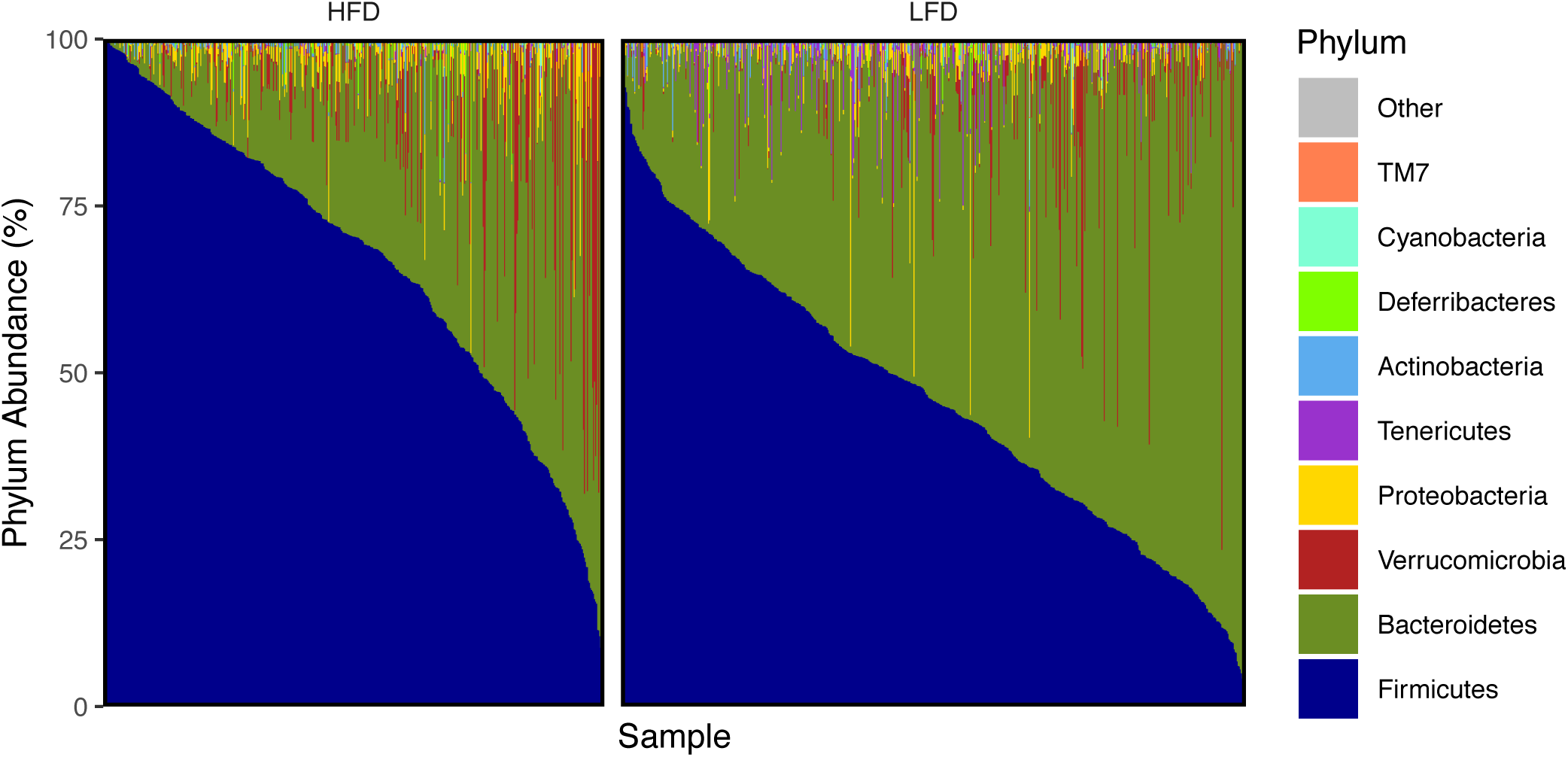
Phylum-level bar plot contrasting HFD and LFD across all studies relating to Figure 2.

**Figure S4.**
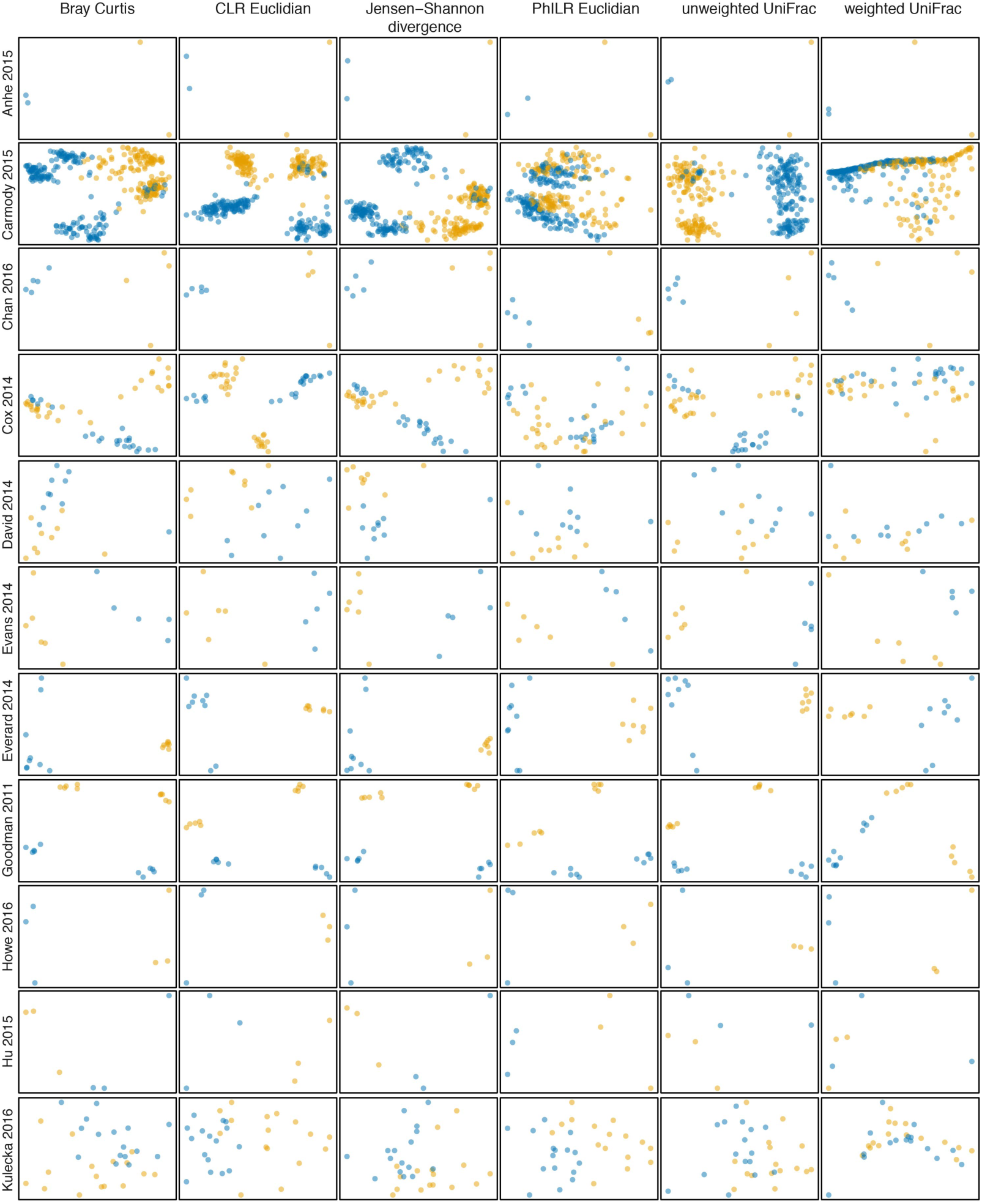
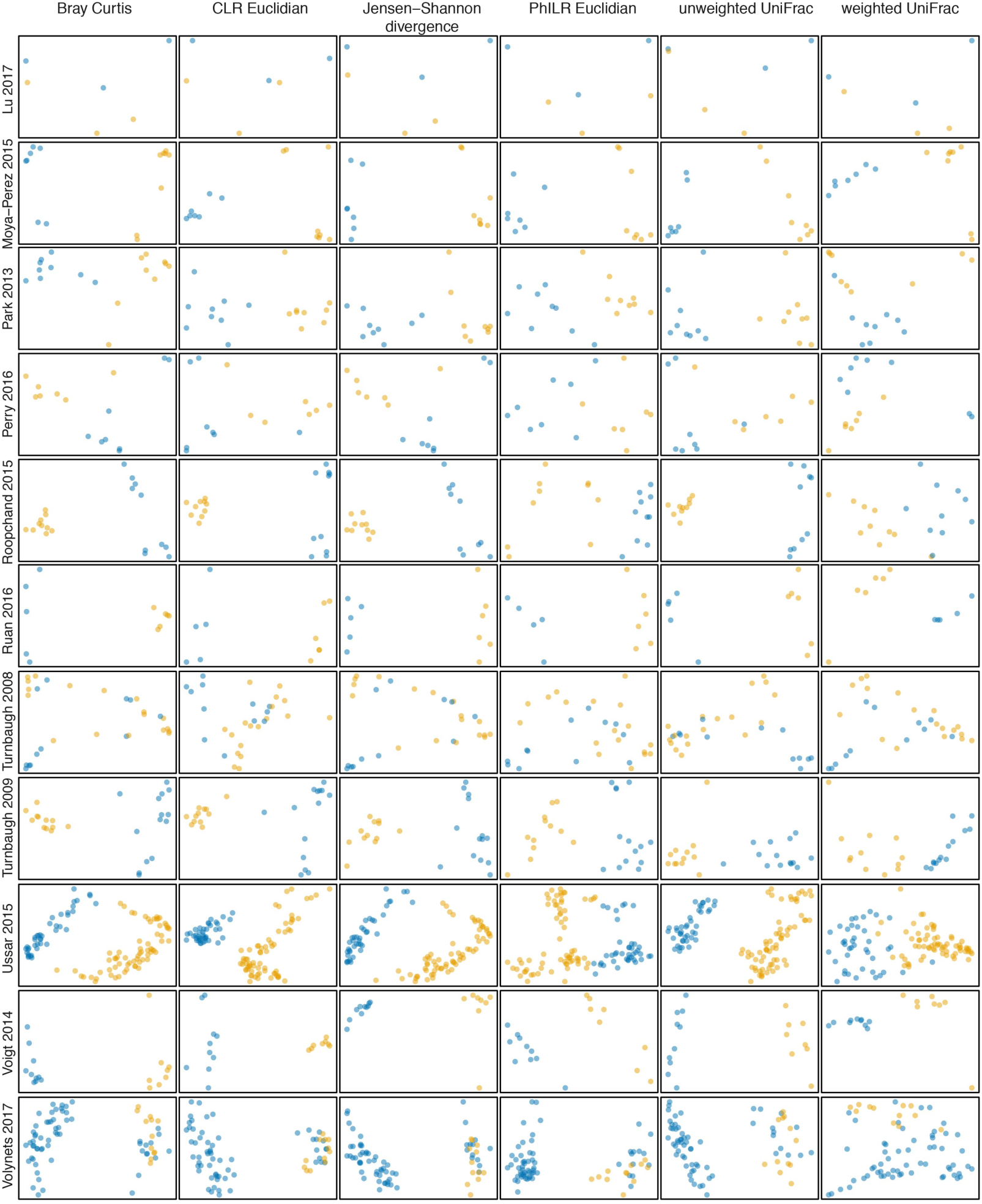
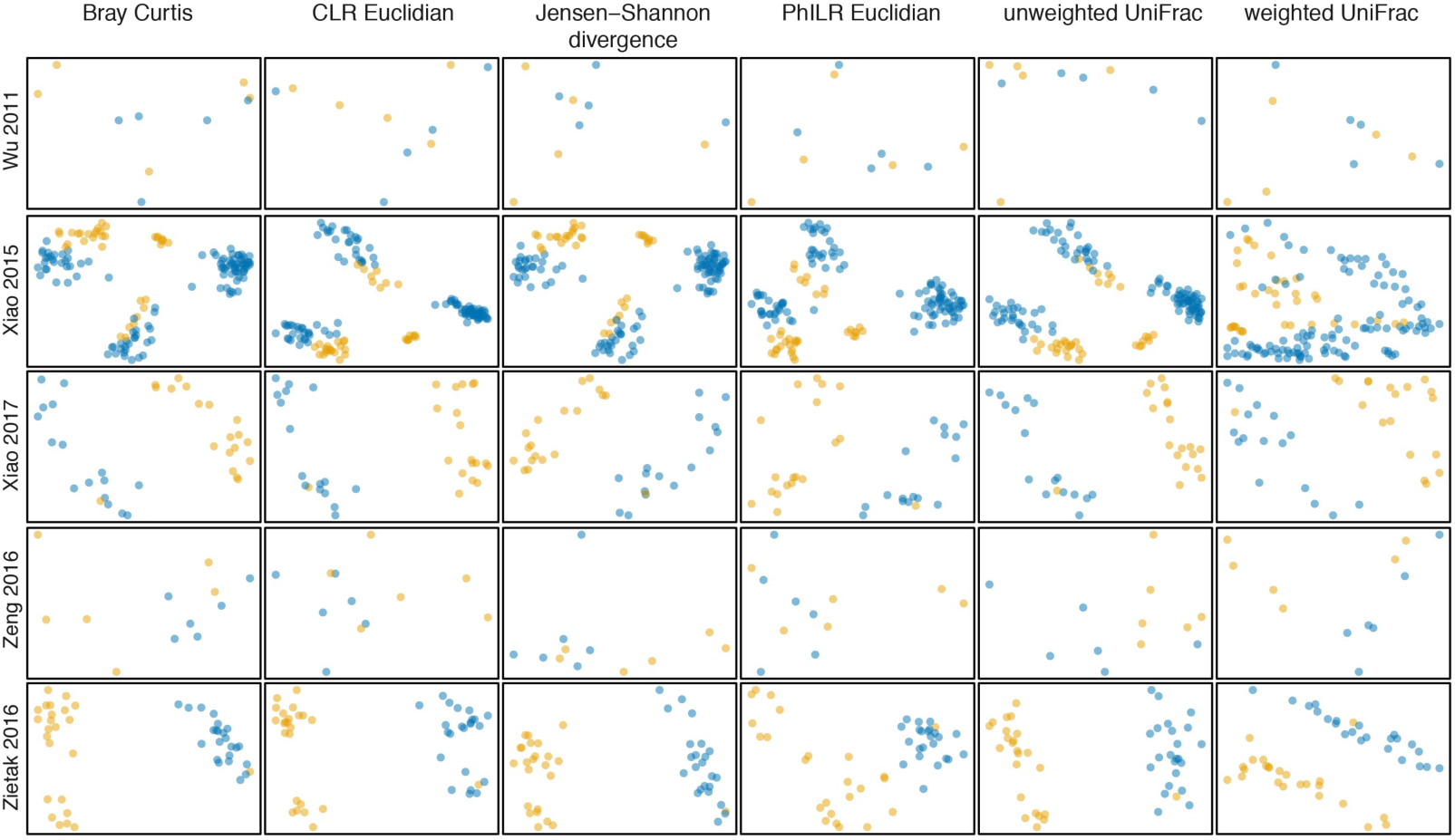
Principal coordinate analysis on a per-study basis relating to Figure 3. X and Y axes represent PC1 and PC2 respectively.

**Figure S5.**
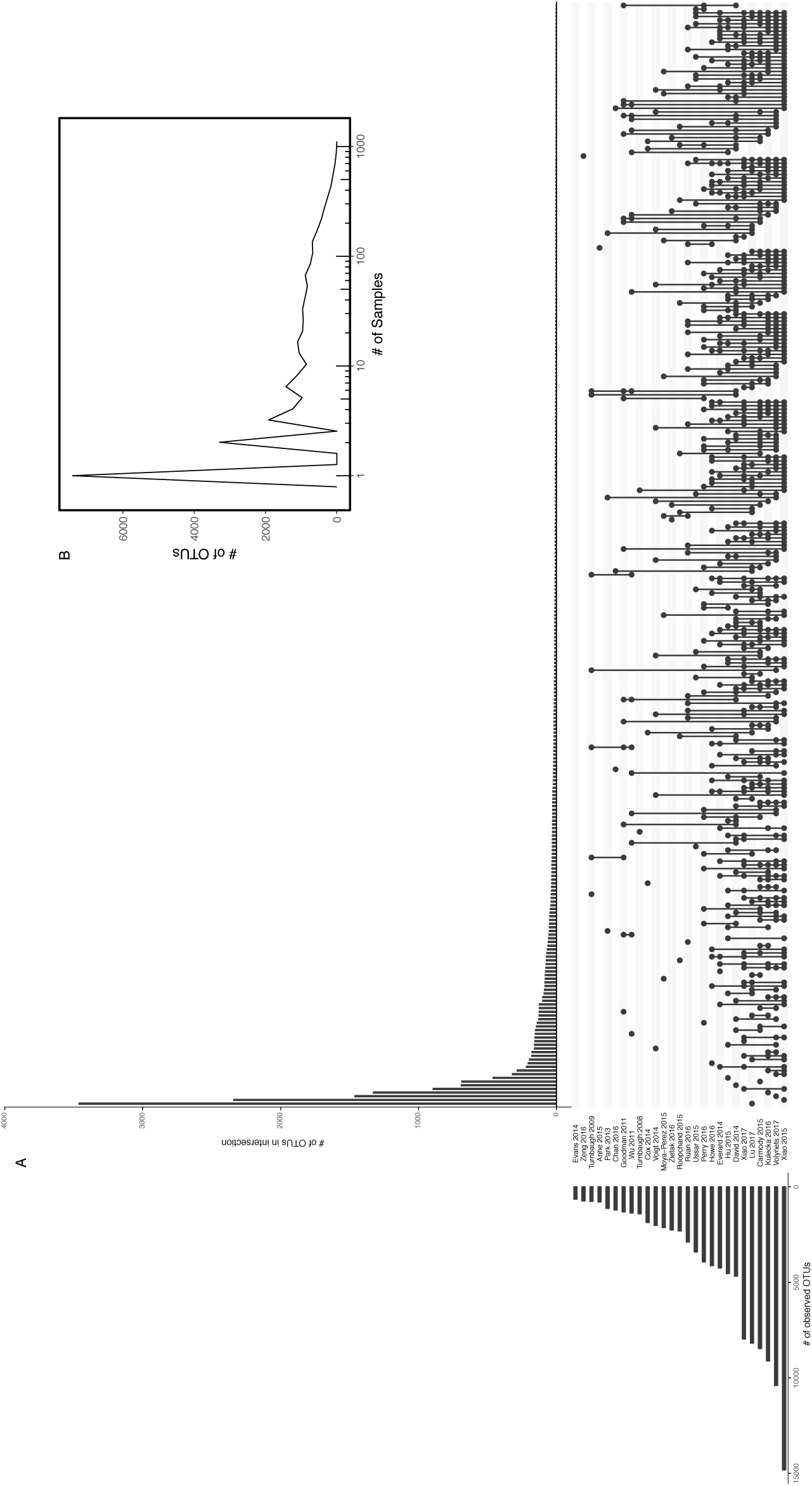
OTU content relating to Figure 3. **(A)** Upset diagram showing overlap in OTU content between studies. **(B)** Distribution of OTUs across samples contributing to OTU table sparsity.

**Figure S6.**
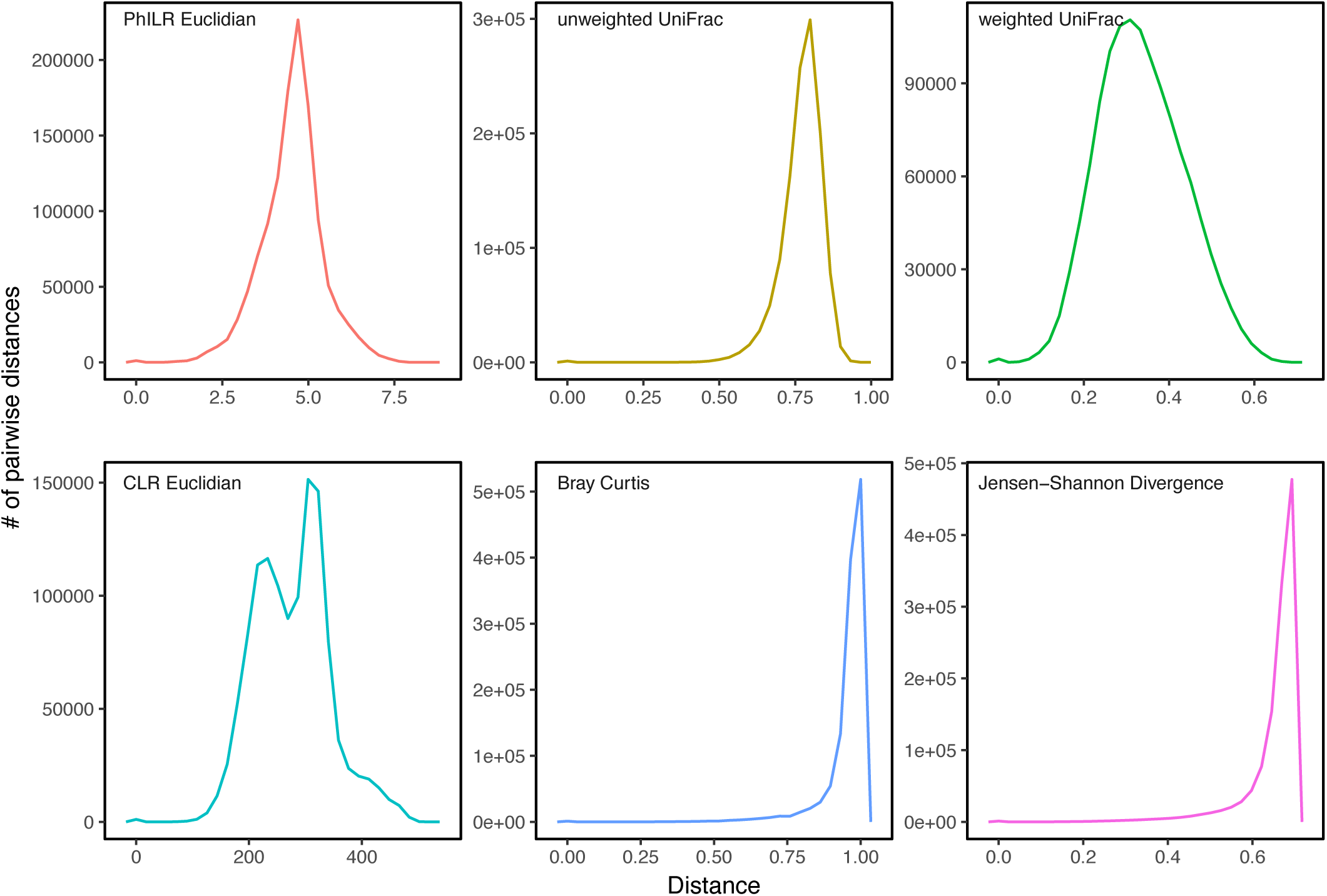
Distribution of intersample distances Figure 3. A high degree of saturation is noted in Bray Curtis (non-overlapping=1) and Jensen-Shannon Divergence (non-overlapping=ln(2)=0.693). CLR euclidean was pre-emptively removed due to possible effects of zero-replacement for logarithmic normalization.

**Figure S7.**
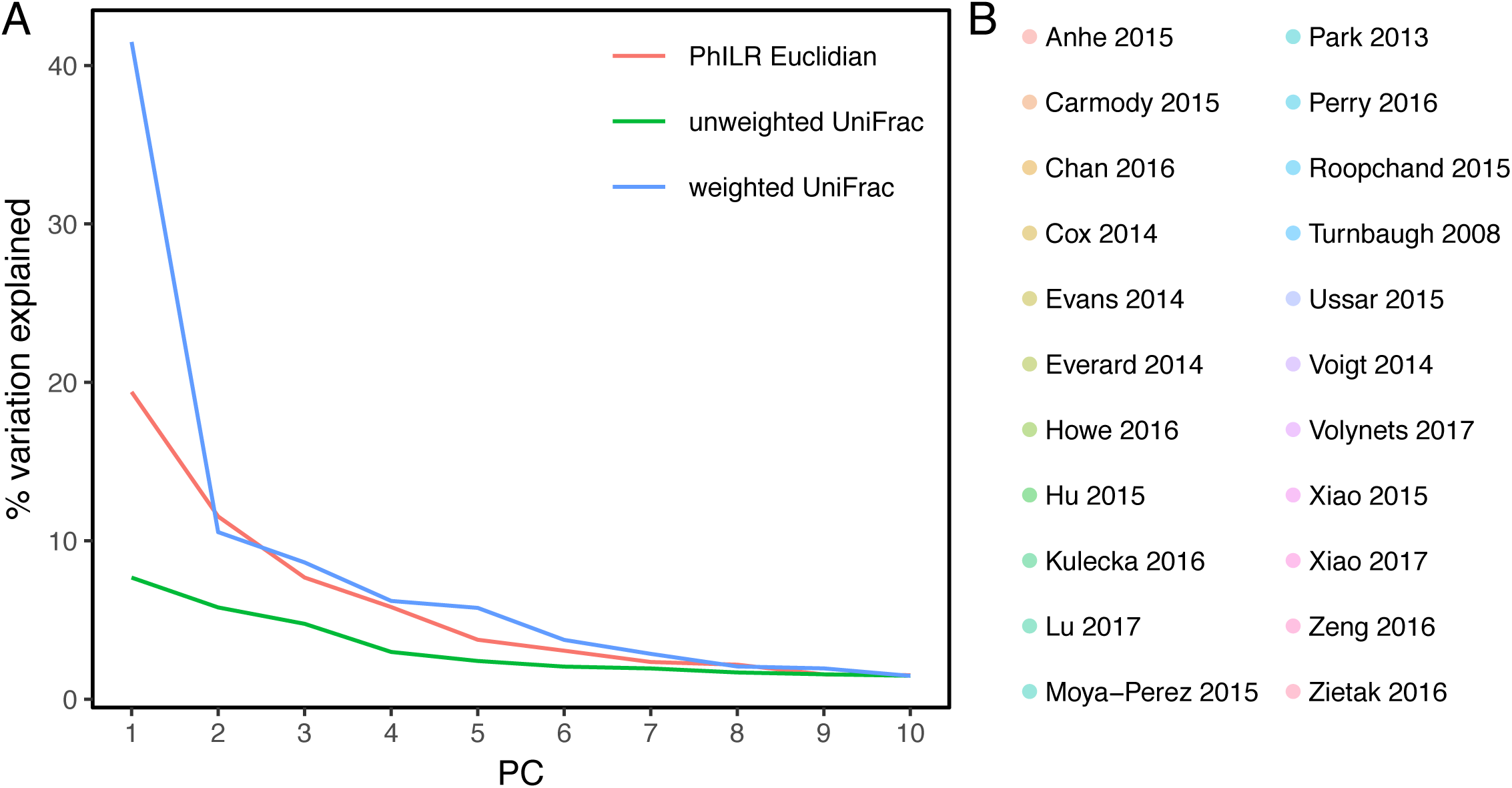
Relating to Figure 3. **(A)** Scree plot showing variation explained by axis for PCoAs in **Figure 3**. **(B)** Color legend for **Figure 3**.

## Supplemental Table Legends

**Supplemental Table 1 relating to Figure 1.**
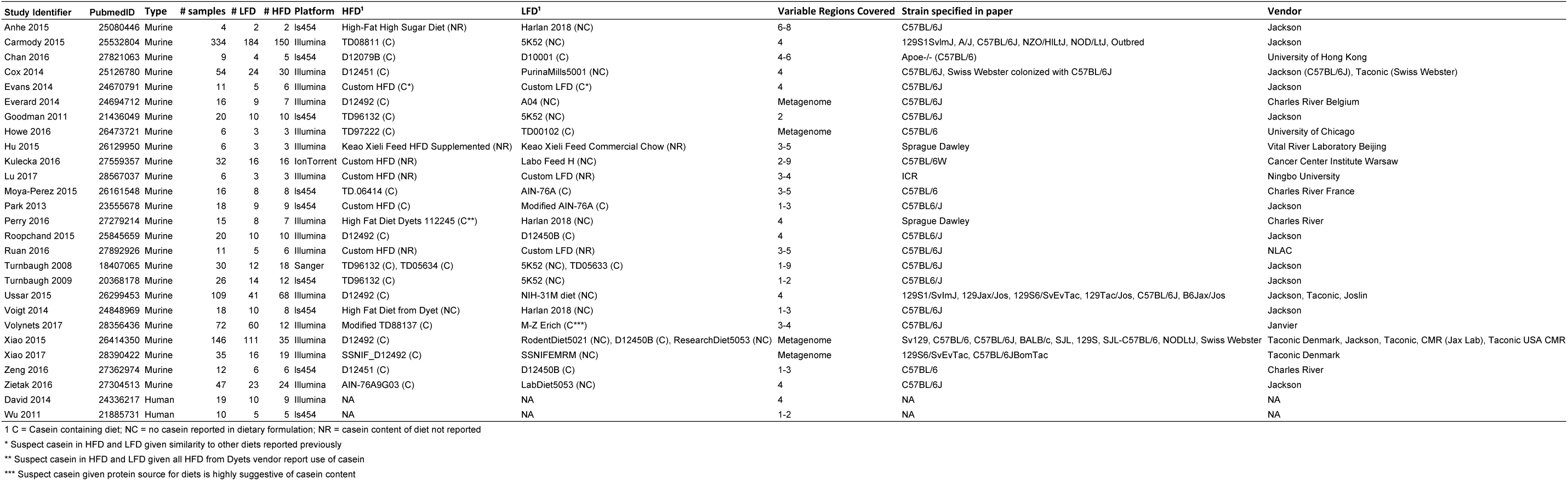
Descriptions of studies meeting inclusion criteria.

**Supplemental Table 2 relating to Figure 1.**
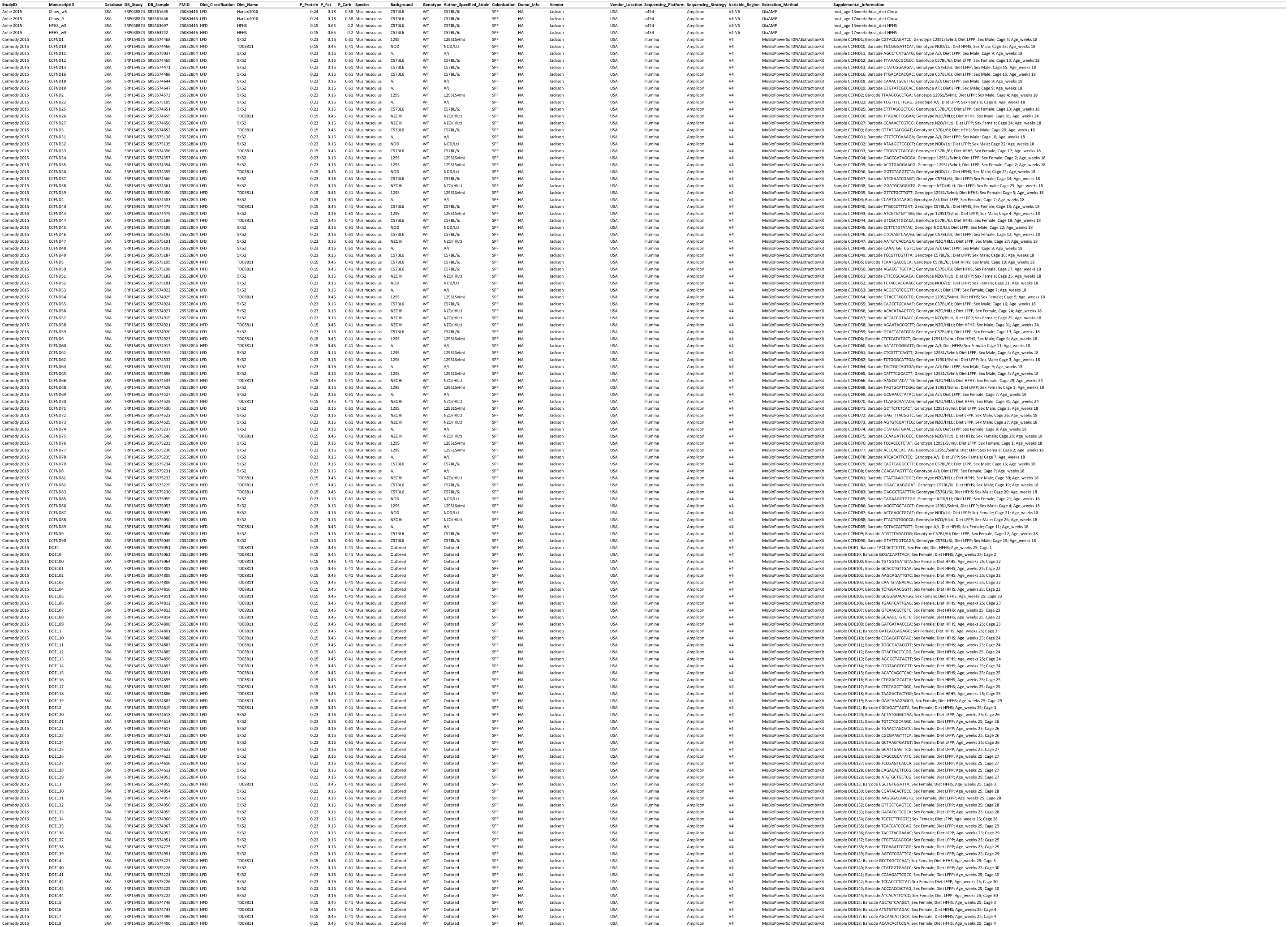
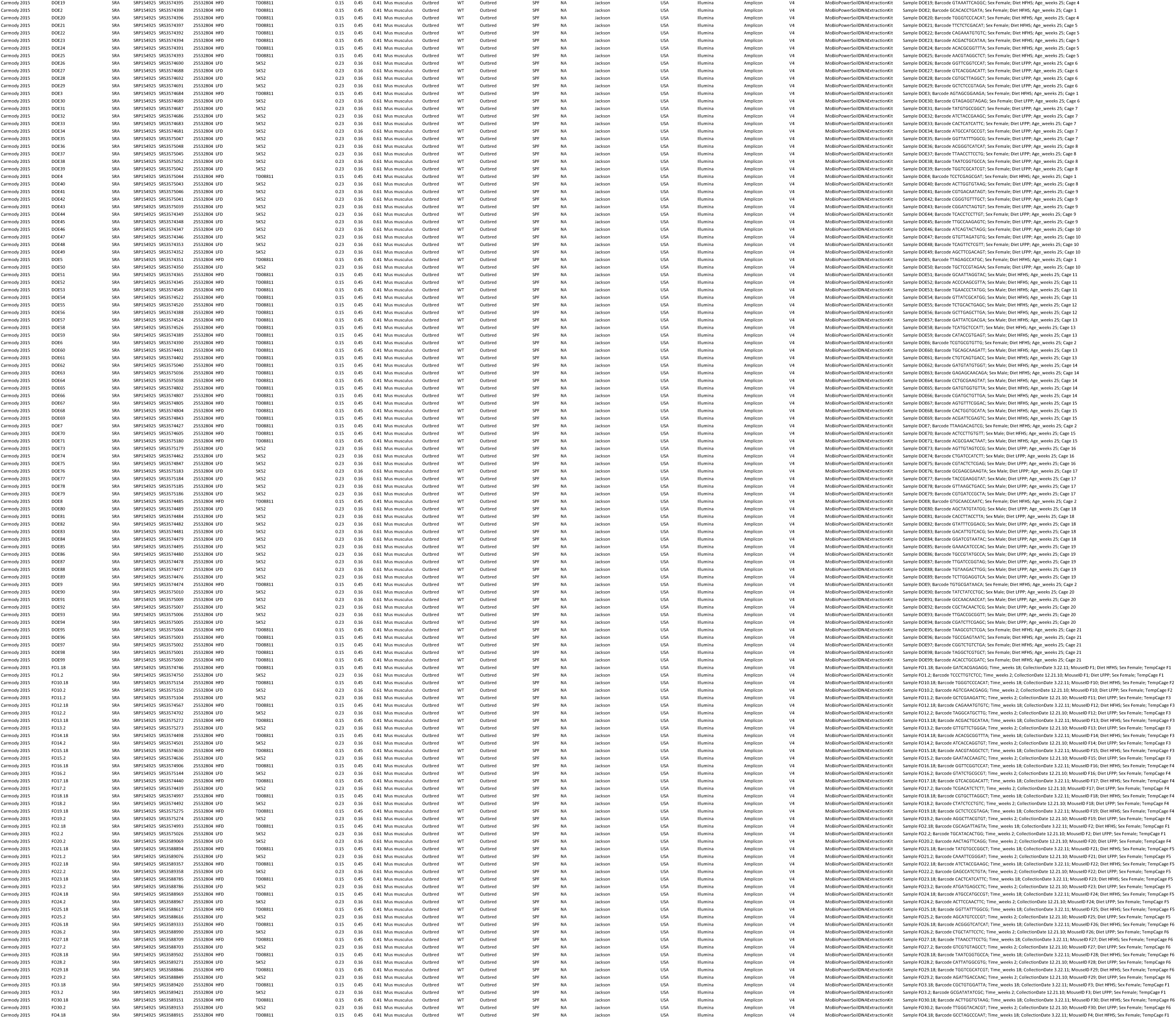
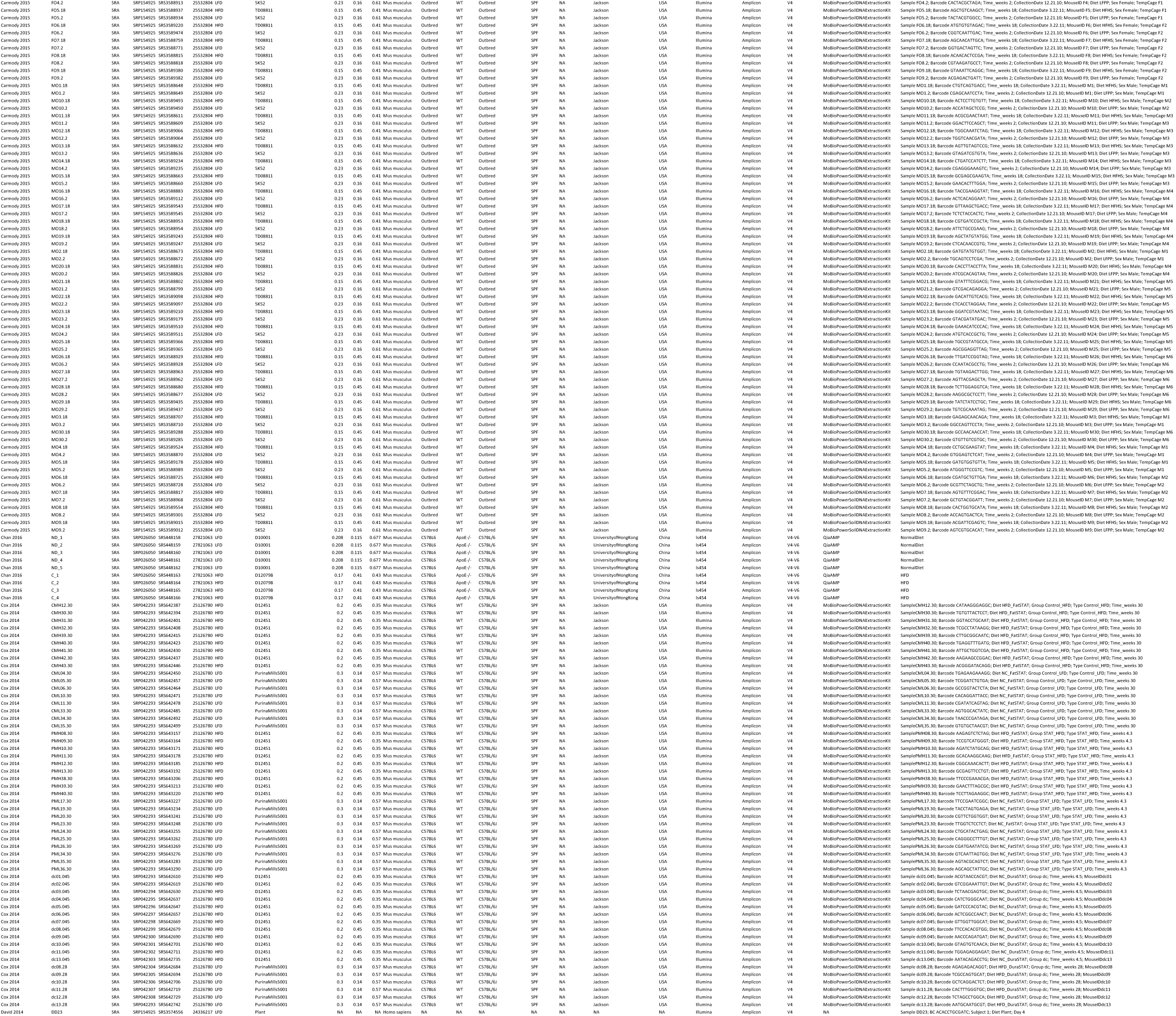
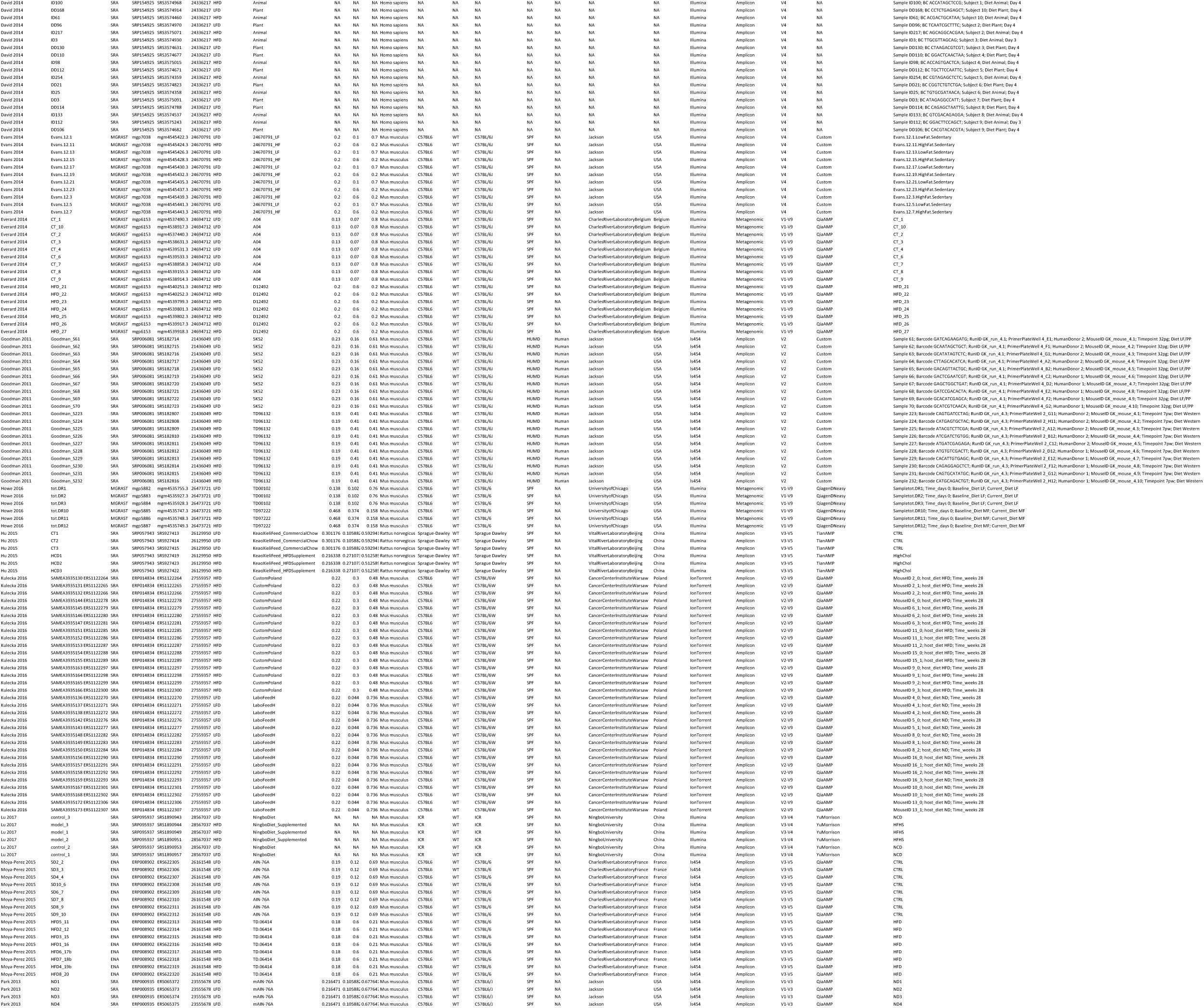
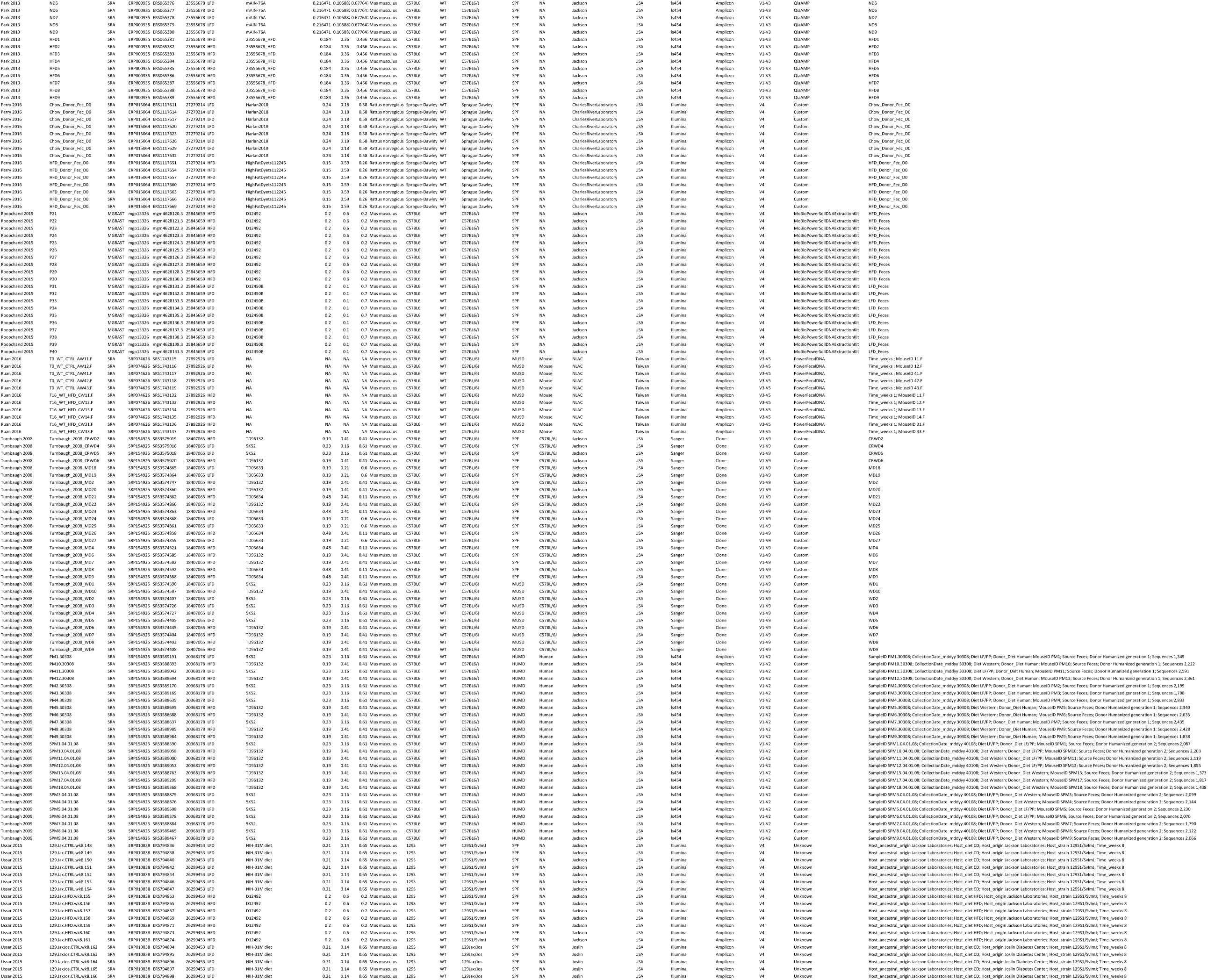
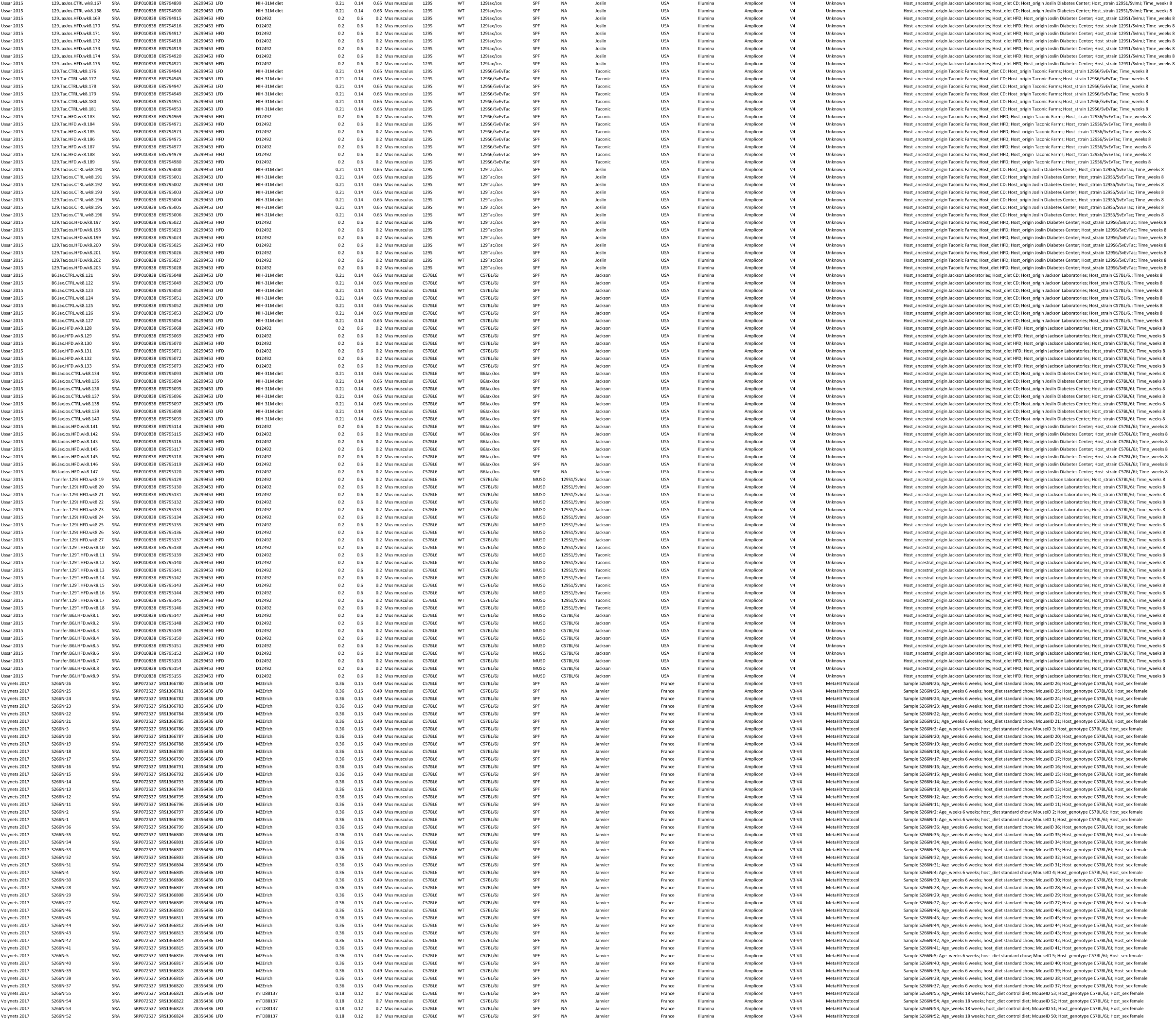
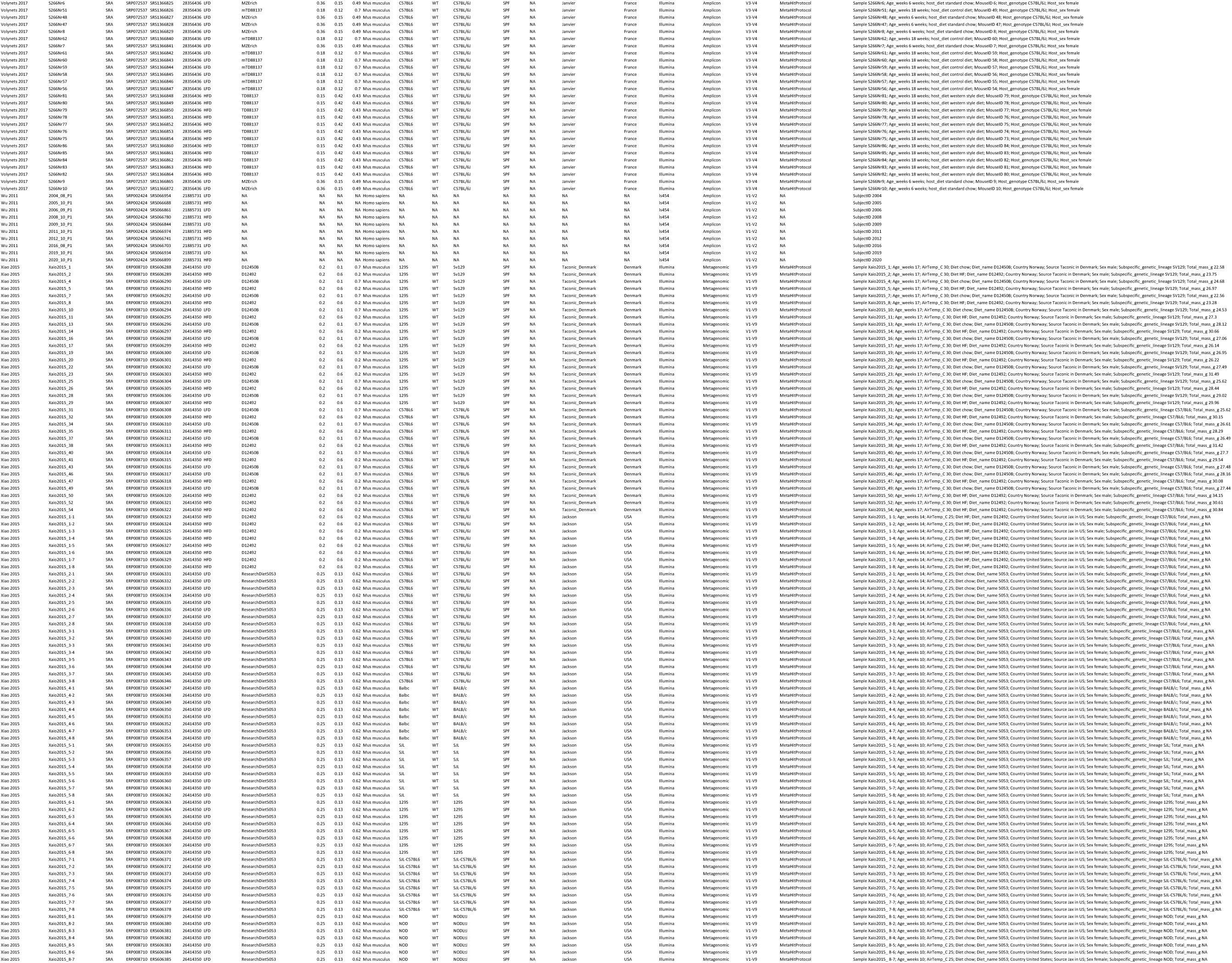
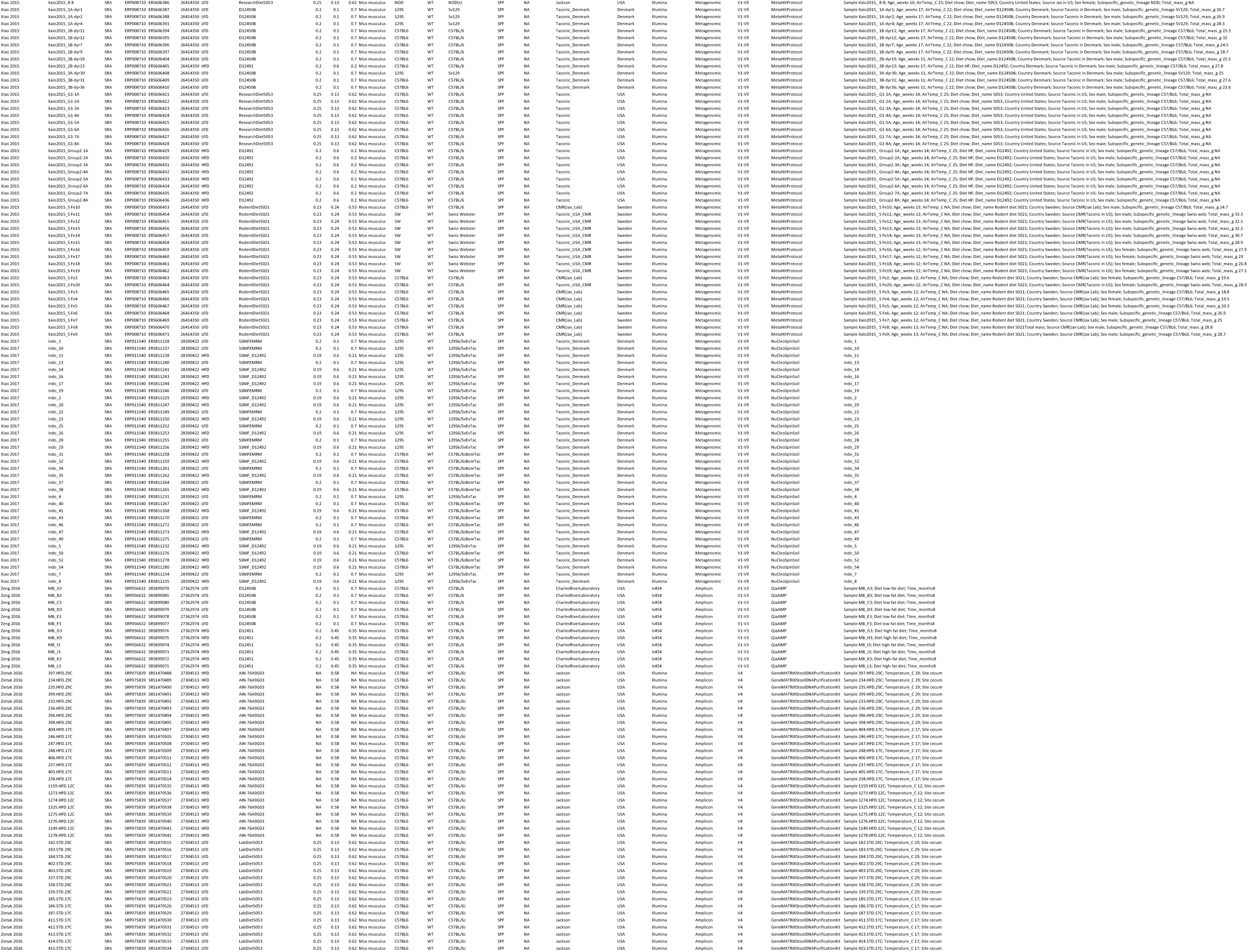
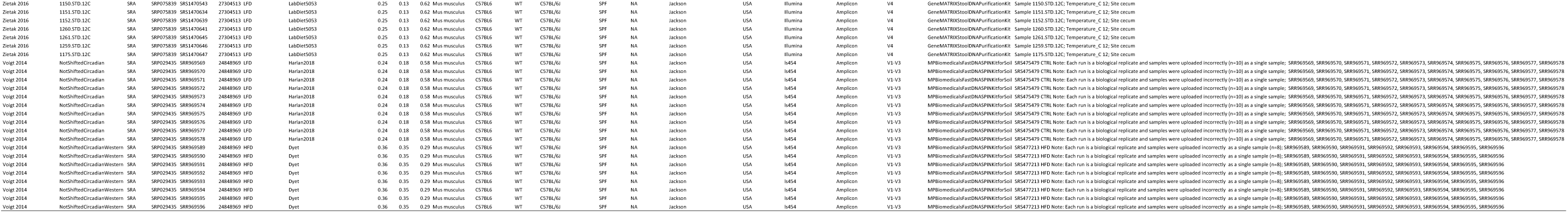
Per-sample metadata.

**Supplemental Table 3 relating to Figure 2 and 3.**
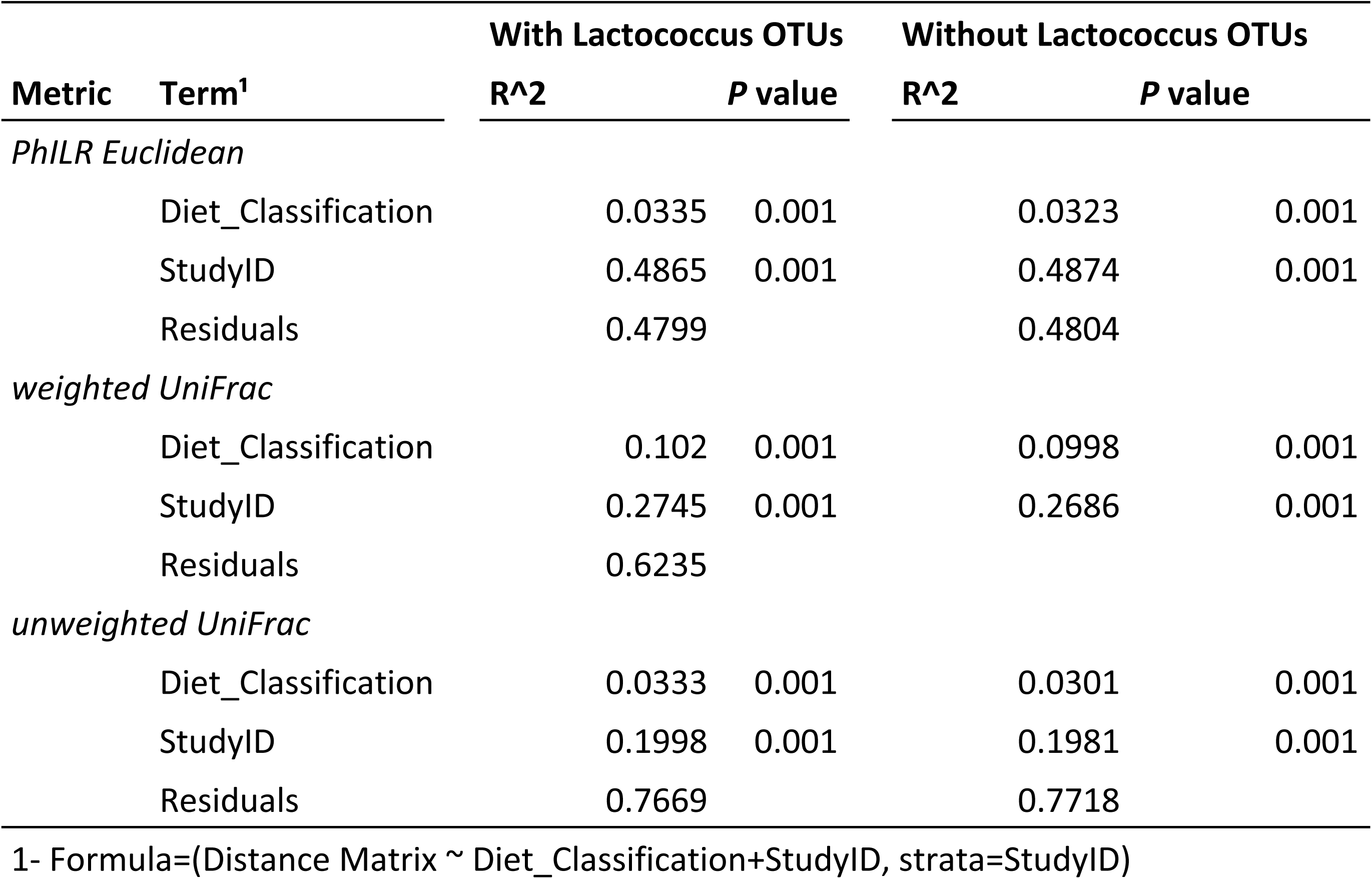
ADONIS tests for entire dataset.

**Supplemental Table 4 relating to Figure 4.**
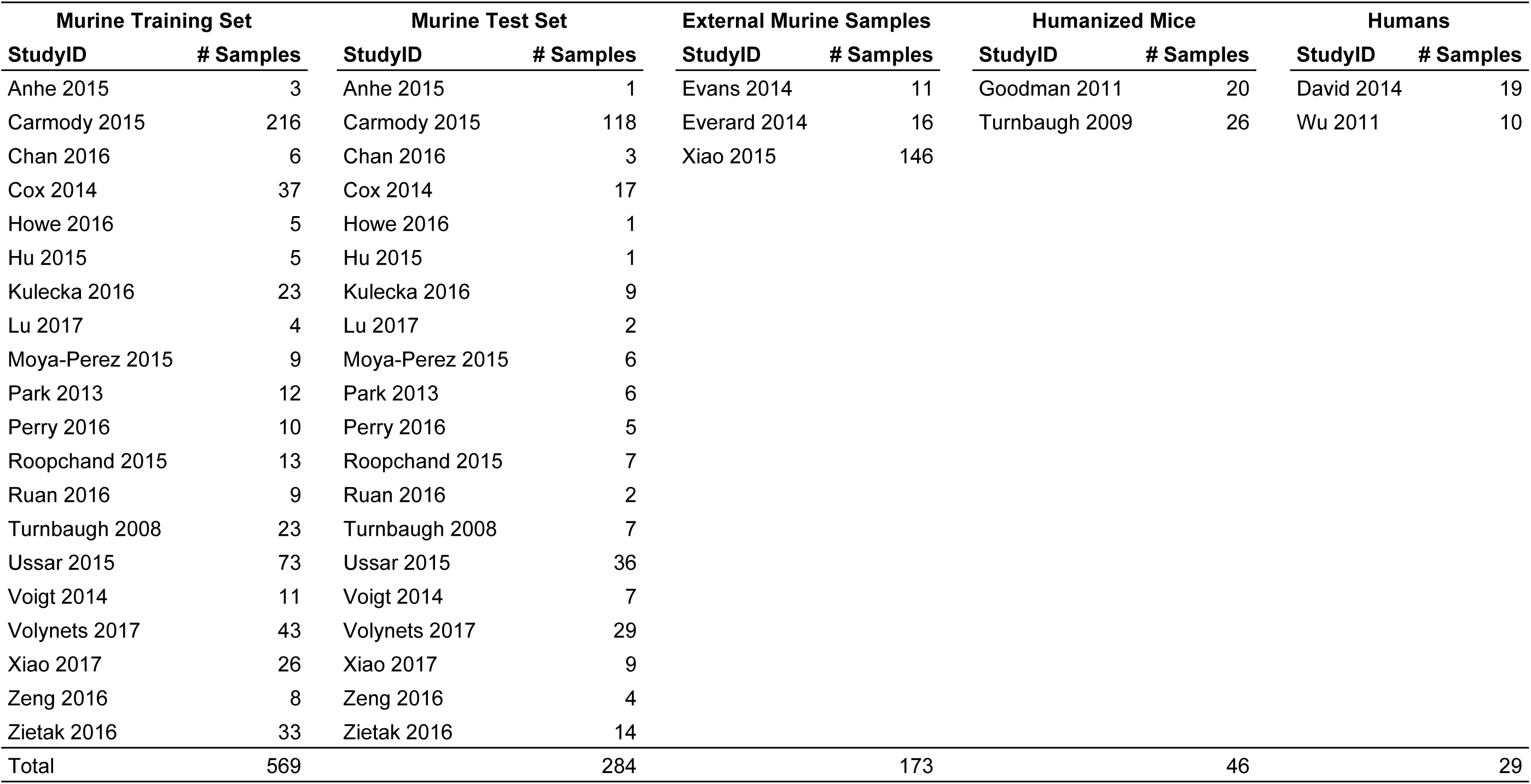
Random Forest Data Sets.

**Supplemental Table 5 relating to Figure 4.**
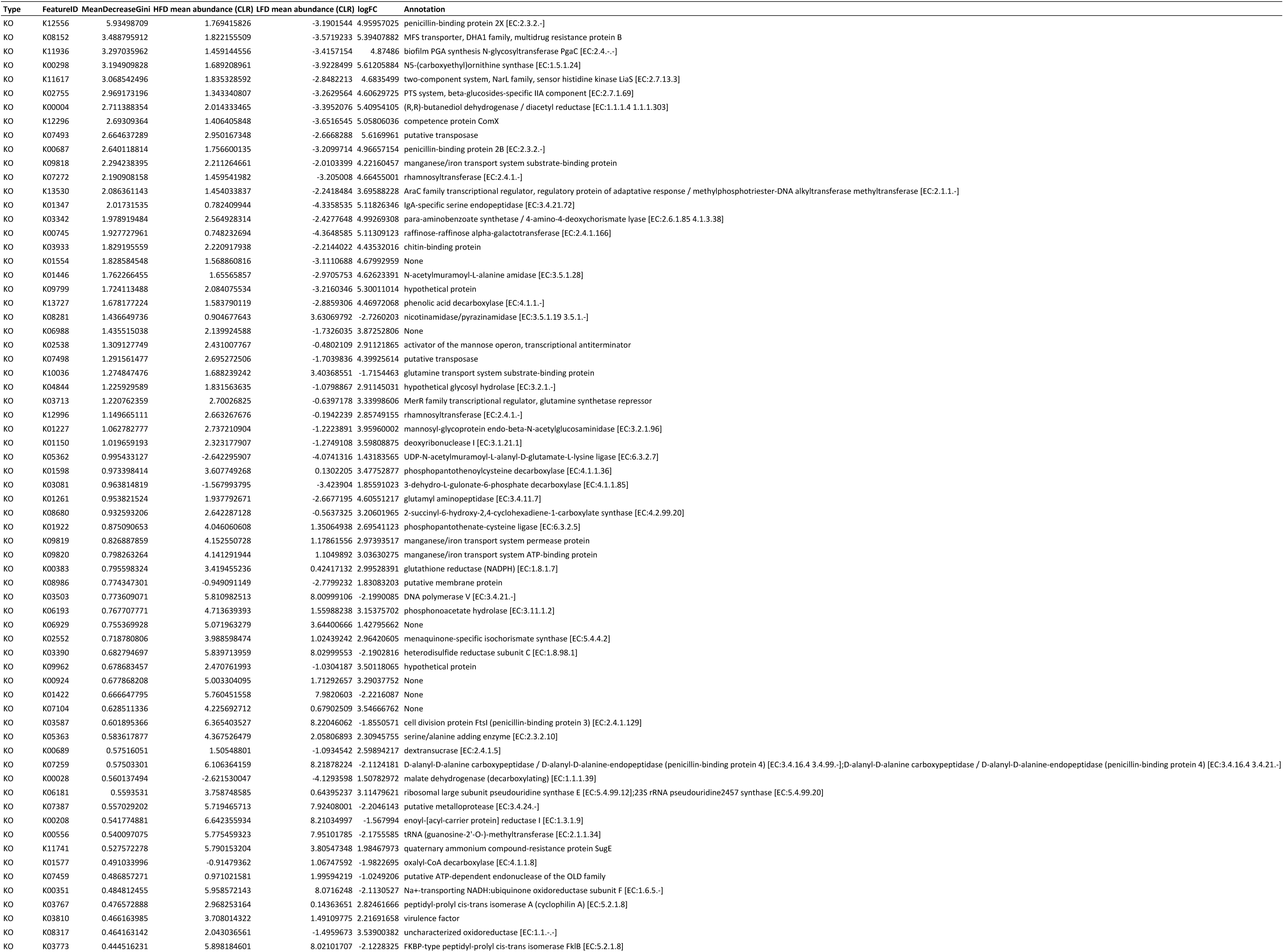
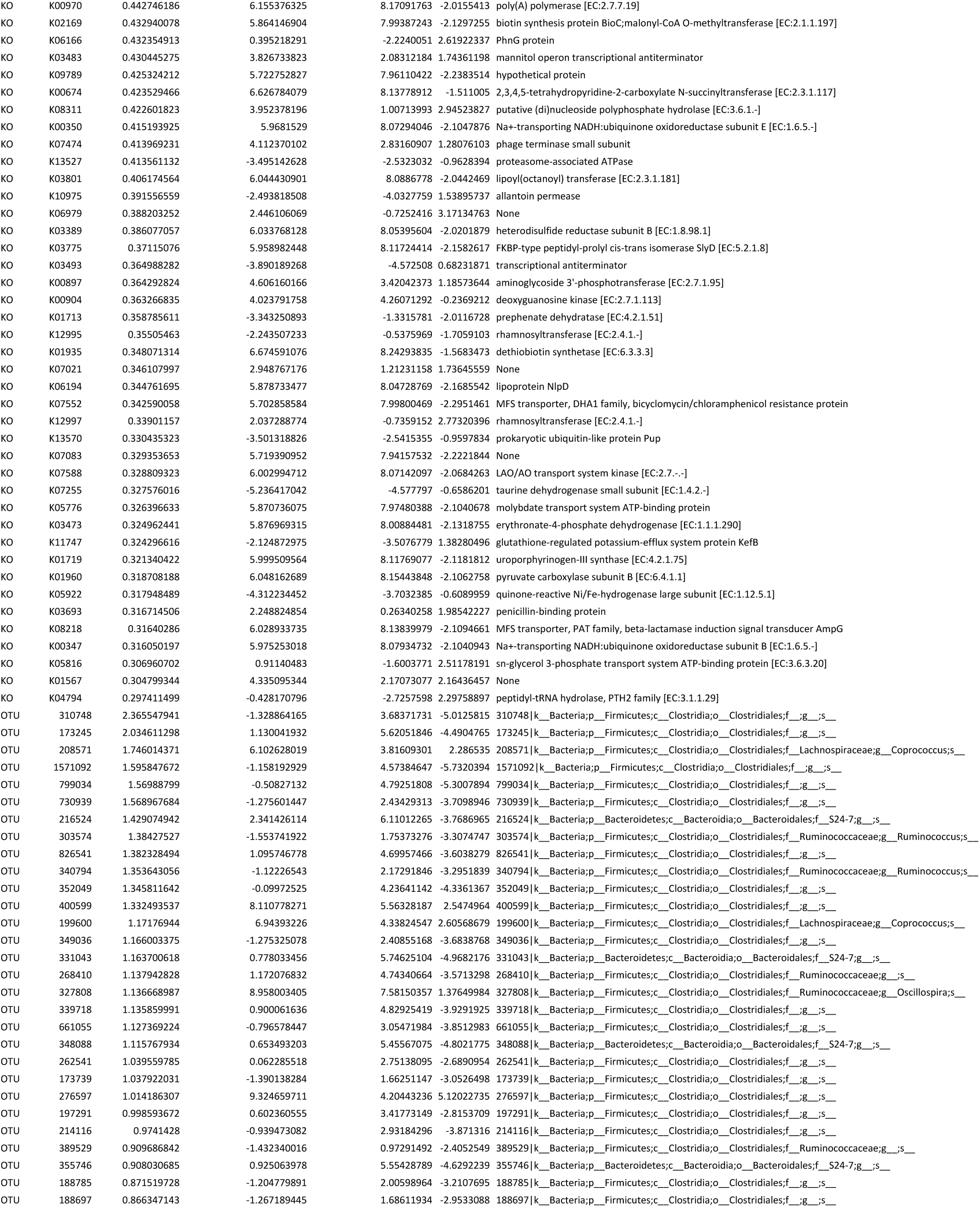
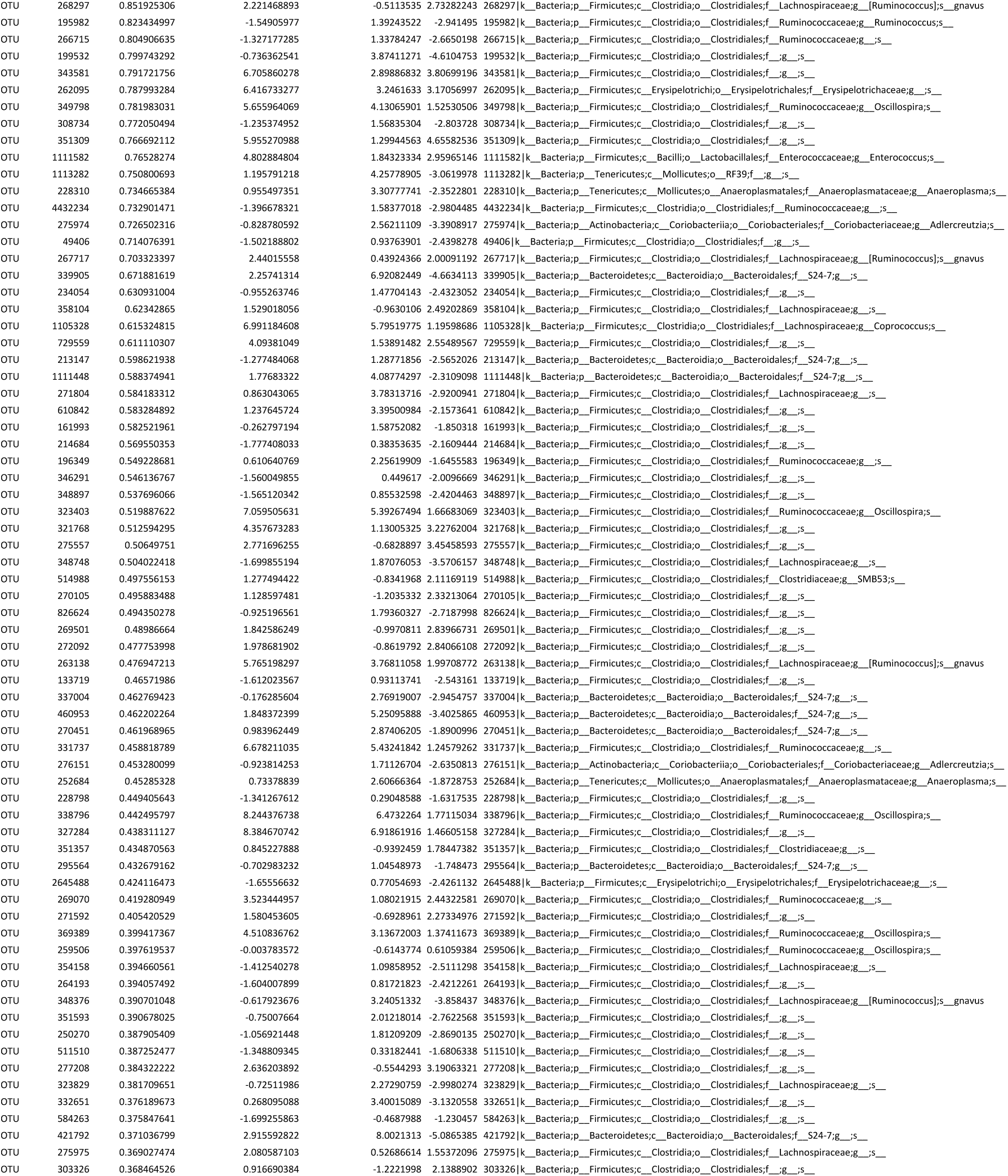
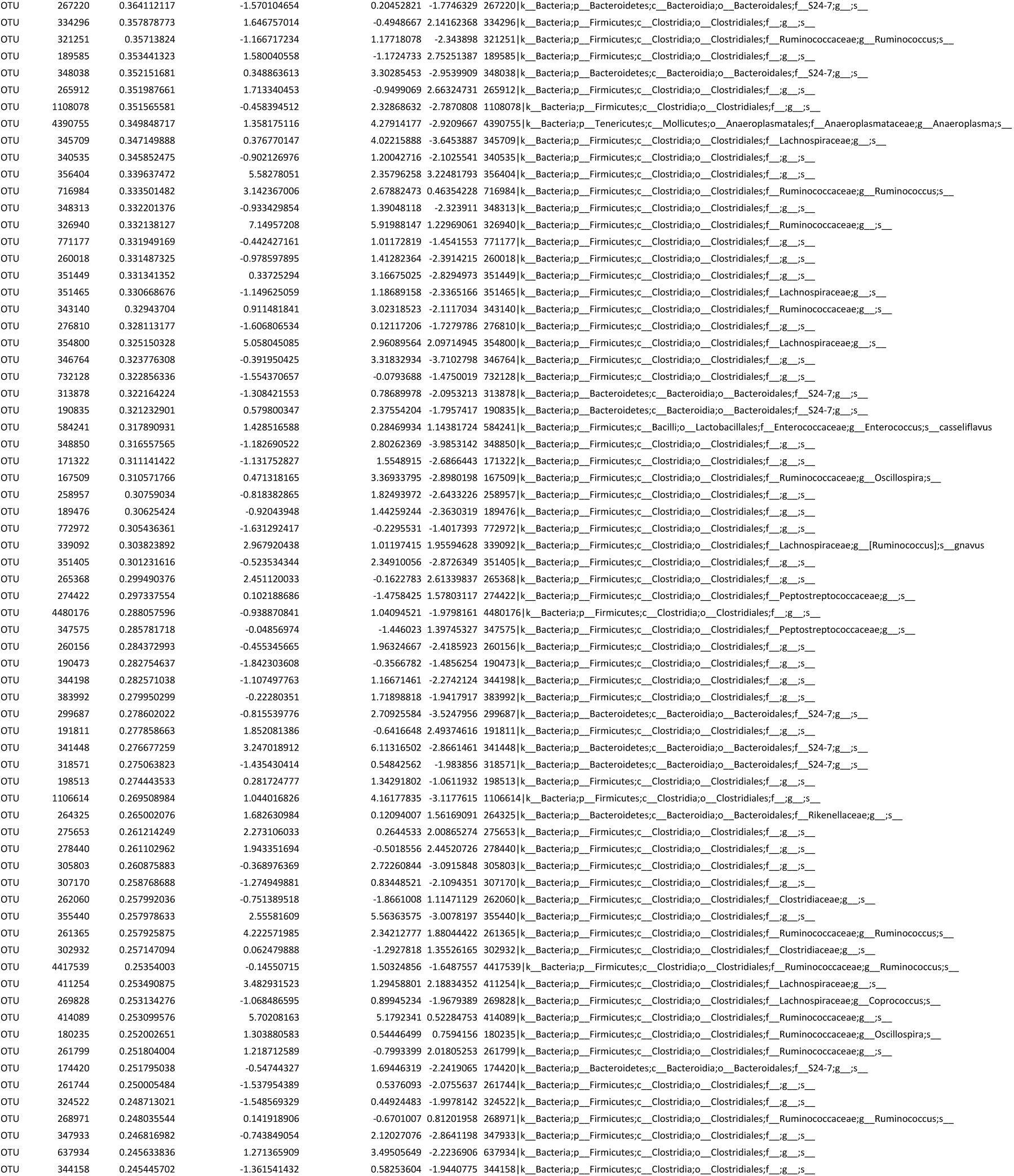
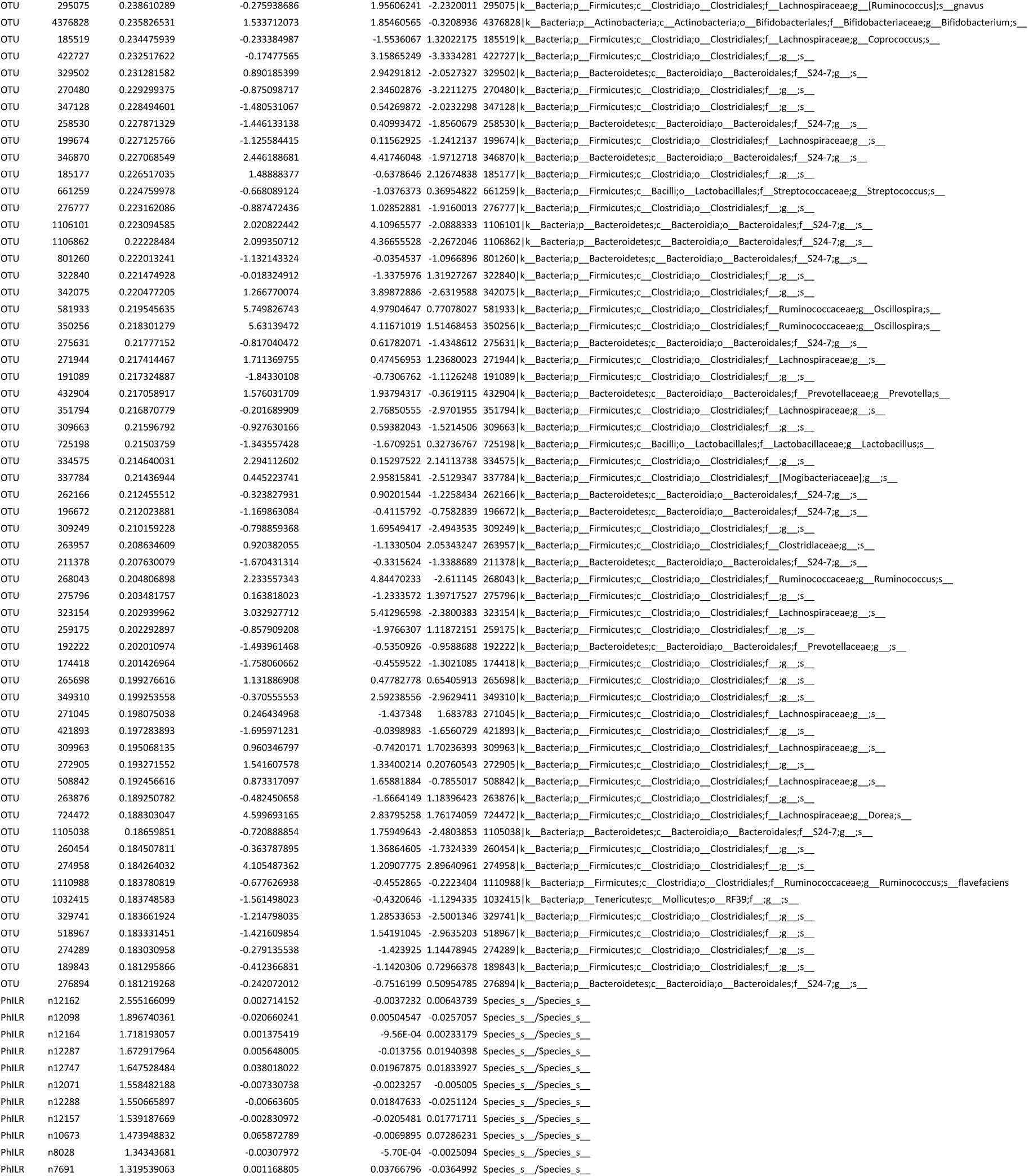
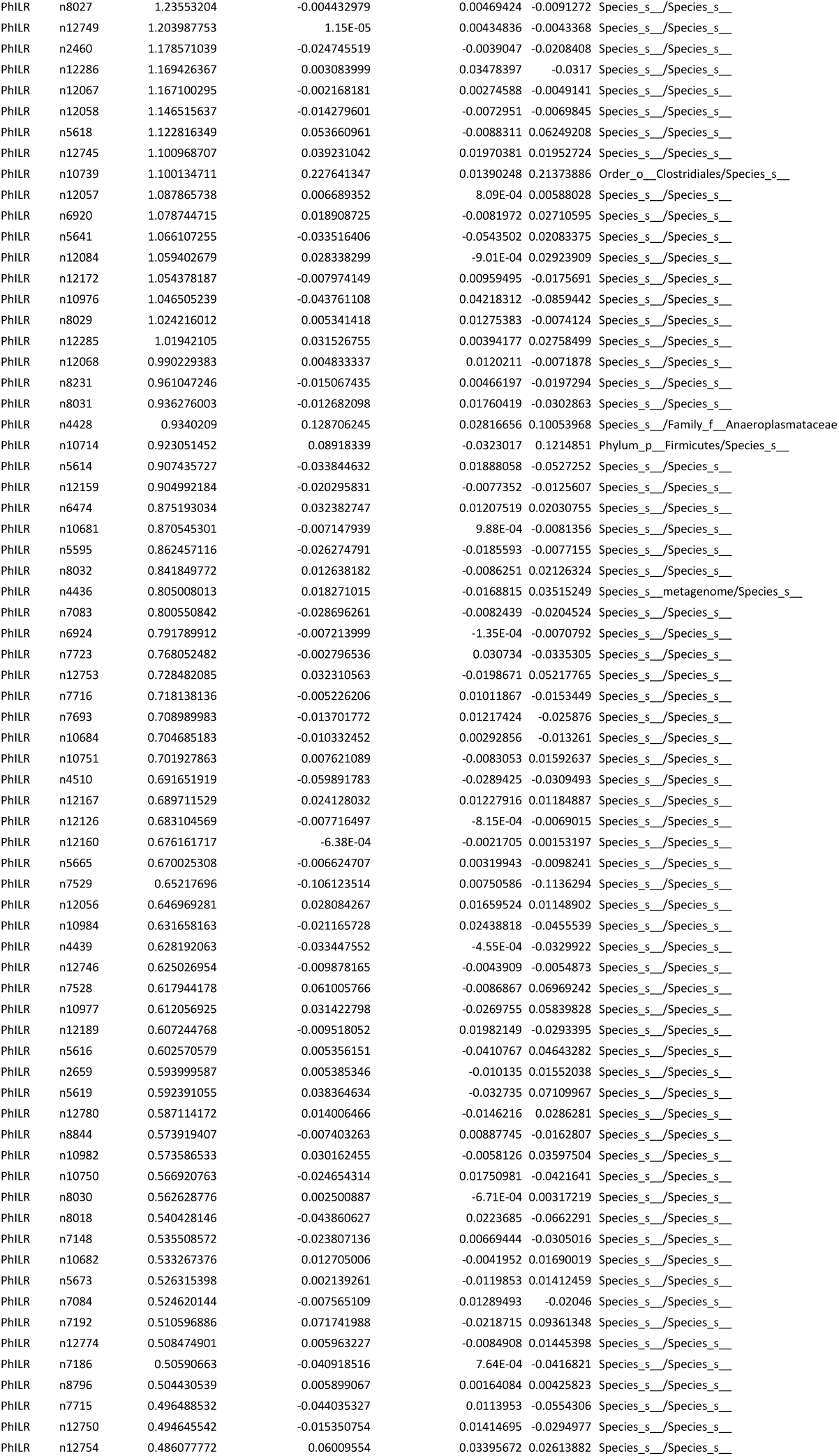
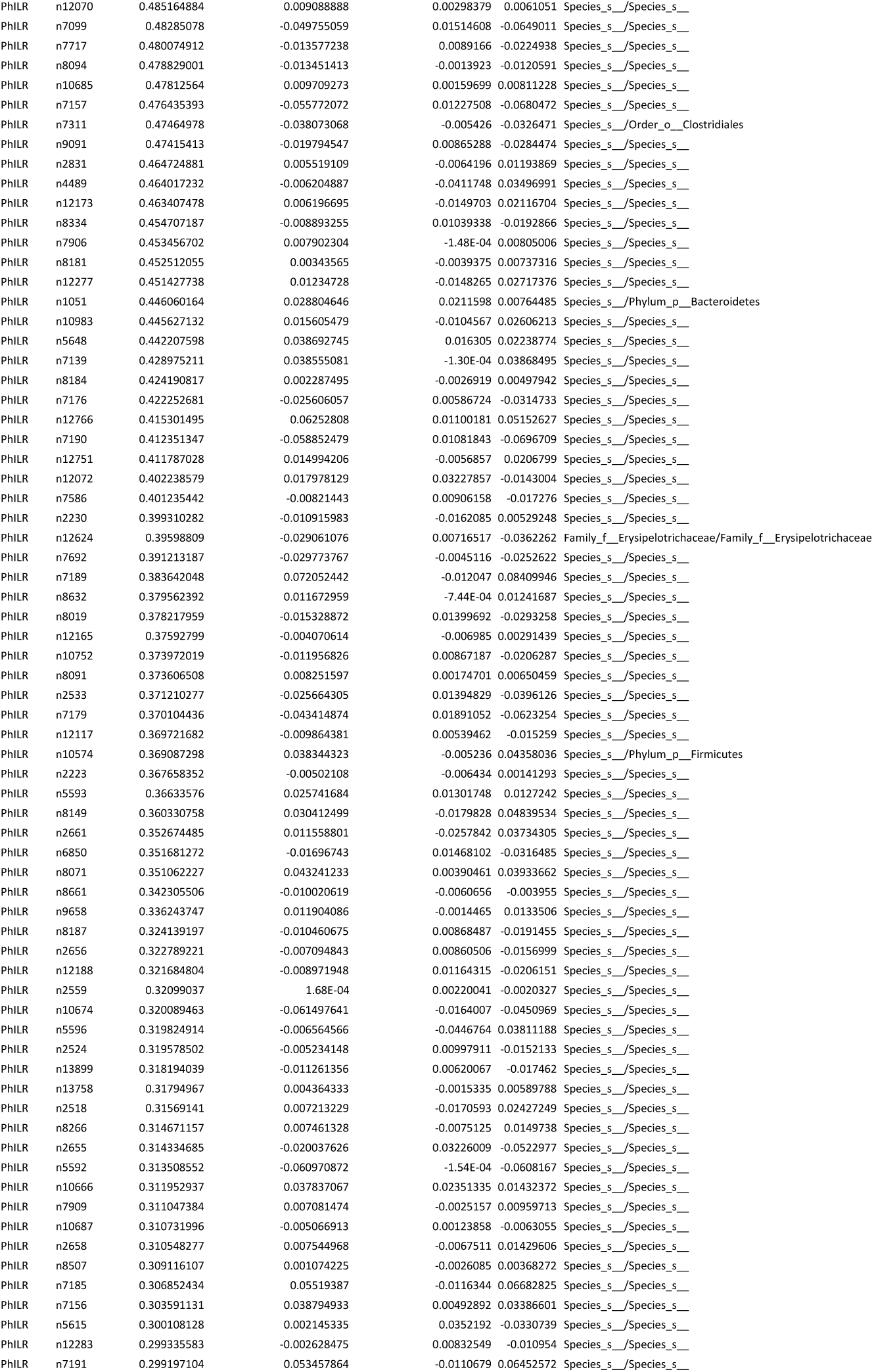
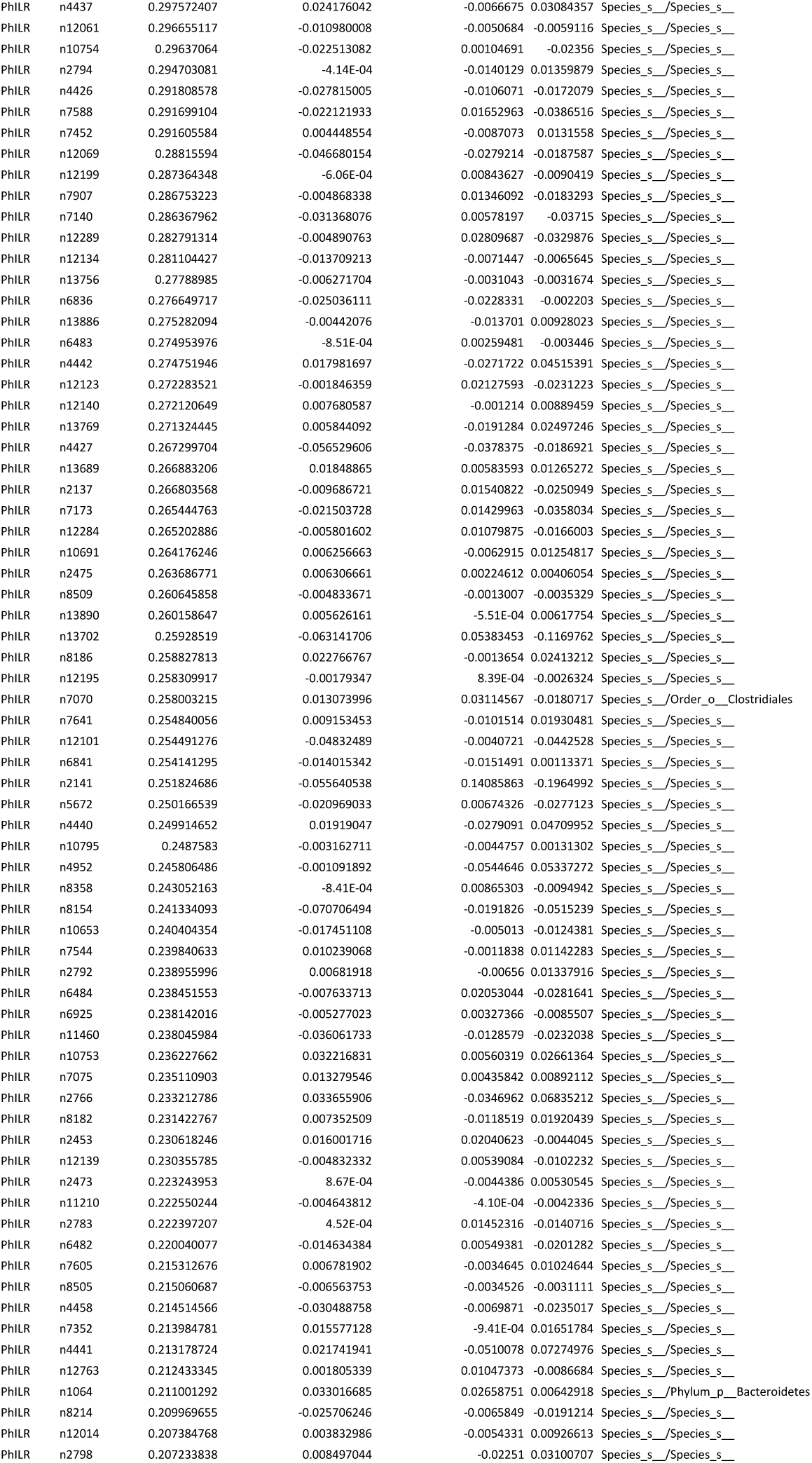
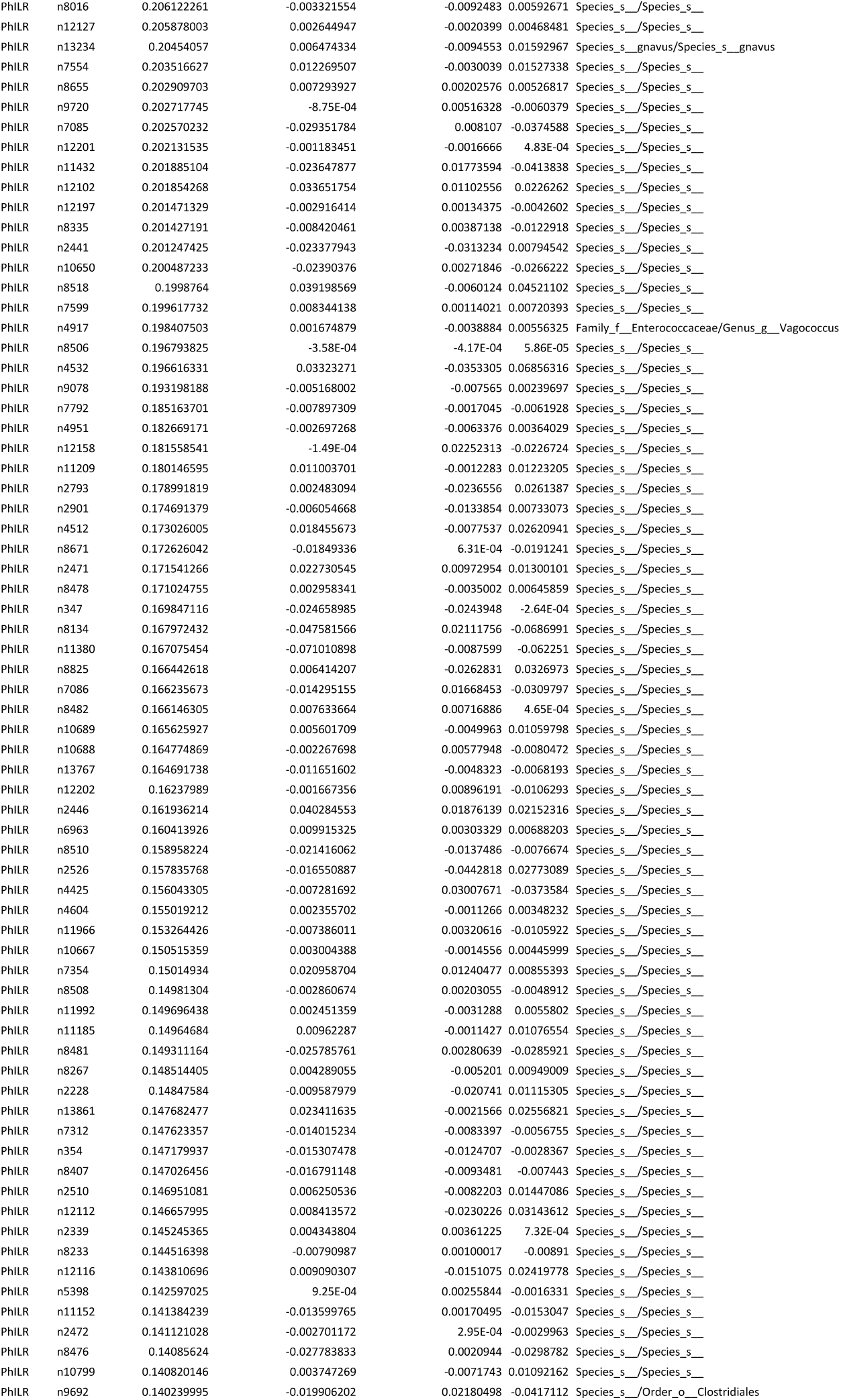
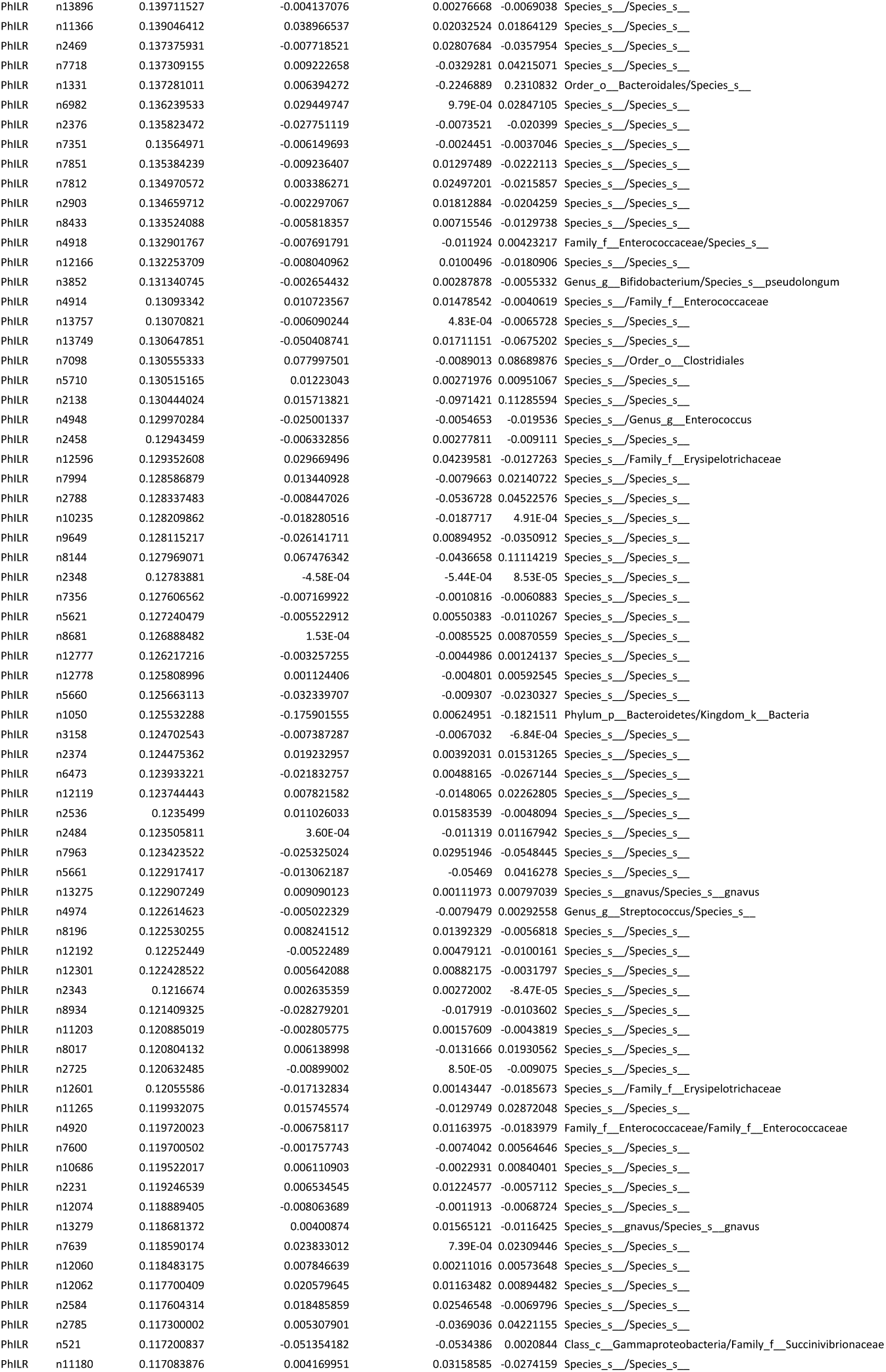
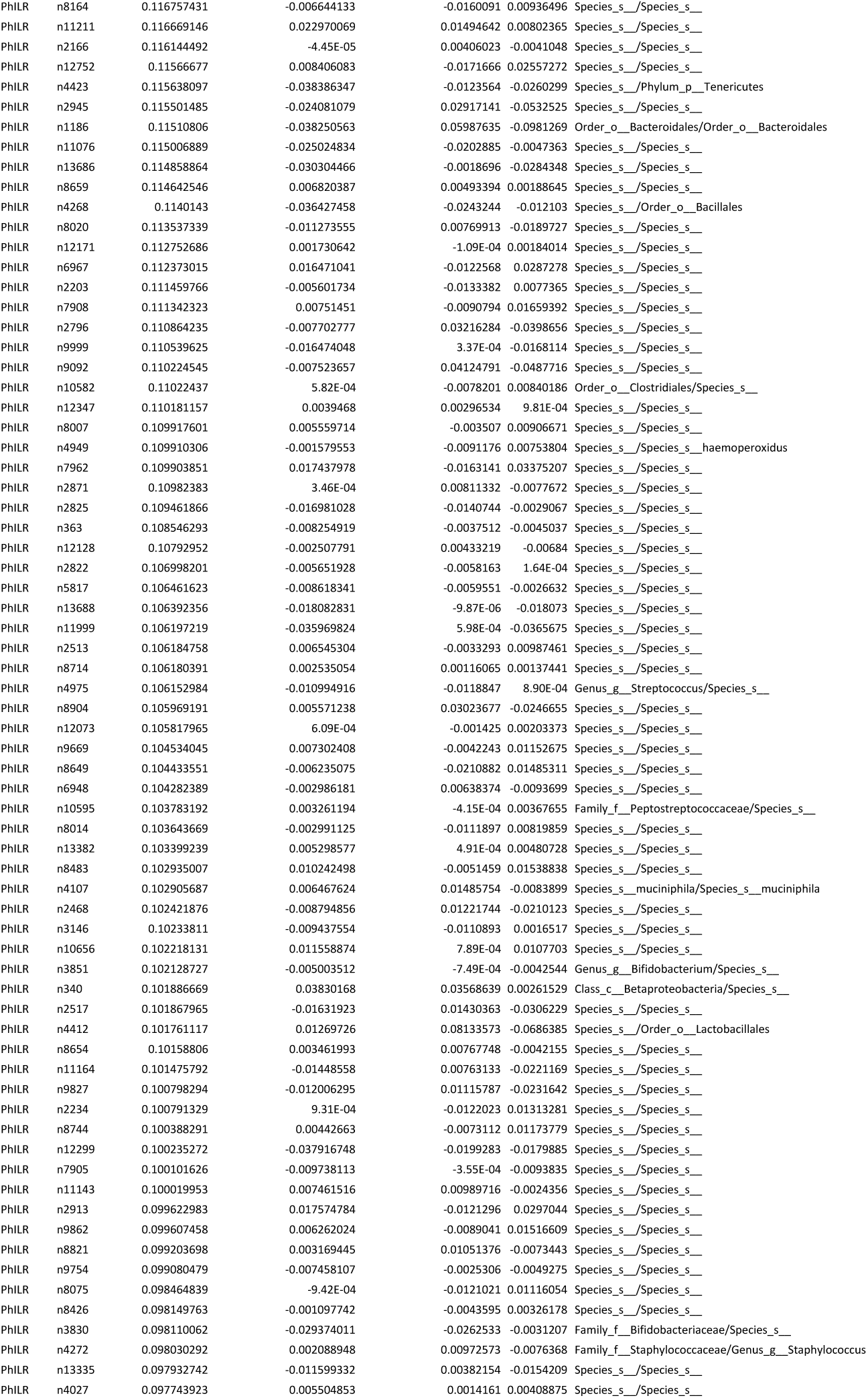
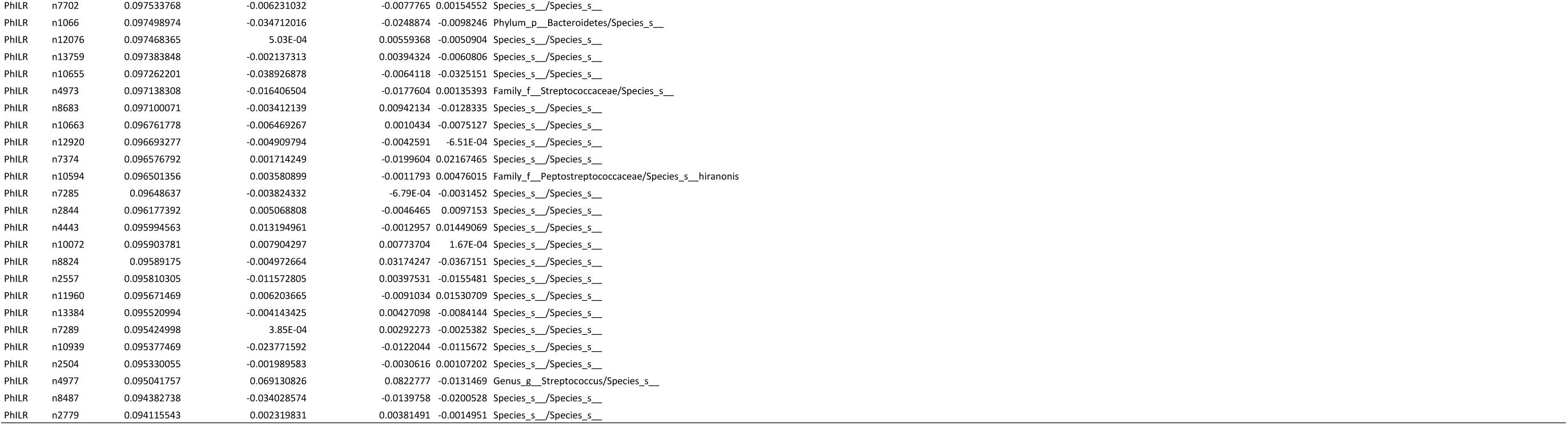
Random forest model features.

**Supplemental Table 6 relating to Figure 4.** Receiver operator curve (ROC) areas under the curve(AUC)

| **Data Set** | **AUC** |  |  |  |
| --- | --- | --- | --- | --- |
|  | **OTU** | **PHILR** | **KO** | **F/B ratio** |
| Murine Training Set | 1.00 | 1.00 | 1.00 | 0.77 |
| Murine Test Set | 0.97 | 0.98 | 0.97 | 0.80 |
| External Murine Sample | 0.94 | 0.90 | 0.78 | 0.79 |
| Humanized Mice | 0.64 | 0.80 | 1.00 | 0.95 |
| Human | 0.60 | 0.75 | 0.88 | 0.43 |

