## Extended Data File 1 for "Diet induces reproducible alterations in the mouse and human gut microbiome": HFD_diversity.html

HFD Meta-analysis: diversity analysis


Code 

- Show All Code
- Hide All Code

### HFD Meta-analysis: diversity analysis

###### *Jordan Bisanz*

#### *2019-01-29 12:56*

### 1 Set up and environment

```
library(tidyverse)
library(readxl)

library(MicrobeR)
library(vegan)
library(phyloseq)
library(philr)

library(doParallel)
library(UpSetR)
library(grid)
library(ggtern)
library(ggtree)
library(ape)
library(picante)
library(randomForest)
library(ROCR)

NSLOTS=6
registerDoParallel(cores=NSLOTS)
knitr::opts_chunk$set(echo = TRUE, message=FALSE, warning=FALSE, tidy=FALSE, cache=FALSE)
HFDcolor="#E69F00"
LFDcolor="#0072B2"
sessionInfo()
```

```
## R version 3.5.0 (2018-04-23)
## Platform: x86_64-apple-darwin15.6.0 (64-bit)
## Running under: macOS  10.14
## 
## Matrix products: default
## BLAS: /Library/Frameworks/R.framework/Versions/3.5/Resources/lib/libRblas.0.dylib
## LAPACK: /Library/Frameworks/R.framework/Versions/3.5/Resources/lib/libRlapack.dylib
## 
## locale:
## [1] en_US.UTF-8/en_US.UTF-8/en_US.UTF-8/C/en_US.UTF-8/en_US.UTF-8
## 
## attached base packages:
## [1] grid      parallel  stats     graphics  grDevices utils     datasets 
## [8] methods   base     
## 
## other attached packages:
##  [1] ROCR_1.0-7          gplots_3.0.1        randomForest_4.6-14
##  [4] picante_1.7         nlme_3.1-137        ape_5.2            
##  [7] ggtree_1.12.0       ggtern_3.0.0        UpSetR_1.4.0       
## [10] doParallel_1.0.11   iterators_1.0.9     foreach_1.4.4      
## [13] philr_1.6.0         phyloseq_1.24.0     vegan_2.5-2        
## [16] lattice_0.20-35     permute_0.9-4       MicrobeR_0.3.1     
## [19] readxl_1.1.0        forcats_0.3.0       stringr_1.3.1      
## [22] dplyr_0.7.5         purrr_0.2.5         readr_1.1.1        
## [25] tidyr_0.8.1         tibble_1.4.2        ggplot2_3.0.0      
## [28] tidyverse_1.2.1    
## 
## loaded via a namespace (and not attached):
##   [1] colorspace_1.3-2    rprojroot_1.3-2     XVector_0.20.0     
##   [4] rstudioapi_0.7      DT_0.4              bit64_0.9-7        
##   [7] lubridate_1.7.4     xml2_1.2.0          codetools_0.2-15   
##  [10] splines_3.5.0       mnormt_1.5-5        robustbase_0.93-2  
##  [13] knitr_1.20          ade4_1.7-11         jsonlite_1.5       
##  [16] broom_0.4.4         cluster_2.0.7-1     latex2exp_0.4.0    
##  [19] compiler_3.5.0      httr_1.3.1          rvcheck_0.1.0      
##  [22] backports_1.1.2     assertthat_0.2.0    Matrix_1.2-14      
##  [25] lazyeval_0.2.1      cli_1.0.0           htmltools_0.3.6    
##  [28] tools_3.5.0         bindrcpp_0.2.2      igraph_1.2.1       
##  [31] gtable_0.2.0        glue_1.2.0          reshape2_1.4.3     
##  [34] fastmatch_1.1-0     Rcpp_0.12.19        Biobase_2.40.0     
##  [37] cellranger_1.1.0    Biostrings_2.48.0   zCompositions_1.1.1
##  [40] multtest_2.36.0     gdata_2.18.0        DECIPHER_2.8.1     
##  [43] psych_1.8.4         tensorA_0.36.1      proto_1.0.0        
##  [46] rvest_0.3.2         phangorn_2.4.0      gtools_3.5.0       
##  [49] DEoptimR_1.0-8      zlibbioc_1.26.0     MASS_7.3-50        
##  [52] scales_0.5.0        hms_0.4.2           biomformat_1.8.0   
##  [55] rhdf5_2.24.0        yaml_2.2.0          memoise_1.1.0      
##  [58] gridExtra_2.3       NADA_1.6-1          stringi_1.2.3      
##  [61] RSQLite_2.1.1       S4Vectors_0.18.3    energy_1.7-5       
##  [64] tidytree_0.1.9      caTools_1.17.1      BiocGenerics_0.26.0
##  [67] boot_1.3-20         truncnorm_1.0-8     bitops_1.0-6       
##  [70] compositions_1.40-2 rlang_0.2.1         pkgconfig_2.0.1    
##  [73] evaluate_0.10.1     Rhdf5lib_1.2.1      bindr_0.1.1        
##  [76] treeio_1.4.1        htmlwidgets_1.2     bit_1.1-14         
##  [79] tidyselect_0.2.4    plyr_1.8.4          magrittr_1.5       
##  [82] R6_2.2.2            IRanges_2.14.10     DBI_1.0.0          
##  [85] pillar_1.2.3        haven_1.1.1         foreign_0.8-70     
##  [88] withr_2.1.2         mgcv_1.8-24         survival_2.42-3    
##  [91] bayesm_3.1-0.1      modelr_0.1.2        crayon_1.3.4       
##  [94] KernSmooth_2.23-15  plotly_4.7.1        rmarkdown_1.10     
##  [97] data.table_1.11.4   blob_1.1.1          digest_0.6.15      
## [100] stats4_3.5.0        munsell_0.5.0       viridisLite_0.3.0  
## [103] quadprog_1.5-5
```

---

### 2 Data Import and normalization

Will generate multiple versions of the OTU table in a list which will include a subsampled, filtered (remove noisy features), 0-replaced, CLR-normalized, and PhILR-normalized versions.

```
metadata<-read_tsv("/Volumes/turnbaughlab/qb3share/jbisanz/HFD_metastudy/CollatedData/metadata_filtered.tsv") %>% mutate(SampleID=DB_Sample) %>% as.data.frame() 
rownames(metadata)<-metadata$DB_Sample

taxonomy<-read_tsv("/Volumes/turnbaughlab/qb3share/jbisanz/HFD_metastudy/dbs/gg_13_8_otus/taxonomy/97_otu_taxonomy.txt", col_names = c("OTU","Taxonomy")) %>%
  separate(Taxonomy, 
           c("Kingdom",
             "Phylum",
             "Class",
             "Order",
             "Family",
             "Genus",
             "Species"
             ), 
           sep="; ", 
           remove=FALSE) %>%
  as.data.frame()
rownames(taxonomy)<-taxonomy$OTU

OTUs<-list()
OTUs$Raw<-read.table("/Volumes/turnbaughlab/qb3share/jbisanz/HFD_metastudy/CollatedData/OTUs.tsv", header=T, sep='\t', row.names=1)
taxonomy<-subset(taxonomy, OTU %in% rownames(OTUs$Raw))

OTUs$Filtered<-Confidence.Filter(OTUs$Raw, 3, 10, TRUE)
OTUs$CZM<-zCompositions::cmultRepl(t(OTUs$Filtered), 
                                   label=0, 
                                   method="CZM", 
                                   output="counts",
                                   suppress.print=TRUE) %>%
                                   t() #with a non-prior based 0 replacement
OTUs$CLR<-apply(log2(OTUs$CZM), 2, function(x)x-mean(x))
OTUs$Subsampled<-Subsample.Table(OTUs$Filtered, VERBOSE=T)
trees<-list()
trees$Raw<-read_tree_greengenes("../../dbs/gg_13_8_otus/trees/97_otus.tree")
trees$Raw$edge.length[is.nan(trees$Raw$edge.length)]<-0 # workaround for import error on a single edge https://forum.qiime2.org/t/greengenes-tree-download-with-branch-lengths/3329/5
trees$Raw<-drop.tip(trees$Raw, tip=trees$Raw$tip.label[!trees$Raw$tip.label %in% rownames(OTUs$Raw)])
trees$PhILR<-drop.tip(trees$Raw, tip=trees$Raw$tip.label[!trees$Raw$tip.label %in% rownames(OTUs$Filtered)])
trees$PhILR<-makeNodeLabel(trees$PhILR, method="number", prefix="n")
trees$Subsampled<-drop.tip(trees$Raw, tip=trees$Raw$tip.label[!trees$Raw$tip.label %in% rownames(OTUs$Subsampled)])
  
OTUs$PhILR<-t(philr(t(OTUs$CZM), trees$PhILR, part.weights='enorm.x.gm.counts', ilr.weights='blw.sqrt'))

saveRDS(OTUs, "RDS/OTUs.RDS")
saveRDS(trees, "RDS/trees.RDS")
saveRDS(taxonomy, "RDS/taxonomy.RDS")
gc()
```

---

### 3 High level visualizations

#### 3.1 Phylum barplots

```
OTUs$TaxaSummary<-Summarize.Taxa(OTUs$Filtered, taxonomy)

phab<-
OTUs$TaxaSummary$Phylum %>%
  Make.Percent() %>%
  as.data.frame() %>%
  rownames_to_column("Phylum") %>%
  as.tibble() %>%
  mutate(Phylum=gsub("..+p__","", Phylum)) %>%
  gather(-Phylum, key="SampleID", value="Abundance")

tp<-
phab %>% group_by(Phylum) %>% summarize(mean=mean(Abundance)) %>%
  arrange(desc(mean)) %>%
  top_n(9, mean) %>%
  bind_rows(., tibble(Phylum="Other", mean=0))

por<-
phab %>%
  filter(Phylum=="Firmicutes") %>%
  arrange(desc(Abundance)) %>%
  pull(SampleID)

phab %>%
  mutate(Phylum=if_else(Phylum %in% tp$Phylum, Phylum, "Other")) %>%
  mutate(Phylum=factor(Phylum, levels = rev(tp$Phylum))) %>%
  left_join(metadata) %>%
  #filter(Colonization=="SPF" | Colonization=="MUSD") %>%
  mutate(SampleID=factor(SampleID, levels=por)) %>%
  ggplot(aes(x=SampleID, y=Abundance, fill=Phylum)) +
  geom_bar(stat="identity") +
  facet_grid(~Diet_Classification, scales="free_x", space = "free") +
  theme_MicrobeR() +
  theme(axis.text.x = element_blank()) +
  theme(axis.ticks.x = element_blank()) +
  coord_cartesian(expand=F) +
  xlab("Sample") +
  ylab("Phylum Abundance (%)") +
  scale_fill_manual(values=rev(c(
    "blue4",
    "olivedrab",
    "firebrick",
    "gold",
    "darkorchid",
    "steelblue2",
    "chartreuse1",
    "aquamarine",
    "coral",
    "grey"
  )))
```

```
ggsave("figures/Phylumplot.pdf", device="pdf", height=3, width=6)
rm(phab, tp, por)
```

#### 3.2 Overlap of OTUs between studies

```
uplot<-
  OTUs$Raw %>%
  as.data.frame() %>%
  rownames_to_column("OTU") %>%
  as_tibble() %>%
  gather(-OTU, key=SampleID, value=Count) %>%
  mutate(Count=if_else(Count==0, 0, 1)) %>%
  left_join(metadata[,c("SampleID","StudyID")]) %>%
  select(-SampleID) %>%
  group_by(StudyID, OTU) %>%
  summarize(Count=max(Count))
  
uplot<-
  uplot %>%
  spread(key=StudyID, value=Count, fill=0) %>%
  as.data.frame() %>%
  upset(., nsets=30, nintersects=300, order.by="freq", show.numbers=F, matrix.dot.alpha=0)
  
uplot
```

```
pdf("figures/upset.pdf", height=11, width=20, useDingbats=F)
  print(uplot)
dev.off()
```

```
## quartz_off_screen 
##                 2
```

```
rm(uplot)
gc()
```

```
##             used  (Mb) gc trigger   (Mb) limit (Mb)  max used   (Mb)
## Ncells   4671539 249.5    7262794  387.9         NA   7262794  387.9
## Vcells 117089858 893.4  327410452 2498.0      16384 406910272 3104.5
```

#### 3.3 Ordinations

##### 3.3.1 Distance Matrix Generation

Then will generate the CLR-euclidian, PhILR-euclidian, weighted and unweighted unifrac, Bray Curtis, and JSD metrics.

```
DistanceMatrices<-list()
DistanceMatrices[["PhILR Euclidian"]]<-dist(t(OTUs$PhILR), method="euclidian"); gc()
DistanceMatrices[["CLR Euclidian"]]<-dist(t(OTUs$CLR), method="euclidian"); gc()
DistanceMatrices[["Bray Curtis"]]<-vegdist(t(Make.Proportion(OTUs$Subsampled)), method="bray"); gc()
DistanceMatrices[["Jensen-Shannon Divergence"]]<-phyloseq::distance(phyloseq(otu_table(Make.Proportion(OTUs$Subsampled), taxa_are_rows = T)), method="jsd", parallel=T); gc()
DistanceMatrices[["weighted UniFrac"]]<-UniFrac(phyloseq(otu_table(Make.Proportion(OTUs$Subsampled), taxa_are_rows = T), phy=trees$Subsampled), weighted=T, parallel=T); gc()
DistanceMatrices[["unweighted UniFrac"]]<-UniFrac(phyloseq(otu_table(Make.Proportion(OTUs$Subsampled), taxa_are_rows = T), phy=trees$Subsampled), weighted=F, parallel=T); gc()
saveRDS(DistanceMatrices, "RDS/DistanceMatrices.RDS")
```

##### 3.3.2 Distance distribution

```
alldists<-
lapply(names(DistanceMatrices), function(dname){
  DistanceMatrices[[dname]] %>%
  as.matrix() %>% 
  as.data.frame() %>%
  rownames_to_column("Subject") %>%
  gather(-Subject, key="Match", value=Distance) %>%
  mutate(Metric=dname)
  }) %>%
  do.call(bind_rows, .)

alldists %>%
  mutate(Metric=factor(Metric, levels=c("PhILR Euclidian", "unweighted UniFrac","weighted UniFrac", "CLR Euclidian","Bray Curtis","Jensen-Shannon Divergence"))) %>%
  ggplot(aes(x=Distance, color=Metric)) +
  geom_freqpoly() +
  theme_MicrobeR() +
  theme(legend.position="none") +
  facet_wrap(~Metric, scales="free")
```

```
ggsave("figures/disthist.pdf", device="pdf", width=7, height=5, useDingbats=F)

rm(alldists)
```

```
lapply(names(DistanceMatrices), function(x){
  DistanceMatrices[[x]] %>%
    as.vector() %>%
    summary() %>%
    broom::tidy() %>%
    mutate(Metric=x) %>%
    select(Metric, everything())
}) %>%
  do.call(bind_rows, .) %>%
  Nice.Table()
```

Because the non-phylogeneticly weighted metrics are so highly saturated in completely non-overlapping samples, will remove them from analysis of the all-sample analysis.

```
DistanceMatrices$`CLR Euclidian`<-NULL
DistanceMatrices$`Bray Curtis`<-NULL
DistanceMatrices$`Jensen-Shannon Divergence`<-NULL
```

##### 3.3.3 Ordinations

Here only ordinating mouse samples that are from SPF mice, or gnotobiotic mice colonized with mouse communities.

```
mousesamps<-metadata %>% filter(Colonization %in% c("SPF")) %>% pull(SampleID)
PCoAs<-list()
for(x in names(DistanceMatrices)){
    message(x)
  PCoAs[[x]]<-
    as.matrix(DistanceMatrices[[x]])[mousesamps,mousesamps] %>%
    as.dist(.) %>%
    ape::pcoa()
}
```

###### 3.3.3.1 Variation Explained

```
lapply(names(PCoAs), function(x){
  PCoAs[[x]]$values %>%
    as.data.frame() %>%
    rownames_to_column("Axis") %>%
    as_tibble() %>%
    mutate(Metric=x) %>%
    mutate(Axis=as.numeric(Axis)) %>%
    filter(Axis<=10)
}) %>%
  do.call(bind_rows, .) %>%
  mutate(Pvar=100*Relative_eig) %>%
  ggplot(aes(x=Axis, y=Pvar, color=Metric)) +
  geom_line() +
  theme_MicrobeR() +
  ylab("% variation explained") +
  xlab("PC") +
  scale_x_continuous(breaks=1:10)
```

```
ggsave("figures/scree.pdf", device="pdf", width=5, height=3, useDingbats=F)
```

```
lapply(names(PCoAs), function(x){
  PCoAs[[x]]$values %>%
    as.data.frame() %>%
    rownames_to_column("Axis") %>%
    as_tibble() %>%
    mutate(Metric=x) %>%
    mutate(Axis=as.numeric(Axis)) %>%
    filter(Axis<=10)
}) %>%
  do.call(bind_rows, .) %>%
  select(Metric, everything()) %>%
  Nice.Table(.)
```

###### 3.3.3.2 Plot by Diet type

```
plotorder<-c("unweighted UniFrac","weighted UniFrac","PhILR Euclidian")

lapply(names(PCoAs), function(x) PCoAs[[x]]$vectors %>% as.data.frame() %>% rownames_to_column("SampleID") %>% select(SampleID, Axis.1, Axis.2, Axis.3) %>% mutate(Metric=x)) %>%
  do.call(bind_rows, .) %>%
  as_tibble() %>%
  left_join(metadata) %>%
  mutate(Metric=factor(Metric, levels=plotorder)) %>%
  ggplot(aes(x=Axis.1, y=Axis.2)) +
  geom_point(aes(color=Diet_Classification), shape=16, alpha=0.3) +
  scale_color_manual(values=c(HFDcolor, LFDcolor)) +
  theme_MicrobeR() +
  facet_wrap(~Metric, scales="free", ncol=4) +
  theme(panel.border = element_blank(), axis.line = element_line()) +
  theme(axis.text = element_blank(), axis.ticks = element_blank()) +
  xlab("") +
  ylab("") +
  theme(legend.position = "none")
```

```
ggsave("figures/PCoAs_diet.pdf", device="pdf", height=2.5, width=6, useDingbats=F)
```

```
lapply(names(PCoAs), function(x) PCoAs[[x]]$vectors %>% as.data.frame() %>% rownames_to_column("SampleID") %>% select(SampleID, Axis.1, Axis.2, Axis.3) %>% mutate(Metric=x)) %>%
  do.call(bind_rows, .) %>%
  as_tibble() %>%
  left_join(metadata) %>%
  mutate(Metric=factor(Metric, levels=plotorder)) %>%
  ggplot(aes(x=Axis.2, y=Axis.3)) +
  geom_point(aes(color=Diet_Classification), shape=16, alpha=0.3) +
  scale_color_manual(values=c(HFDcolor, LFDcolor)) +
  theme_MicrobeR() +
  facet_wrap(~Metric, scales="free", ncol=4) +
  theme(panel.border = element_blank(), axis.line = element_line()) +
  theme(axis.text = element_blank(), axis.ticks = element_blank()) +
  xlab("") +
  ylab("") +
  theme(legend.position = "none")
```

```
ggsave("figures/PCoAs_diet_2v3.pdf", device="pdf", height=2.5, width=6, useDingbats=F)
```

###### 3.3.3.3 Plot by Study

```
ps<-
  lapply(names(PCoAs), function(x) PCoAs[[x]]$vectors %>% as.data.frame() %>% rownames_to_column("SampleID") %>% select(SampleID, Axis.1, Axis.2, Axis.3) %>% mutate(Metric=x)) %>%
  do.call(bind_rows, .) %>%
  as_tibble() %>%
  left_join(metadata) %>%
  mutate(Metric=factor(Metric, levels=plotorder)) %>%
  ggplot(aes(x=Axis.1, y=Axis.2)) +
  geom_point(aes(color=StudyID), shape=16, alpha=0.4) +
  theme_MicrobeR() +
  facet_wrap(~Metric, scales="free", ncol=4) +
  theme(panel.border = element_blank(), axis.line = element_line()) +
  theme(axis.text = element_blank(), axis.ticks = element_blank()) +
  xlab("") +
  ylab("") +
  theme(legend.position = "none")
ps
```

```
ggsave("figures/PCoAs_study.pdf", device="pdf", height=2.5, width=6, useDingbats=F)
```

```
ps<-
  lapply(names(PCoAs), function(x) PCoAs[[x]]$vectors %>% as.data.frame() %>% rownames_to_column("SampleID") %>% select(SampleID, Axis.1, Axis.2, Axis.3) %>% mutate(Metric=x)) %>%
  do.call(bind_rows, .) %>%
  as_tibble() %>%
  left_join(metadata) %>%
  mutate(Metric=factor(Metric, levels=plotorder)) %>%
  ggplot(aes(x=Axis.2, y=Axis.3)) +
  geom_point(aes(color=StudyID), shape=16, alpha=0.4) +
  theme_MicrobeR() +
  facet_wrap(~Metric, scales="free", ncol=4) +
  theme(panel.border = element_blank(), axis.line = element_line()) +
  theme(axis.text = element_blank(), axis.ticks = element_blank()) +
  xlab("PC2") +
  ylab("PC3") +
  theme(legend.position = "none")
ps
```

```
ggsave("figures/PCoAs_study_2v3.pdf", device="pdf", height=2.5, width=6, useDingbats=F)
```

```
ps<-ps + theme(legend.position="right")
ps<-cowplot::get_legend(ps)
plot(ps)
```

```
ggsave("figures/PCoAs_study_legend.pdf", ps, device="pdf", height=5, width=3, useDingbats=F)
```

##### 3.3.4 ADONIS

```
  ADONIS<-tibble()
  for(x in names(DistanceMatrices)){
    message(x)
    td<-as.matrix(DistanceMatrices[[x]])[mousesamps,mousesamps] %>% as.dist()
    tm<-metadata %>% as.data.frame() %>% remove_rownames() %>% column_to_rownames("SampleID")
    tm<-tm[labels(td),]
    ta<-adonis(td~Diet_Classification+StudyID, data=tm, strata=tm$StudyID, permutations=999, parallel=NSLOTS)
      message("ADONIS complete for ",x)
    ta<-ta$aov.tab %>% 
      as.data.frame() %>% 
      rownames_to_column("Term") %>%
      mutate(Metric=x) %>% 
      select(Metric, everything()) %>%
      rename(Pvalue=`Pr(>F)`)
    ADONIS<-bind_rows(ADONIS, ta)
    rm(td, tm, ta)
    gc()
  }
  saveRDS(ADONIS, "RDS/ADONIS.RDS")

write_tsv(ADONIS, "figures/ADONIS_wstrata.txt")
Nice.Table(ADONIS)
```

#### 3.4 Conclusions from high level visualizations

Based on the data generated above, it appears clear that there is some effect of dietary fat on microbiota composition. This can be observed in the PCoA visualizations and the apparent differences in the Firmicutes content seen in the boxplots. These comparison inevitably required a high level of subsampling, and as such further analysis will focus on within study normalization.

```
rm(DistanceMatrices, PCoAs, ps, plotorder, x, mousesamps)
gc()
```

```
##             used  (Mb) gc trigger   (Mb) limit (Mb)  max used   (Mb)
## Ncells   4760324 254.3    7262794  387.9         NA   7262794  387.9
## Vcells 116329933 887.6  327410452 2498.0      16384 406910272 3104.5
```

---

### 4 Per Study Analysis

#### 4.1 processStudy function

To conduct this analysis, will use one function to analyze each study using the same approach but with study-appropriate normalization

```
processStudy<-function(Study){
# PS is the per study object which is a named list with the following objects:
  # Study
  # Metadata
  # AlphaDiversity
  # AlphaDiversity_Stats
  # DistanceMatrices
  # PCoAs
  # ADONISs
  # FBratio
  # OTUs_raw
  # OTUs_subsampled
  # OTUs_clr
  # OTUs_PhILR
  # OTUs_filtered
  # Tree_GG
  # Tree_PhILR
  # Subsample_depth
 PS<-list()
  PS$Study<-Study
  PS$Metadata<-subset(metadata, StudyID==Study)
  PS$OTUs_raw<-OTUs$Raw[,PS$Metadata$SampleID]
  PS$OTUs_raw<-PS$OTUs_raw[rowSums(PS$OTUs_raw)>=1,]
  message(Study, " table has dimensions:", paste(dim(PS$OTUs_raw), collapse="x"))

  
  PS$Subsample_depth<-min(colSums(PS$OTUs_raw))
  PS$OTUs_subsampled<-Subsample.Table(PS$OTUs_raw, DEPTH = PS$Subsample_depth, VERBOSE = T)
  PS$OTUs_filtered<-Confidence.Filter(PS$OTUs_raw, 3, 10)
  
  PS$Tree_GG<-drop.tip(trees$Raw, trees$Raw$tip.label[!trees$Raw$tip.label %in% rownames(PS$OTUs_subsampled)])

  message("FBratioing")
  PS$FBratio<-
    Summarize.Taxa(PS$OTUs_raw, taxonomy %>% as.data.frame() %>% remove_rownames() %>% column_to_rownames("OTU"))$Phylum %>%
    Make.Percent() %>%
    t() %>%
    as.data.frame() %>%
    rownames_to_column("SampleID") %>%
    mutate(FBratio=(`k__Bacteria;p__Firmicutes`+0.1)/(`k__Bacteria;p__Bacteroidetes`+0.1)) %>%
    select(SampleID, FBratio)

  message("Alpha diversity")

  PS$AlphaDiversity<-data.frame(
    Shannon=vegan::diversity(PS$OTUs_subsampled, index="shannon", MARGIN=2),
    Chao1=vegan::estimateR(t(PS$OTUs_subsampled)) %>% t() %>% as.data.frame %>% rename(Chao1=S.chao1) %>% pull(Chao1),
    FaithsPD=picante::pd(t(PS$OTUs_subsampled), PS$Tree_GG, include.root=F)$PD
  )  %>% rownames_to_column("SampleID") %>% 
    select(SampleID, everything()) %>% 
    left_join(PS$FBratio) %>%
    as_tibble() %>%
    gather(-SampleID, key="Metric", value="Diversity") %>%
    left_join(PS$Metadata[,c("SampleID","Diet_Classification", "StudyID")]) %>%
    select(SampleID, StudyID, Diet_Classification, Metric, Diversity)
  
  PS$AlphaDiversity <-
    PS$AlphaDiversity %>%
    left_join(
      PS$AlphaDiversity %>%
      group_by(Metric, Diet_Classification) %>%
      summarize(mean=mean(log2(Diversity))) %>%
      spread(key=Diet_Classification, value=mean) %>%
      rename(mean_log2HFD=HFD, mean_log2LFD=LFD)
    ) %>%
    mutate(log2FC=log2(Diversity)-mean_log2LFD)

  PS$AlphaDiversity_Stats <-
    PS$AlphaDiversity %>%
    group_by(Metric) %>% 
    do(
      broom::tidy(t.test(log2FC~Diet_Classification, data=., conf.int=TRUE, conf.level=0.95))
    ) %>%
    mutate(StudyID=Study) %>%
    select(StudyID, Metric, log2FC=estimate, Pvalue=p.value, mean_LFD=estimate2, mean_HFD=estimate1, CI_low=conf.low, CI_high=conf.high)

  PS$OTUs_clr<-Make.CLR(PS$OTUs_filtered, CZM=TRUE)
  
  PS$DistanceMatrices<-list()
  message("PhILR Euclidian")
  
  PS$Tree_PhILR<-drop.tip(trees$Raw, trees$Raw$tip.label[!trees$Raw$tip.label %in% rownames(PS$OTUs_filtered)])
  PS$Tree_PhILR<-makeNodeLabel(PS$Tree_PhILR, method="number", prefix="n")
  PS$OTUs_PhILR<-t(philr(t(PS$OTUs_filtered+0.5), PS$Tree_PhILR, part.weights='enorm.x.gm.counts', ilr.weights='blw.sqrt'))

  PS$DistanceMatrices[["PhILR Euclidian"]]<-dist(t(PS$OTUs_PhILR), method="euclidian")
    message("CLR Euclidian")
  PS$DistanceMatrices[["CLR Euclidian"]]<-dist(t(PS$OTUs_clr), method="euclidian")
    message("Bray Curtis")
  PS$DistanceMatrices[["Bray Curtis"]]<-vegdist(t(Make.Proportion(PS$OTUs_subsampled)), method="bray")
    message("Wunifrac")
  PS$DistanceMatrices[["weighted UniFrac"]]<-UniFrac(phyloseq(otu_table(Make.Proportion(PS$OTUs_subsampled), taxa_are_rows = T), phy=PS$Tree_GG), weighted=T, parallel=T)
    message("UWunifrac")
  PS$DistanceMatrices[["unweighted UniFrac"]]<-UniFrac(phyloseq(otu_table(Make.Proportion(PS$OTUs_subsampled), taxa_are_rows = T), phy=PS$Tree_GG), weighted=F, parallel=T)
    message("JSD")
  PS$DistanceMatrices[["Jensen-Shannon divergence"]]<-phyloseq::distance(phyloseq(otu_table(Make.Proportion(PS$OTUs_subsampled), taxa_are_rows = T)), method="jsd", parallel=T)
  
  message("PCoAing")
  PS$PCoAs<-lapply(PS$DistanceMatrices, pcoa)
  
  message("ADONISing")

  PS$ADONISs<-lapply(PS$DistanceMatrices, function(x) adonis(x~PS$Metadata[match(labels(x), PS$Metadata$SampleID),]$Diet_Classification))

  PS$ADONISs<-
    lapply(names(PS$ADONISs), function(x){
    PS$ADONISs[[x]]$aov.tab %>% 
      as.data.frame() %>%
      rownames_to_column("Term") %>%
      as.tibble() %>%
      mutate(Metric=x) %>%
      select(Metric, Term, R2, Pvalue=`Pr(>F)`) %>%
      mutate(Term=if_else(Term=="PS$Metadata[match(labels(x), PS$Metadata$SampleID), ]$Diet_Classification", "Diet_Classification", Term))
      }) %>%
    do.call(bind_rows,.)
    gc()
  return(PS)
}
```

#### 4.2 Process Studies

```
  PerStudy<-list()
  for(Study in (unique(metadata$StudyID))){
    message("--------------------->", Study)
    PerStudy[[Study]]<-processStudy(Study)
  }
  saveRDS(PerStudy, "RDS/PerStudy.RDS")
```

#### 4.3 Forest Plot

In the forest plot I will also include all studies pooled. Due to differences in baseline, I will use the log2 fold changes to pool studies and calculate the summed effect.

##### 4.3.1 Alpha div metrics

```
#AlphaCombined<-
#lapply(PerStudy, function(x) x$AlphaDiversity) %>%
#  do.call(bind_rows, .) %>%
#  filter(!StudyID %in% c("David 2014", "Wu 2011")) %>% #remove human samples
#  group_by(Metric) %>%
#  do(
#      broom::tidy(t.test(log2FC~Diet_Classification, data=., conf.int=TRUE, conf.level=0.95))
#    ) %>%
#    mutate(StudyID="Combined") %>%
#    select(StudyID, Metric, log2FC=estimate, Pvalue=p.value, mean_LFD=estimate2, mean_HFD=estimate1, CI_low=conf.low, CI_high=conf.high)

mod<-lapply(PerStudy, function(x) x$AlphaDiversity) %>%
  do.call(bind_rows, .) %>%
  mutate(Diet_Classification=factor(Diet_Classification, levels=c("LFD","HFD"))) %>%
  filter(!StudyID %in% c("David 2014", "Wu 2011")) %>% #remove human samples
  group_by(Metric)

AlphaCombined<-tibble(StudyID=character(0), Metric=character(0), log2FC=numeric(0), Pvalue=numeric(0), mean_LFD=numeric(0), mean_HFD=numeric(0), CI_low=numeric(0), CI_high=numeric(0))
for(i in unique(mod$Metric)){
  fit<-lmerTest:::lmer(log2FC~Diet_Classification+(1|StudyID), data=subset(mod, Metric==i))
  cf<-confint(fit,level = 0.95)
  
  AlphaCombined<-bind_rows(AlphaCombined, tibble(
    StudyID="Combined", 
    Metric=i, 
    log2FC=summary(fit)$coefficients["Diet_ClassificationHFD", "Estimate"], 
    Pvalue=anova(fit)$`Pr(>F)`, 
    mean_LFD=NA, 
    mean_HFD=NA, 
    CI_low=cf["Diet_ClassificationHFD",1], 
    CI_high=cf["Diet_ClassificationHFD",2]
  ))
}


lapply(PerStudy, function(x) x$AlphaDiversity_Stats) %>%
  do.call(bind_rows, .) %>%
    filter(!StudyID %in% c("David 2014", "Wu 2011", "Anhe 2015")) %>% #Anhe 2015 removed as no replicates so any comparison is not really appropriate
  bind_rows(AlphaCombined) %>%
  mutate(Significant=case_when(
    Pvalue<0.05 & log2FC>0 ~ "* up",
    Pvalue<0.05 & log2FC<0 ~ "* down",
    TRUE~"ns"
  )) %>%
  ungroup() %>%
  mutate(Metric=factor(Metric, levels=c("Chao1","Shannon", "FaithsPD","FBratio"))) %>%
  arrange(desc(StudyID)) %>%
  mutate(StudyID=factor(StudyID, levels=c("Combined", grep("Combined", unique(StudyID), invert=T, value=T)))) %>%
  ggplot(aes(x=log2FC, y=StudyID, color=Significant)) +
  geom_vline(xintercept = 0, linetype="dashed", color="grey50") +
  geom_errorbarh(aes(xmin=CI_low, xmax=CI_high), height=0 ) +
  geom_point() +
  facet_grid(~Metric, scales="free_x") +
  theme_MicrobeR() +
  scale_color_manual(values=c(LFDcolor, HFDcolor, "black"))  +
  theme(panel.border = element_blank(), axis.line = element_line()) +
  theme(axis.text.x=element_text(angle=45, hjust=1))
```

```
ggsave("figures/forestalpha.pdf", device="pdf", height=4, width=6, useDingbats=F)
```

###### 4.3.1.1 Combined Alpha Diversity Stats

```
Nice.Table(AlphaCombined)
```

##### 4.3.2 Check for a correlation between Alpha diversity and fat content

```
lapply(PerStudy, function(x) x$AlphaDiversity) %>%
  do.call(bind_rows, .) %>%
  filter(!StudyID %in% c("David 2014", "Wu 2011")) %>%
  left_join(metadata) %>%
  mutate(Diversity=if_else(Metric=="FBratio", log2(Diversity), Diversity)) %>%
  group_by(StudyID, P_Fat, Metric, Diet_Classification) %>%
  summarize(mean=mean(Diversity), sd=sd(Diversity)) %>%
  ungroup() %>%
  mutate(Metric=factor(Metric, levels=c("FBratio","Chao1","Shannon","FaithsPD"))) %>%
  ggplot(aes(x=P_Fat, y=mean, ymin=mean-sd, ymax=mean+sd, fill=Diet_Classification)) +
  geom_smooth(fill="grey80", color="grey50") +
  geom_errorbar(width=0) +
  geom_point(shape=21) +
  facet_wrap(~Metric, scales="free", nrow=1) +
  theme_MicrobeR() +
  xlab("% Fat") +
  ylab("log2(HFD/LFD)±SD") +
  scale_fill_manual(values=c(HFDcolor, LFDcolor)) +
  theme(legend.position = "none")
```

```
ggsave("figures/alphacor.pdf", device="pdf", height=3, width=8, useDingbats=F)
```

```
lapply(PerStudy, function(x) x$AlphaDiversity) %>%
  do.call(bind_rows, .) %>%
  filter(!StudyID %in% c("David 2014", "Wu 2011")) %>%
  left_join(metadata) %>%
  mutate(Diversity=if_else(Metric=="FBratio", log2(Diversity), Diversity)) %>%
  group_by(StudyID, P_Fat, Metric, Diet_Classification) %>%
  summarize(mean=mean(Diversity), sd=sd(Diversity)) %>%
  group_by(Metric) %>%
  do(
    broom::tidy(
      cor.test(.$mean,.$P_Fat, method="spearman")
    )
  ) %>% 
  Nice.Table()
```

While alpha diversity is not correlated with the percentage of dietary fat, the F/B ratio is.

---

##### 4.3.3 Bdiv metrics

```
lapply(names(PerStudy), function(x) PerStudy[[x]]$ADONISs %>% mutate(StudyID=x)) %>%
  do.call(bind_rows, .) %>%
    filter(!StudyID %in% c("David 2014", "Wu 2011", "Anhe 2015")) %>%
  bind_rows(ADONIS %>% mutate(StudyID="Combined")) %>%
  filter(Term=="Diet_Classification") %>%
  ungroup() %>%
  mutate(Metric=gsub("Euclidean","Euclidian", Metric)) %>%
  mutate(Metric=factor(Metric, levels=c("Bray Curtis","weighted UniFrac", "unweighted UniFrac","Jensen-Shannon divergence","CLR Euclidian","PhILR Euclidian"))) %>%
  arrange(desc(StudyID)) %>%
  mutate(StudyID=factor(StudyID, levels=c("Combined", grep("Combined", unique(StudyID), invert=T, value=T)))) %>%
  mutate(Sig=if_else(Pvalue<0.05, "*","ns")) %>%
  ggplot(aes(x=R2, y=StudyID, color=Metric)) +
  geom_point(shape=1, alpha=0.5) +
  geom_point(aes(alpha=Sig), shape=16) +
  theme_MicrobeR() +
  theme(panel.border = element_blank(), axis.line = element_line()) +
  theme(axis.text.x=element_text(angle=45, hjust=1)) +
  scale_alpha_manual(values=c(0.5,0))
```

```
ggsave("figures/forestbeta.pdf", device="pdf", height=4, width=4, useDingbats=F)
```

##### 4.3.4 Ordination separately by study

```
lapply(names(PerStudy), function(x){
  lapply(names(PerStudy[[x]]$PCoAs), function(y){
    PerStudy[[x]]$PCoAs[[y]]$vectors %>% 
      as.data.frame() %>% 
      rownames_to_column("SampleID") %>% 
      mutate(Metric=y) %>%
      select(Metric, SampleID, Axis.1, Axis.2, Axis.3)
  }) %>%
    do.call(bind_rows, .) %>%
    mutate(StudyID=x) %>%
    select(StudyID, everything())
}) %>%
  do.call(bind_rows, .) %>%
  left_join(metadata[,c("SampleID","Diet_Classification")]) %>%
  ggplot(aes(x=Axis.1, y=Axis.2, color=Diet_Classification)) +
  geom_point(alpha=0.5, shape=16) +
  facet_wrap(StudyID~Metric, scales="free", ncol=6) +
  theme_MicrobeR() +
  theme(axis.ticks = element_blank(), axis.text = element_blank()) +
  scale_color_manual(values=c(HFDcolor, LFDcolor)) +
  theme(legend.position="none") +
  xlab("PC1") + ylab("PC2") +
  theme(panel.spacing=unit(0 , "lines"))
```

```
ggsave("figures/PerStudyPCoA.pdf", device="pdf", height=40, width=10, useDingbats=F)
```

```
save.image(paste0("RDS/Session",format(Sys.time(),"%d%b%Y_%H%M%S"),".Rdata"))
gc()
```

```
##             used   (Mb) gc trigger   (Mb) limit (Mb)  max used   (Mb)
## Ncells   5211124  278.4    9407048  502.4         NA   9407048  502.4
## Vcells 136950925 1044.9  377317640 2878.8      16384 471646259 3598.4
```

---

Return to main page
