## Extended Data File 1 for "Diet induces reproducible alterations in the mouse and human gut microbiome": HFD_diversity_nol.html

HFD Meta-analysis: diversity analysis without Lactococcus


Code 

- Show All Code
- Hide All Code

### HFD Meta-analysis: diversity analysis without Lactococcus

###### *Jordan Bisanz*

#### *2019-01-29 13:50*

This document is otherwise identical to HFD\_diversity.Rmd but starts by importing a version of the OTU table that lacks Lactococcus OTUs.

```
metadata<-read_tsv("/Volumes/turnbaughlab/qb3share/jbisanz/HFD_metastudy/CollatedData/metadata_filtered.tsv") %>% mutate(SampleID=DB_Sample) %>% as.data.frame() 
rownames(metadata)<-metadata$DB_Sample

taxonomy<-read_tsv("/Volumes/turnbaughlab/qb3share/jbisanz/HFD_metastudy/dbs/gg_13_8_otus/taxonomy/97_otu_taxonomy.txt", col_names = c("OTU","Taxonomy")) %>%
  separate(Taxonomy, 
           c("Kingdom",
             "Phylum",
             "Class",
             "Order",
             "Family",
             "Genus",
             "Species"
             ), 
           sep="; ", 
           remove=FALSE) %>%
  as.data.frame()
rownames(taxonomy)<-taxonomy$OTU

OTUs<-list()
OTUs$Raw<-read.table("/Volumes/turnbaughlab/qb3share/jbisanz/HFD_metastudy/CollatedData/OTUs_nol.tsv", header=T, sep='\t', row.names=1)
taxonomy<-subset(taxonomy, OTU %in% rownames(OTUs$Raw))

OTUs$Filtered<-Confidence.Filter(OTUs$Raw, 3, 10, TRUE)
OTUs$CZM<-zCompositions::cmultRepl(t(OTUs$Filtered), 
                                   label=0, 
                                   method="CZM", 
                                   output="counts",
                                   suppress.print=TRUE) %>%
                                   t() #with a non-prior based 0 replacement
OTUs$CLR<-apply(log2(OTUs$CZM), 2, function(x)x-mean(x))
OTUs$Subsampled<-Subsample.Table(OTUs$Filtered, VERBOSE=T)
trees<-list()
trees$Raw<-read_tree_greengenes("../../dbs/gg_13_8_otus/trees/97_otus.tree")
trees$Raw$edge.length[is.nan(trees$Raw$edge.length)]<-0 # workaround for import error on a single edge https://forum.qiime2.org/t/greengenes-tree-download-with-branch-lengths/3329/5
trees$Raw<-drop.tip(trees$Raw, tip=trees$Raw$tip.label[!trees$Raw$tip.label %in% rownames(OTUs$Raw)])
trees$PhILR<-drop.tip(trees$Raw, tip=trees$Raw$tip.label[!trees$Raw$tip.label %in% rownames(OTUs$Filtered)])
trees$PhILR<-makeNodeLabel(trees$PhILR, method="number", prefix="n")
trees$Subsampled<-drop.tip(trees$Raw, tip=trees$Raw$tip.label[!trees$Raw$tip.label %in% rownames(OTUs$Subsampled)])
  
OTUs$PhILR<-t(philr(t(OTUs$CZM), trees$PhILR, part.weights='enorm.x.gm.counts', ilr.weights='blw.sqrt'))

saveRDS(OTUs, "RDS/OTUs.RDS")
saveRDS(trees, "RDS/trees.RDS")
saveRDS(taxonomy, "RDS/taxonomy.RDS")
gc()
```

```
pdf("figures/upset.pdf", height=11, width=20, useDingbats=F)
  print(uplot)
dev.off()
```

```
## quartz_off_screen 
##                 2
```

```
rm(uplot)
gc()
```

```
##             used  (Mb) gc trigger   (Mb) limit (Mb)  max used   (Mb)
## Ncells   4671394 249.5    7263018  387.9         NA   7263018  387.9
## Vcells 116758317 890.8  326662569 2492.3      16384 406044288 3097.9
```

```
rm(DistanceMatrices, PCoAs, ps, plotorder, x, mousesamps)
gc()
```

```
##             used  (Mb) gc trigger   (Mb) limit (Mb)  max used   (Mb)
## Ncells   4760176 254.3    7263018  387.9         NA   7263018  387.9
## Vcells 116000022 885.1  326662569 2492.3      16384 406044288 3097.9
```

##### 4.3.1 Alpha div metrics

```
#AlphaCombined<-
#lapply(PerStudy, function(x) x$AlphaDiversity) %>%
#  do.call(bind_rows, .) %>%
#  filter(!StudyID %in% c("David 2014", "Wu 2011")) %>% #remove human samples
#  group_by(Metric) %>%
#  do(
#      broom::tidy(t.test(log2FC~Diet_Classification, data=., conf.int=TRUE, conf.level=0.95))
#    ) %>%
#    mutate(StudyID="Combined") %>%
#    select(StudyID, Metric, log2FC=estimate, Pvalue=p.value, mean_LFD=estimate2, mean_HFD=estimate1, CI_low=conf.low, CI_high=conf.high)

mod<-lapply(PerStudy, function(x) x$AlphaDiversity) %>%
  do.call(bind_rows, .) %>%
  mutate(Diet_Classification=factor(Diet_Classification, levels=c("LFD","HFD"))) %>%
  filter(!StudyID %in% c("David 2014", "Wu 2011")) %>% #remove human samples
  group_by(Metric)

AlphaCombined<-tibble(StudyID=character(0), Metric=character(0), log2FC=numeric(0), Pvalue=numeric(0), mean_LFD=numeric(0), mean_HFD=numeric(0), CI_low=numeric(0), CI_high=numeric(0))
for(i in unique(mod$Metric)){
  fit<-lmerTest:::lmer(log2FC~Diet_Classification+(1|StudyID), data=subset(mod, Metric==i))
  cf<-confint(fit,level = 0.95)
  
  AlphaCombined<-bind_rows(AlphaCombined, tibble(
    StudyID="Combined", 
    Metric=i, 
    log2FC=NA, 
    Pvalue=anova(fit)$`Pr(>F)`, 
    mean_LFD=NA, 
    mean_HFD=NA, 
    CI_low=cf["Diet_ClassificationHFD",1], 
    CI_high=cf["Diet_ClassificationHFD",2]
  ))
}


lapply(PerStudy, function(x) x$AlphaDiversity_Stats) %>%
  do.call(bind_rows, .) %>%
    filter(!StudyID %in% c("David 2014", "Wu 2011", "Anhe 2015")) %>% #Anhe 2015 removed as no replicates so any comparison is not really appropriate
  bind_rows(AlphaCombined) %>%
  mutate(Significant=case_when(
    Pvalue<0.05 & log2FC>0 ~ "* up",
    Pvalue<0.05 & log2FC<0 ~ "* down",
    TRUE~"ns"
  )) %>%
  ungroup() %>%
  mutate(Metric=factor(Metric, levels=c("Chao1","Shannon", "FaithsPD","FBratio"))) %>%
  arrange(desc(StudyID)) %>%
  mutate(StudyID=factor(StudyID, levels=c("Combined", grep("Combined", unique(StudyID), invert=T, value=T)))) %>%
  ggplot(aes(x=log2FC, y=StudyID, color=Significant)) +
  geom_vline(xintercept = 0, linetype="dashed", color="grey50") +
  geom_errorbarh(aes(xmin=CI_low, xmax=CI_high), height=0 ) +
  geom_point() +
  facet_grid(~Metric, scales="free_x") +
  theme_MicrobeR() +
  scale_color_manual(values=c(LFDcolor, HFDcolor, "black"))  +
  theme(panel.border = element_blank(), axis.line = element_line()) +
  theme(axis.text.x=element_text(angle=45, hjust=1))
```

```
ggsave("figures/PerStudyPCoA.pdf", device="pdf", height=40, width=10, useDingbats=F)
```

```
save.image(paste0("RDS/Session",format(Sys.time(),"%d%b%Y_%H%M%S"),".Rdata"))
gc()
```

```
##             used   (Mb) gc trigger   (Mb) limit (Mb)  max used   (Mb)
## Ncells   5210181  278.3    9413071  502.8         NA   9413071  502.8
## Vcells 136542472 1041.8  376456078 2872.2      16384 470569675 3590.2
```

---

Return to main page
