## Extended Data File 1 for "Diet induces reproducible alterations in the mouse and human gut microbiome": HFD_features.html

HFD Meta-analysis: Features


Code 

- Show All Code
- Hide All Code

### HFD Meta-analysis: Features

###### *Jordan Bisanz*

#### *2019-01-29 14:07*

### 1 Set up and environment

```
library(tidyverse)
library(readxl)

library(MicrobeR)
library(philr)

library(ggtree)
library(ape)
library(randomForest)
library(ROCR)

knitr::opts_chunk$set(echo = TRUE, message=FALSE, warning=FALSE, tidy=FALSE, cache=FALSE)
HFDcolor="#E69F00"
LFDcolor="#0072B2"
sessionInfo()
```

```
## R version 3.5.0 (2018-04-23)
## Platform: x86_64-apple-darwin15.6.0 (64-bit)
## Running under: macOS  10.14
## 
## Matrix products: default
## BLAS: /Library/Frameworks/R.framework/Versions/3.5/Resources/lib/libRblas.0.dylib
## LAPACK: /Library/Frameworks/R.framework/Versions/3.5/Resources/lib/libRlapack.dylib
## 
## locale:
## [1] en_US.UTF-8/en_US.UTF-8/en_US.UTF-8/C/en_US.UTF-8/en_US.UTF-8
## 
## attached base packages:
## [1] stats     graphics  grDevices utils     datasets  methods   base     
## 
## other attached packages:
##  [1] ROCR_1.0-7          gplots_3.0.1        randomForest_4.6-14
##  [4] ape_5.2             ggtree_1.12.0       philr_1.6.0        
##  [7] MicrobeR_0.3.1      readxl_1.1.0        forcats_0.3.0      
## [10] stringr_1.3.1       dplyr_0.7.5         purrr_0.2.5        
## [13] readr_1.1.1         tidyr_0.8.1         tibble_1.4.2       
## [16] ggplot2_3.0.0       tidyverse_1.2.1    
## 
## loaded via a namespace (and not attached):
##   [1] colorspace_1.3-2    rprojroot_1.3-2     XVector_0.20.0     
##   [4] rstudioapi_0.7      DT_0.4              bit64_0.9-7        
##   [7] lubridate_1.7.4     xml2_1.2.0          codetools_0.2-15   
##  [10] splines_3.5.0       mnormt_1.5-5        knitr_1.20         
##  [13] ade4_1.7-11         jsonlite_1.5        phyloseq_1.24.0    
##  [16] broom_0.4.4         cluster_2.0.7-1     compiler_3.5.0     
##  [19] httr_1.3.1          rvcheck_0.1.0       backports_1.1.2    
##  [22] assertthat_0.2.0    Matrix_1.2-14       lazyeval_0.2.1     
##  [25] cli_1.0.0           htmltools_0.3.6     tools_3.5.0        
##  [28] bindrcpp_0.2.2      igraph_1.2.1        gtable_0.2.0       
##  [31] glue_1.2.0          reshape2_1.4.3      fastmatch_1.1-0    
##  [34] Rcpp_0.12.19        Biobase_2.40.0      cellranger_1.1.0   
##  [37] Biostrings_2.48.0   zCompositions_1.1.1 multtest_2.36.0    
##  [40] gdata_2.18.0        nlme_3.1-137        DECIPHER_2.8.1     
##  [43] iterators_1.0.9     psych_1.8.4         rvest_0.3.2        
##  [46] phangorn_2.4.0      gtools_3.5.0        zlibbioc_1.26.0    
##  [49] MASS_7.3-50         scales_0.5.0        hms_0.4.2          
##  [52] parallel_3.5.0      biomformat_1.8.0    rhdf5_2.24.0       
##  [55] yaml_2.2.0          memoise_1.1.0       NADA_1.6-1         
##  [58] stringi_1.2.3       RSQLite_2.1.1       S4Vectors_0.18.3   
##  [61] foreach_1.4.4       tidytree_0.1.9      permute_0.9-4      
##  [64] caTools_1.17.1      BiocGenerics_0.26.0 truncnorm_1.0-8    
##  [67] bitops_1.0-6        rlang_0.2.1         pkgconfig_2.0.1    
##  [70] evaluate_0.10.1     lattice_0.20-35     Rhdf5lib_1.2.1     
##  [73] bindr_0.1.1         treeio_1.4.1        htmlwidgets_1.2    
##  [76] bit_1.1-14          tidyselect_0.2.4    plyr_1.8.4         
##  [79] magrittr_1.5        R6_2.2.2            IRanges_2.14.10    
##  [82] picante_1.7         DBI_1.0.0           pillar_1.2.3       
##  [85] haven_1.1.1         foreign_0.8-70      withr_2.1.2        
##  [88] mgcv_1.8-24         survival_2.42-3     modelr_0.1.2       
##  [91] crayon_1.3.4        KernSmooth_2.23-15  plotly_4.7.1       
##  [94] rmarkdown_1.10      grid_3.5.0          data.table_1.11.4  
##  [97] blob_1.1.1          vegan_2.5-2         digest_0.6.15      
## [100] stats4_3.5.0        munsell_0.5.0       viridisLite_0.3.0  
## [103] quadprog_1.5-5
```

---

### 2 Data Import

Here we are reusing data normalized during the diversity analysis. Using CLR normalized OTUs and PICRUSt KOs, and ILR (PhILR).

```
metadata<-read_tsv("/Volumes/turnbaughlab/qb3share/jbisanz/HFD_metastudy/CollatedData/metadata_filtered.tsv") %>% mutate(SampleID=DB_Sample) %>% mutate(Diet_Classification=factor(Diet_Classification, levels=c("HFD","LFD"))) %>% as.data.frame()
rownames(metadata)<-metadata$DB_Sample

OTUs<-readRDS("/Volumes/turnbaughlab/qb3share/jbisanz/HFD_metastudy/Markdown/Diversity/RDS/OTUs.RDS")
trees<-readRDS("/Volumes/turnbaughlab/qb3share/jbisanz/HFD_metastudy/Markdown/Diversity/RDS/trees.RDS")
tree<-trees$PhILR
rm(trees)
taxonomy<-readRDS("/Volumes/turnbaughlab/qb3share/jbisanz/HFD_metastudy/Markdown/Diversity/RDS/taxonomy.RDS")
```

Also getting the KEGG orthology data.

```
KO<-read.table("/Volumes/turnbaughlab/qb3share/jbisanz/HFD_metastudy/CollatedData/PICRUSt.tsv", header=T, row.names=1, sep='\t')
KO<-Make.CLR(KO, PRIOR = 0.5)
```

Will now build a single object containing the data we want to use for downstream analysis.

```
RFdata<-list()
RFdata$KO<-KO
RFdata$OTU<-OTUs$CLR
RFdata$PhILR<-OTUs$PhILR
rm(OTUs, KO)
saveRDS(RFdata, "RDS/RFdata.RDS")
gc()
```

```
##            used  (Mb) gc trigger   (Mb) limit (Mb)  max used   (Mb)
## Ncells  4351706 232.5    7295320  389.7         NA   7225400  385.9
## Vcells 47795726 364.7  155456529 1186.1      16384 175514190 1339.1
```

### 3 Assigning Tasks

Will use a sequential validation approach. Will have 4 groups of studies: \* Human Studies (David 2014 and Wu 2011) \* Humanized mice (from Goodman 2011 and Turnbaugh 2009) \* 3 Randomly selected SPF and MUSD studies to validation across studies (External validation) \* 1/3 validation set of SPF and MUSD mice (Internal Validation) \* 2/3 training set of SPF and MUSD mice (Training Set)

```
set.seed(1811)
ValidStudies<-
  metadata %>%
  filter(!StudyID %in% c("David 2014", "Wu 2011","Goodman 2011")) %>%
  pull(StudyID) %>%
  unique() %>%
  sample(., 3)
print(paste("The following studies will act as randomly selected external validation studies:", paste(ValidStudies, collapse=", "), "."))
```

```
## [1] "The following studies will act as randomly selected external validation studies: Everard 2014, Xiao 2015, Evans 2014 ."
```

```
metadata<-
  metadata %>%
  select(StudyID, SampleID, Species, Background, Colonization, Diet_Classification) %>%
  mutate(Task=case_when(
    StudyID %in% ValidStudies ~ "External Murine Sample",
    Species == "Homo sapiens" ~ "Human",
    Colonization == "HUMD" ~ "Humanized Mice",
    TRUE ~ "Test/Training Set"
  ))

set.seed(1811)
TrainingSet<-
  metadata %>%
    filter(Task=="Test/Training Set") %>%
    pull(SampleID) %>%
    sample(., round(2/3*(length(.)), 0))

metadata<-
  metadata %>%
  mutate(Task=case_when(
    SampleID %in% TrainingSet ~ "Murine Training Set",
    Task == "Test/Training Set" ~ "Murine Test Set",
    TRUE ~ Task
  ))

saveRDS(metadata, "RDS/RFmetadata.RDS")
```

#### 3.1 Summary of assignments

```
metadata %>%
  group_by(Task) %>%
  summarize(Nsample=length(SampleID)) %>%
  arrange(desc(Nsample)) %>%
  Nice.Table()
```

### 4 10-fold cross validation for feature selection

```
CV<-list()
  for(x in names(RFdata)){
    message(date(), " ", x)
    CV[[x]]<-
      rfcv(
          trainx=t(RFdata[[x]][,subset(metadata, Task=="Murine Training Set")$SampleID]),
          trainy=subset(metadata, Task=="Murine Training Set")$Diet_Classification,
          cv.fold=10
    )
  saveRDS(CV, "RDS/CVs.RDS")
  }
```

```
do.call(rbind, lapply(names(CV), function(x) data.frame(Error=CV[[x]]$error.cv) %>% rownames_to_column("nFeature") %>% mutate(Data=x))) %>%
  as.tibble() %>%
  mutate(nFeature=as.numeric(nFeature)) -> CVerror

ggplot(CVerror, aes(x=nFeature, y=Error, group=Data, color=Data)) +
  geom_line() +
  geom_point() +
  theme_classic() +
  theme(legend.position=c(0.8, 0.8)) +
  xlab("# features") +
  ylab("cross-validation error rate (%)") +
  #scale_x_continuous(breaks=seq(0,15000,2000))
  scale_x_continuous(trans="log2", breaks=2^(0:14)) +
  theme(axis.text.x=element_text(angle=45, hjust=1))
```

```
ggsave("figures/cv_curves.pdf", device="pdf", height=4, width=4, useDingbats=F)
```

#### 4.1 Error by number of features

```
CVerror %>% Nice.Table()
```

Based on the results and the sautration that occurs going forward will use the number of features below based on where error rate was saturated in the plots above.

```
NFeats<-list()
NFeats$OTU<-229
NFeats$KO<-216
NFeats$PhILR<-229
```

#### 4.2 Informative Features

Will rerun the random forests and extract the N most important features based on the mean decrease in GINI.

```
FEAT<-list()
  for(x in names(RFdata)){
    message(date(), " ", x)
    FEAT[[x]]<-
      randomForest(
          x=t(RFdata[[x]][,subset(metadata, Task=="Murine Training Set")$SampleID]),
          y=subset(metadata, Task=="Murine Training Set")$Diet_Classification,
          importance=TRUE
    )
  saveRDS(FEAT, "RDS/FEAT.RDS")
  }
```

```
Features<-
lapply(names(FEAT), function(x) {
  FEAT[[x]]$importance %>% 
    as.data.frame() %>% 
    rownames_to_column("Feature") %>% 
    arrange(desc(MeanDecreaseGini)) %>% 
    top_n(NFeats[[x]], MeanDecreaseGini) %>%
    mutate(Type=x)
  }) %>%
  do.call(bind_rows, .) %>%
  select(Type, Feature, everything())
```

Will now annotate these top features.

```
Annotations<-list()
Annotations$KO<-read_tsv("/Volumes/turnbaughlab/qb3share/jbisanz/HFD_metastudy/PICRUSt/mgm4535747.3_mgm4535747.3.picrust", skip=2, col_names=F) %>% mutate(Type="KO") %>% select(Type, Feature=X1, Annotation=X3)
Annotations$OTU<-taxonomy %>% mutate(Annotation=paste0(OTU, "|", gsub(" ","", Taxonomy))) %>% mutate(Type="OTU") %>% mutate(OTU=as.character(OTU)) %>% select(Type, Feature=OTU, Annotation)
Annotations$PhILR<-lapply(subset(Features, Type=="PhILR")$Feature, function(x) tibble(Type="PhILR", Feature=x, Annotation=name.balance(tr=tree, tax=taxonomy, coord=x))) %>% do.call(bind_rows, .)
Features<-Features %>% left_join(do.call(bind_rows, Annotations))
rm(Annotations)
gc()
```

```
##            used  (Mb) gc trigger   (Mb) limit (Mb)  max used   (Mb)
## Ncells  4560839 243.6    7295320  389.7         NA   7295320  389.7
## Vcells 49097452 374.6  155456529 1186.1      16384 175514190 1339.1
```

And finally add an estimate of differential abundance between them which will be the log fold change (mean HFD - mean LFD).

```
FCs<-
rbind(RFdata$KO, RFdata$OTU, RFdata$PhILR)[Features$Feature,] %>%
  as.data.frame() %>%
  rownames_to_column("Feature") %>%
  as_tibble() %>%
  gather(-Feature, key=SampleID, value=Abundance) %>%
  left_join(metadata %>% select(SampleID, Task, Diet_Classification)) %>%
  filter(Task=="Murine Training Set") %>%
  group_by(Feature, Diet_Classification) %>%
  summarize(mean=mean(Abundance)) %>%
  spread(key=Diet_Classification, value=mean) %>%
  mutate(logFC=HFD-LFD) %>%
  ungroup() %>%
  select(Feature, HFD_mean=HFD, LFD_mean=LFD, logFC)

Features<-Features %>% as_tibble() %>% left_join(FCs)
```

#### 4.3 Most Informative Features

```
write_tsv(Features,"figures/MostImportantFeatures.txt")
Nice.Table(Features)
```

---

### 5 Visualizations of most important features

#### 5.1 OTUs

```
Features %>%
  filter(Type=="OTU") %>%
  top_n(20, MeanDecreaseGini) %>%
  mutate(Annotation=gsub("__"," ", Annotation)) %>%
  mutate(Annotation=factor(Annotation, levels=Annotation)) %>%
  ggplot(aes(x=Annotation, y=MeanDecreaseGini, fill=logFC)) +
  geom_bar(stat="identity", color="black") +
  theme_MicrobeR() +
  theme(axis.text.x=element_text(angle=45, hjust=1)) +
  scale_fill_gradient2(low=LFDcolor, high=HFDcolor) +
  theme(legend.position=c(0.9,0.7))
```

```
ggsave("figures/MostImpOTUs.pdf", device="pdf", height=8, width=6, useDingbats=F)
```

From this it is clear that Lactococcus are major drivers of signal.

#### 5.2 PhILR Nodes with OTUs

Here the importance of the node is being denoted by its shading while the OTUs are also being superimposed with their fold change as calculated above.

```
ptree<-drop.tip(tree, tree$tip.label[!tree$tip.label %in% Features$Feature]) #ptree<-tree #reduced for simplified viewing
plottree<-ggtree(ptree, layout = "fan", color="black") #
plottree<-plottree %<+% (Features %>% select(Feature, everything()))
# annotate a set of nodes of interest
interest<-c("p__Bacteroidetes","p__Firmicutes","g__Lactococcus")
for( i in interest){
  tips<-taxonomy %>% filter(grepl(i, Taxonomy)) %>% filter(OTU %in% ptree$tip.label) %>% pull(OTU)
  nd<-getMRCA(phy=ptree, tip=as.character(tips))
  plottree<- 
    plottree + geom_hilight(
    node=nd,
    fill="grey50",
    alpha=0.4
    ) +
    geom_cladelabel(node=nd, label=i)
}

plottree +
  geom_tippoint(aes(size=MeanDecreaseGini, fill=logFC), color="black", shape=21, alpha=0.9) +
  scale_fill_gradient2(low=LFDcolor, high=HFDcolor) +
  geom_nodepoint(aes(size=MeanDecreaseGini)) +
  theme(legend.position="right")
```

```
ggsave("figures/rftree.pdf", device="pdf", height=9, width=11, useDingbats=F)
```

#### 5.3 KOs

```
enrich<-clusterProfiler::enrichKEGG(Features %>% filter(Type=="KO") %>% pull(Feature), organism="ko", pAdjustMethod="BH")

enrich<-as.data.frame(enrich) %>% 
  as.tibble() %>%
  filter(p.adjust<0.1)
Nice.Table(enrich)
```

Now we can plot the network.

```
d<-as.data.frame(enrich)
membership<-strsplit(d$geneID, split="/")
names(membership)<-d$ID
membership<-plyr::ldply(membership, data.frame)
colnames(membership)<-c("Pathway","Feature")
membership<-membership %>% left_join(Features) %>% left_join(d, by=c("Pathway"="ID"))

#http://kateto.net/network-visualization
#For each node, want: ID  \t  FDR \t log2FoldChange
nodes<-Features %>% filter(Type=="KO") %>% select(Feature, MeanDecreaseGini, logFC)
nodes$Type<-"Feature"
nodes<-nodes[nodes$Feature %in% unlist(strsplit(d$geneID, split="/")),]
dd<-d
dd$log2FoldChange<-NA
dd<-dd[,c("Description","log2FoldChange","p.adjust")]
dd$Type<-"Pathway"
colnames(dd)<-colnames(nodes)
nodes<-rbind(nodes, dd)

nodes<-nodes
links<-membership[,c("Description","Feature")]

net <- igraph::graph_from_data_frame(d=links, vertices=nodes, directed=F) 
#plot_network(net) #using phyloseq's code
g<-net
edgeDF <- data.frame(igraph::get.edgelist(g))
    edgeDF$id <- 1:length(edgeDF[, 1])
    vertDF <- igraph::layout.auto(g)
    colnames(vertDF) <- c("x", "y")
    vertDF <- data.frame(value = igraph::get.vertex.attribute(g, "name"),
        vertDF)
    
#extraData = nodes[as.character(vertDF$value), drop = FALSE]
#vertDF <- data.frame(vertDF, extraData)
vertDF<-vertDF %>% left_join(nodes, by=c("value"="Feature"))
    
    graphDF <- merge(reshape2::melt(edgeDF, id = "id"), vertDF, 
        by = "value")
    
    point_size="MeanDecreaseGini"
    color="log2FC"

 # vertDF$FDR.Colonization[vertDF$Type=="Pathway"]=0.001
  #vertDF$log2FoldChange[vertDF$log2FoldChange>6]=6
  #vertDF$log2FoldChange[vertDF$log2FoldChange<(-6)]=-6  
    vertDF<-vertDF %>% mutate(MeanDecreaseGini=if_else(Type=="Pathway", 12.5, MeanDecreaseGini))

    Knet<-
    ggplot() +
    geom_line(data=graphDF, aes(x=x, y=y, group = id), color="grey", alpha=0.4) +
    geom_point(data=vertDF, aes(x,y, fill = logFC, size = log2(MeanDecreaseGini), shape=Type), color="grey80") +
    geom_text(data=subset(vertDF, Type=="Pathway"), aes(x,y, label = value), size = 2, hjust = 0, vjust=1) +
    coord_equal() +
    scale_fill_gradientn(colors=c("cornflowerblue", "white", "indianred", "darkred")) +
    theme_minimal() +
    theme(axis.text.x=element_blank(), axis.text.y=element_blank(),axis.ticks=element_blank()) +
    theme(axis.title.x=element_blank(),axis.title.y=element_blank()) +
    theme(panel.grid.major=element_blank(), panel.grid.minor=element_blank()) +
    scale_shape_manual(values=c(21, 23))

Knet
```

```
ggsave("figures/RFpicrust_wolabs.pdf", Knet, device="pdf", width=8, height=8)

Knet + geom_text(data=subset(vertDF, Type=="Feature"), aes(x,y, label = value), size = 2, hjust = 0, vjust=1)
```

```
ggsave("figures/RFpicrust_wlabs.pdf", device="pdf", width=8, height=8)
```

### 6 Testing models on external data

```
models<-lapply(RFdata, function(x){
     randomForest(
        x=t(x[rownames(x) %in% Features$Feature,subset(metadata, Task=="Murine Training Set")$SampleID]),
        y=subset(metadata, Task=="Murine Training Set")$Diet_Classification)
})

models
```

```
## $KO
## 
## Call:
##  randomForest(x = t(x[rownames(x) %in% Features$Feature, subset(metadata,      Task == "Murine Training Set")$SampleID]), y = subset(metadata,      Task == "Murine Training Set")$Diet_Classification) 
##                Type of random forest: classification
##                      Number of trees: 500
## No. of variables tried at each split: 14
## 
##         OOB estimate of  error rate: 7.21%
## Confusion matrix:
##     HFD LFD class.error
## HFD 258  17  0.06181818
## LFD  24 270  0.08163265
## 
## $OTU
## 
## Call:
##  randomForest(x = t(x[rownames(x) %in% Features$Feature, subset(metadata,      Task == "Murine Training Set")$SampleID]), y = subset(metadata,      Task == "Murine Training Set")$Diet_Classification) 
##                Type of random forest: classification
##                      Number of trees: 500
## No. of variables tried at each split: 15
## 
##         OOB estimate of  error rate: 9.14%
## Confusion matrix:
##     HFD LFD class.error
## HFD 248  27  0.09818182
## LFD  25 269  0.08503401
## 
## $PhILR
## 
## Call:
##  randomForest(x = t(x[rownames(x) %in% Features$Feature, subset(metadata,      Task == "Murine Training Set")$SampleID]), y = subset(metadata,      Task == "Murine Training Set")$Diet_Classification) 
##                Type of random forest: classification
##                      Number of trees: 500
## No. of variables tried at each split: 15
## 
##         OOB estimate of  error rate: 8.26%
## Confusion matrix:
##     HFD LFD class.error
## HFD 254  21  0.07636364
## LFD  26 268  0.08843537
```

In addition, will create a simple logistic model based solely on the F/B ratio.

```
FB<-readRDS("/Volumes/turnbaughlab/qb3share/jbisanz/HFD_metastudy/Markdown/Diversity/RDS/OTUs.RDS")$TaxaSummary$Phylum + 0.5
RFdata$FBratio<-
  FB %>%
  as.data.frame() %>%
  Make.Percent() %>%
  as.data.frame() %>%
  rownames_to_column("Phylum") %>%
  as.tibble() %>%
  filter(grepl("Firmicutes|Bacteroidetes", Phylum)) %>%
  gather(-Phylum, key=Sample, value=abundance) %>%
  spread(key=Phylum, value=abundance) %>%
  mutate(log2FB=log2(`k__Bacteria;p__Firmicutes`/`k__Bacteria;p__Bacteroidetes`)) %>%
  select(SampleID=Sample, log2FB) %>%
  left_join(metadata)
models$FBratio<-glm(Diet_Classification~log2FB, data=subset(RFdata$FBratio, Task=="Murine Training Set"), family=binomial(link="logit"))
models$FBratio
```

```
## 
## Call:  glm(formula = Diet_Classification ~ log2FB, family = binomial(link = "logit"), 
##     data = subset(RFdata$FBratio, Task == "Murine Training Set"))
## 
## Coefficients:
## (Intercept)       log2FB  
##      0.6135      -0.5487  
## 
## Degrees of Freedom: 568 Total (i.e. Null);  567 Residual
## Null Deviance:       788.2 
## Residual Deviance: 648   AIC: 652
```

```
summary(models$FBratio)
```

```
## 
## Call:
## glm(formula = Diet_Classification ~ log2FB, family = binomial(link = "logit"), 
##     data = subset(RFdata$FBratio, Task == "Murine Training Set"))
## 
## Deviance Residuals: 
##     Min       1Q   Median       3Q      Max  
## -1.8721  -1.0163   0.4025   0.9634   2.2939  
## 
## Coefficients:
##             Estimate Std. Error z value Pr(>|z|)    
## (Intercept)  0.61346    0.10828   5.665 1.47e-08 ***
## log2FB      -0.54871    0.05741  -9.558  < 2e-16 ***
## ---
## Signif. codes:  0 '***' 0.001 '**' 0.01 '*' 0.05 '.' 0.1 ' ' 1
## 
## (Dispersion parameter for binomial family taken to be 1)
## 
##     Null deviance: 788.17  on 568  degrees of freedom
## Residual deviance: 648.04  on 567  degrees of freedom
## AIC: 652.04
## 
## Number of Fisher Scoring iterations: 4
```

```
saveRDS(models, "RDS/models.RDS")
```

```
Predictions<-list()
AUCs<-list()
for(Tasks in unique(metadata$Task)){
  Predictions[[Tasks]]<-list()
  AUCs[[Tasks]]<-list()
  for(Types in c("PhILR","KO","OTU","FBratio")){
      message(Tasks,"->",Types)
    if(Types!="FBratio"){
    tm<-predict(models[[Types]], newdata=t(RFdata[[Types]][,subset(metadata, Task==Tasks)$SampleID]), type="prob")[,2] %>%
        prediction(., subset(metadata, Task==Tasks)$Diet_Classification) %>%
        performance(., "tpr","fpr")
    Predictions[[Tasks]][[Types]]<-
        tibble(Task=Tasks, Type=Types, FPR=unlist(tm@x.values), TPR=unlist(tm@y.values))
    
    auc<-predict(models[[Types]], newdata=t(RFdata[[Types]][,subset(metadata, Task==Tasks)$SampleID]), type="prob")[,2] %>%
        prediction(., subset(metadata, Task==Tasks)$Diet_Classification) %>%
        performance(., "auc")
    
      AUCs[[Tasks]][[Types]]<-
        tibble(
          Task=Tasks,
          Type=Types,
          AUC=auc@y.values[[1]]
        )
    } else {
      #for FBratio only
      tm<-predict(object=models[[Types]], newdata=subset(RFdata[[Types]], Task==Tasks ), type="response") %>% # must select the appropriate samples
      prediction(., subset(RFdata[[Types]], Task==Tasks )$Diet_Classification) %>%
      performance(., "tpr","fpr")
    
     Predictions[[Tasks]][[Types]]<-
        tibble(Task=Tasks, Type=Types, FPR=unlist(tm@x.values), TPR=unlist(tm@y.values))
    
    auc<-predict(object=models[[Types]], newdata=subset(RFdata[[Types]], Task==Tasks ), type="response") %>% # must select the appropriate samples
      prediction(., subset(RFdata[[Types]], Task==Tasks )$Diet_Classification) %>%
      performance(., "auc")
    
      AUCs[[Tasks]][[Types]]<-
        tibble(
          Task=Tasks,
          Type=Types,
          AUC=auc@y.values[[1]]
        )
    }
  }
}


Predictions<-lapply(Predictions, function(x) do.call(bind_rows, x)) %>% do.call(bind_rows, .)
AUCs<-lapply(AUCs, function(x) do.call(bind_rows, x)) %>% do.call(bind_rows, .)
```

##### 6.0.1 AUCs

```
Nice.Table(AUCs %>% spread(key=Type, value=AUC))
```

```
write_tsv(AUCs,"figures/AUCs.txt")
```

##### 6.0.2 ROCs

```
Predictions %>%
  mutate(Type=factor(Type, levels=c("OTU","PhILR","FBratio","KO"))) %>%
  mutate(Task=factor(Task, levels=c("Murine Training Set","Murine Test Set","External Murine Sample","Humanized Mice","Human"))) %>%
ggplot(aes(x=FPR, y=TPR, color=Task, group=Task)) +
        geom_abline(color="grey", linetype="dashed") +
        geom_line() +
        xlab("false positive") +
        ylab("true positive") +
        theme_MicrobeR() +
        facet_grid(~Type) +
        theme(axis.text.x=element_text(angle=45, hjust=1))
```

```
ggsave("figures/ROCs.pdf", device="pdf", height=2, width=6, useDingbats=F)
```

---

### 7 Conclusions

Based on the results obtained above, it is clear that OTU and PhILR data does not abstract well to humanized mice or humans. Neither does simple FB ratio. Interestingly, when functional annotations are considered, the model does translate to human samples with decent performance (AUC=0.895).

```
save.image(paste0("RDS/Session",format(Sys.time(),"%d%b%Y_%H%M%S"),".Rdata"))
gc()
```

```
##            used  (Mb) gc trigger   (Mb) limit (Mb)  max used   (Mb)
## Ncells  5226972 279.2    9884600  527.9         NA   9884600  527.9
## Vcells 51197828 390.7  150413752 1147.6      16384 185985415 1419.0
```

---

Return to main page
