## Extended Data File 1 for "Diet induces reproducible alterations in the mouse and human gut microbiome": HFD_features_nol.html

HFD Meta-analysis: Features without Lactococcus


Code 

- Show All Code
- Hide All Code

### HFD Meta-analysis: Features without Lactococcus

###### *Jordan Bisanz*

#### *2019-01-29 14:11*

### 1 Set up and environment

```
library(Biostrings)
library(tidyverse)
library(readxl)

library(MicrobeR)
library(philr)

library(ggtree)
library(ape)
library(randomForest)
library(ROCR)

knitr::opts_chunk$set(echo = TRUE, message=FALSE, warning=FALSE, tidy=FALSE, cache=FALSE)
HFDcolor="#E69F00"
LFDcolor="#0072B2"
sessionInfo()
```

```
## R version 3.5.0 (2018-04-23)
## Platform: x86_64-apple-darwin15.6.0 (64-bit)
## Running under: macOS  10.14
## 
## Matrix products: default
## BLAS: /Library/Frameworks/R.framework/Versions/3.5/Resources/lib/libRblas.0.dylib
## LAPACK: /Library/Frameworks/R.framework/Versions/3.5/Resources/lib/libRlapack.dylib
## 
## locale:
## [1] en_US.UTF-8/en_US.UTF-8/en_US.UTF-8/C/en_US.UTF-8/en_US.UTF-8
## 
## attached base packages:
## [1] stats4    parallel  stats     graphics  grDevices utils     datasets 
## [8] methods   base     
## 
## other attached packages:
##  [1] ROCR_1.0-7          gplots_3.0.1        randomForest_4.6-14
##  [4] ape_5.2             ggtree_1.12.0       philr_1.6.0        
##  [7] MicrobeR_0.3.1      readxl_1.1.0        forcats_0.3.0      
## [10] stringr_1.3.1       dplyr_0.7.5         purrr_0.2.5        
## [13] readr_1.1.1         tidyr_0.8.1         tibble_1.4.2       
## [16] ggplot2_3.0.0       tidyverse_1.2.1     Biostrings_2.48.0  
## [19] XVector_0.20.0      IRanges_2.14.10     S4Vectors_0.18.3   
## [22] BiocGenerics_0.26.0
## 
## loaded via a namespace (and not attached):
##  [1] colorspace_1.3-2    rprojroot_1.3-2     rstudioapi_0.7     
##  [4] DT_0.4              bit64_0.9-7         lubridate_1.7.4    
##  [7] xml2_1.2.0          codetools_0.2-15    splines_3.5.0      
## [10] mnormt_1.5-5        knitr_1.20          ade4_1.7-11        
## [13] jsonlite_1.5        phyloseq_1.24.0     broom_0.4.4        
## [16] cluster_2.0.7-1     compiler_3.5.0      httr_1.3.1         
## [19] rvcheck_0.1.0       backports_1.1.2     assertthat_0.2.0   
## [22] Matrix_1.2-14       lazyeval_0.2.1      cli_1.0.0          
## [25] htmltools_0.3.6     tools_3.5.0         bindrcpp_0.2.2     
## [28] igraph_1.2.1        gtable_0.2.0        glue_1.2.0         
## [31] reshape2_1.4.3      fastmatch_1.1-0     Rcpp_0.12.19       
## [34] Biobase_2.40.0      cellranger_1.1.0    zCompositions_1.1.1
## [37] multtest_2.36.0     gdata_2.18.0        nlme_3.1-137       
## [40] DECIPHER_2.8.1      iterators_1.0.9     psych_1.8.4        
## [43] rvest_0.3.2         phangorn_2.4.0      gtools_3.5.0       
## [46] zlibbioc_1.26.0     MASS_7.3-50         scales_0.5.0       
## [49] hms_0.4.2           biomformat_1.8.0    rhdf5_2.24.0       
## [52] yaml_2.2.0          memoise_1.1.0       NADA_1.6-1         
## [55] stringi_1.2.3       RSQLite_2.1.1       foreach_1.4.4      
## [58] tidytree_0.1.9      permute_0.9-4       caTools_1.17.1     
## [61] truncnorm_1.0-8     bitops_1.0-6        rlang_0.2.1        
## [64] pkgconfig_2.0.1     evaluate_0.10.1     lattice_0.20-35    
## [67] Rhdf5lib_1.2.1      bindr_0.1.1         treeio_1.4.1       
## [70] htmlwidgets_1.2     bit_1.1-14          tidyselect_0.2.4   
## [73] plyr_1.8.4          magrittr_1.5        R6_2.2.2           
## [76] picante_1.7         DBI_1.0.0           pillar_1.2.3       
## [79] haven_1.1.1         foreign_0.8-70      withr_2.1.2        
## [82] mgcv_1.8-24         survival_2.42-3     modelr_0.1.2       
## [85] crayon_1.3.4        KernSmooth_2.23-15  plotly_4.7.1       
## [88] rmarkdown_1.10      grid_3.5.0          data.table_1.11.4  
## [91] blob_1.1.1          vegan_2.5-2         digest_0.6.15      
## [94] munsell_0.5.0       viridisLite_0.3.0   quadprog_1.5-5
```

---

### 2 Data Import

Here we are reusing data normalized during the diversity analysis. Using CLR normalized OTUs and PICRUSt KOs, and ILR (PhILR).

```
metadata<-read_tsv("/Volumes/turnbaughlab/qb3share/jbisanz/HFD_metastudy/CollatedData/metadata_filtered.tsv") %>% mutate(SampleID=DB_Sample) %>% mutate(Diet_Classification=factor(Diet_Classification, levels=c("HFD","LFD"))) %>% as.data.frame()
rownames(metadata)<-metadata$DB_Sample

OTUs<-readRDS("/Volumes/turnbaughlab/qb3share/jbisanz/HFD_metastudy/Markdown/Diversity_nol/RDS/OTUs.RDS")
trees<-readRDS("/Volumes/turnbaughlab/qb3share/jbisanz/HFD_metastudy/Markdown/Diversity_nol/RDS/trees.RDS")
tree<-trees$PhILR
rm(trees)
taxonomy<-readRDS("/Volumes/turnbaughlab/qb3share/jbisanz/HFD_metastudy/Markdown/Diversity_nol/RDS/taxonomy.RDS")
```

Also getting the KEGG orthology data.

```
KO<-read.table("/Volumes/turnbaughlab/qb3share/jbisanz/HFD_metastudy/CollatedData/PICRUSt_nol.tsv", header=T, row.names=1, sep='\t')
KO<-Make.CLR(KO, PRIOR = 0.5)
```

Will now build a single object containing the data we want to use for downstream analysis.

```
RFdata<-list()
RFdata$KO<-KO
RFdata$OTU<-OTUs$CLR
RFdata$PhILR<-OTUs$PhILR
rm(OTUs, KO)
saveRDS(RFdata, "RDS/RFdata.RDS")
gc()
```

```
##            used  (Mb) gc trigger   (Mb) limit (Mb)  max used   (Mb)
## Ncells  4358735 232.8    7681620  410.3         NA   6150575  328.5
## Vcells 47716893 364.1  137735670 1050.9      16384 155909640 1189.5
```

```
NFeats<-list()
NFeats$OTU<-228
NFeats$KO<-108
NFeats$PhILR<-456
```

#### 4.2 Informative Features

Will rerun the random forests and extract the N most important features based on the mean decrease in GINI.

```
FEAT<-list()
  for(x in names(RFdata)){
    message(date(), " ", x)
    FEAT[[x]]<-
      randomForest(
          x=t(RFdata[[x]][,subset(metadata, Task=="Murine Training Set")$SampleID]),
          y=subset(metadata, Task=="Murine Training Set")$Diet_Classification,
          importance=TRUE
    )
  saveRDS(FEAT, "RDS/FEAT.RDS")
  }
```

```
Features<-
lapply(names(FEAT), function(x) {
  FEAT[[x]]$importance %>% 
    as.data.frame() %>% 
    rownames_to_column("Feature") %>% 
    arrange(desc(MeanDecreaseGini)) %>% 
    top_n(NFeats[[x]], MeanDecreaseGini) %>%
    mutate(Type=x)
  }) %>%
  do.call(bind_rows, .) %>%
  select(Type, Feature, everything())
```

Will now annotate these top features.

```
Annotations<-list()
Annotations$KO<-read_tsv("/Volumes/turnbaughlab/qb3share/jbisanz/HFD_metastudy/PICRUSt/mgm4535747.3_mgm4535747.3.picrust", skip=2, col_names=F) %>% mutate(Type="KO") %>% select(Type, Feature=X1, Annotation=X3)
Annotations$OTU<-taxonomy %>% mutate(Annotation=paste0(OTU, "|", gsub(" ","", Taxonomy))) %>% mutate(Type="OTU") %>% mutate(OTU=as.character(OTU)) %>% select(Type, Feature=OTU, Annotation)
Annotations$PhILR<-lapply(subset(Features, Type=="PhILR")$Feature, function(x) tibble(Type="PhILR", Feature=x, Annotation=name.balance(tr=tree, tax=taxonomy, coord=x))) %>% do.call(bind_rows, .)
Features<-Features %>% left_join(do.call(bind_rows, Annotations))
rm(Annotations)
gc()
```

```
##            used  (Mb) gc trigger   (Mb) limit (Mb)  max used   (Mb)
## Ncells  4568502 244.0    7681620  410.3         NA   7681620  410.3
## Vcells 49043858 374.2  137744950 1051.0      16384 155909640 1189.5
```

And finally add an estimate of differential abundance between them which will be the log fold change (mean HFD - mean LFD).

```
FCs<-
rbind(RFdata$KO, RFdata$OTU, RFdata$PhILR)[Features$Feature,] %>%
  as.data.frame() %>%
  rownames_to_column("Feature") %>%
  as_tibble() %>%
  gather(-Feature, key=SampleID, value=Abundance) %>%
  left_join(metadata %>% select(SampleID, Task, Diet_Classification)) %>%
  filter(Task=="Murine Training Set") %>%
  group_by(Feature, Diet_Classification) %>%
  summarize(mean=mean(Abundance)) %>%
  spread(key=Diet_Classification, value=mean) %>%
  mutate(logFC=HFD-LFD) %>%
  ungroup() %>%
  select(Feature, HFD_mean=HFD, LFD_mean=LFD, logFC)

Features<-Features %>% as_tibble() %>% left_join(FCs)
```

#### 4.3 Most Informative Features

```
write_tsv(Features,"figures/MostImportantFeatures.txt")
Nice.Table(Features)
```

---

### 5 Visualizations of most important features

#### 5.1 OTUs

```
Features %>%
  filter(Type=="OTU") %>%
  top_n(20, MeanDecreaseGini) %>%
  mutate(Annotation=gsub("__"," ", Annotation)) %>%
  mutate(Annotation=factor(Annotation, levels=Annotation)) %>%
  ggplot(aes(x=Annotation, y=MeanDecreaseGini, fill=logFC)) +
  geom_bar(stat="identity", color="black") +
  theme_MicrobeR() +
  theme(axis.text.x=element_text(angle=45, hjust=1)) +
  scale_fill_gradient2(low=LFDcolor, high=HFDcolor) +
  theme(legend.position=c(0.9,0.7))
```

```
ggsave("figures/MostImpOTUs.pdf", device="pdf", height=8, width=6, useDingbats=F)
```

#### 5.2 PhILR Nodes with OTUs

Here the importance of the node is being denoted by its shading while the OTUs are also being superimposed with their fold change as calculated above.

```
ptree<-drop.tip(tree, tree$tip.label[!tree$tip.label %in% Features$Feature]) #ptree<-tree #reduced for simplified viewing
plottree<-ggtree(ptree, layout = "fan", color="black") #
plottree<-plottree %<+% (Features %>% select(Feature, everything()))
# annotate a set of nodes of interest
interest<-c("p__Bacteroidetes","p__Firmicutes","g__Lactococcus")
for( i in interest){
  tips<-taxonomy %>% filter(grepl(i, Taxonomy)) %>% filter(OTU %in% ptree$tip.label) %>% pull(OTU)
  nd<-getMRCA(phy=ptree, tip=as.character(tips))
  plottree<- 
    plottree + geom_hilight(
    node=nd,
    fill="grey50",
    alpha=0.4
    ) +
    geom_cladelabel(node=nd, label=i, offset = 0.03)
}

plottree +
  geom_tippoint(aes(size=MeanDecreaseGini, fill=logFC), color="black", shape=21, alpha=0.9) +
  scale_fill_gradient2(low=LFDcolor, high=HFDcolor) +
  geom_nodepoint(aes(size=MeanDecreaseGini)) +
  #geom_nodelab() +
  theme(legend.position="right") +
  geom_cladelabel(node=name.to.nn(ptree, "n12059"), label="x") +
  geom_hilight(node=name.to.nn(ptree, "n12059"),fill="grey50",alpha=0.4) +
  geom_cladelabel(node=name.to.nn(ptree, "n8144"), label="y") +
  geom_hilight(node=name.to.nn(ptree, "n8144"),fill="grey50",alpha=0.4)
```

```
ggsave("figures/rftree.pdf", device="pdf", height=9, width=11, useDingbats=F)
```

In this plot we see what appears to be two highly responsive clades marked x and y above. Will now try to better establish what exactly the taxonomy of these clades is as the Green Genes taxonomy is not informative. Taxonomy of the OTU sequences will be reclassified against a more recent version of the SILVA database using the taxonomic assignment tool of Dada2.

```
Clades<-
bind_rows(
tibble(Clade="y", OTU=extract.clade(ptree, "n8144")$tip.label),
tibble(Clade="x", OTU=extract.clade(ptree, "n12059")$tip.label)
)
gg<-readDNAStringSet("/Volumes/turnbaughlab/qb3share/jbisanz/HFD_metastudy/dbs/gg_13_8_otus/rep_set/97_otus.fasta")
gg<-gg[names(gg) %in% Clades$OTU]
gg<-tibble(OTU=names(gg), Sequence=as.character(gg))

Clades<-Clades %>% left_join(gg) %>% left_join(taxonomy %>% mutate(OTU=as.character(OTU)) %>% select(OTU, GG_taxonomy=Taxonomy))
silva<-dada2::assignTaxonomy(seqs = Clades$Sequence, refFasta = "/Volumes/turnbaughlab/qb3share/jbisanz/HFD_metastudy/dbs/SILVA_132/silva_nr_v132_train_set.fa.gz")
silva<-silva %>% as.data.frame() %>% rownames_to_column("Sequence") %>% mutate(Silva_taxonomy=paste(Kingdom, Phylum, Class, Order, Family, Genus, sep="; ")) %>% select(Sequence, Silva_taxonomy)

Clades<-Clades %>% left_join(silva) %>% select(Clade, OTU, GG_taxonomy, Silva_taxonomy, Sequence)
rm(gg, silva)
Nice.Table(Clades)
```

```
write_tsv(Clades, "figures/Clades.txt")
```

#### 5.3 KOs

```
enrich<-clusterProfiler::enrichKEGG(Features %>% filter(Type=="KO") %>% pull(Feature), organism="ko", pAdjustMethod="BH")

enrich<-as.data.frame(enrich) %>% 
  as.tibble() %>%
  filter(p.adjust<0.1)
Nice.Table(enrich)
```

```
## $KO
## 
## Call:
##  randomForest(x = t(x[rownames(x) %in% Features$Feature, subset(metadata,      Task == "Murine Training Set")$SampleID]), y = subset(metadata,      Task == "Murine Training Set")$Diet_Classification) 
##                Type of random forest: classification
##                      Number of trees: 500
## No. of variables tried at each split: 10
## 
##         OOB estimate of  error rate: 8.08%
## Confusion matrix:
##     HFD LFD class.error
## HFD 256  19  0.06909091
## LFD  27 267  0.09183673
## 
## $OTU
## 
## Call:
##  randomForest(x = t(x[rownames(x) %in% Features$Feature, subset(metadata,      Task == "Murine Training Set")$SampleID]), y = subset(metadata,      Task == "Murine Training Set")$Diet_Classification) 
##                Type of random forest: classification
##                      Number of trees: 500
## No. of variables tried at each split: 15
## 
##         OOB estimate of  error rate: 9.67%
## Confusion matrix:
##     HFD LFD class.error
## HFD 247  28  0.10181818
## LFD  27 267  0.09183673
## 
## $PhILR
## 
## Call:
##  randomForest(x = t(x[rownames(x) %in% Features$Feature, subset(metadata,      Task == "Murine Training Set")$SampleID]), y = subset(metadata,      Task == "Murine Training Set")$Diet_Classification) 
##                Type of random forest: classification
##                      Number of trees: 500
## No. of variables tried at each split: 21
## 
##         OOB estimate of  error rate: 8.26%
## Confusion matrix:
##     HFD LFD class.error
## HFD 252  23  0.08363636
## LFD  24 270  0.08163265
```

```
ggsave("figures/ROCs.pdf", device="pdf", height=2, width=6, useDingbats=F)
```

```
AUCs %>%
  mutate(Type=factor(Type, levels=c("OTU","PhILR","FBratio","KO"))) %>%
  mutate(Task=factor(Task, levels=c("Murine Training Set","Murine Test Set","External Murine Sample","Humanized Mice","Human"))) %>%
  ggplot(aes(x=Task, y=AUC, group=Type, fill=Type)) +
  geom_bar(stat="identity", position=position_dodge(), color="black") +
  theme_MicrobeR() +
  coord_cartesian(ylim=c(0.4,1.02), expand=0) +
  scale_fill_viridis_d()
```

```
ggsave("figures/Auc_plots.pdf", device="pdf", height=2, width=6, useDingbats=F)
```

#### 6.1 Prediction performance on a per study basis

Here will just check if some studies are better predicted than others.

```
#Now Task is really coding for Study
Predictions<-list()
AUCs<-list()
for(Tasks in unique(metadata$StudyID)){
  Predictions[[Tasks]]<-list()
  AUCs[[Tasks]]<-list()
  for(Types in c("PhILR","KO","OTU","FBratio")){
      message(Tasks,"->",Types)
    if(Types!="FBratio"){
    tm<-predict(models[[Types]], newdata=t(RFdata[[Types]][,subset(metadata, StudyID==Tasks)$SampleID]), type="prob")[,2] %>%
        prediction(., subset(metadata, StudyID==Tasks)$Diet_Classification) %>%
        performance(., "tpr","fpr")
    Predictions[[Tasks]][[Types]]<-
        tibble(Task=Tasks, Type=Types, FPR=unlist(tm@x.values), TPR=unlist(tm@y.values))
    
    auc<-predict(models[[Types]], newdata=t(RFdata[[Types]][,subset(metadata, StudyID==Tasks)$SampleID]), type="prob")[,2] %>%
        prediction(., subset(metadata, StudyID==Tasks)$Diet_Classification) %>%
        performance(., "auc")
    
      AUCs[[Tasks]][[Types]]<-
        tibble(
          Task=Tasks,
          Type=Types,
          AUC=auc@y.values[[1]]
        )
    } else {
      #for FBratio only
      tm<-predict(object=models[[Types]], newdata=subset(RFdata[[Types]], StudyID==Tasks ), type="response") %>% # must select the appropriate samples
      prediction(., subset(RFdata[[Types]], StudyID==Tasks )$Diet_Classification) %>%
      performance(., "tpr","fpr")
    
     Predictions[[Tasks]][[Types]]<-
        tibble(Task=Tasks, Type=Types, FPR=unlist(tm@x.values), TPR=unlist(tm@y.values))
    
    auc<-predict(object=models[[Types]], newdata=subset(RFdata[[Types]], StudyID==Tasks ), type="response") %>% # must select the appropriate samples
      prediction(., subset(RFdata[[Types]], StudyID==Tasks )$Diet_Classification) %>%
      performance(., "auc")
    
      AUCs[[Tasks]][[Types]]<-
        tibble(
          Task=Tasks,
          Type=Types,
          AUC=auc@y.values[[1]]
        )
    }
  }
}


Predictions<-lapply(Predictions, function(x) do.call(bind_rows, x)) %>% do.call(bind_rows, .)
AUCs<-lapply(AUCs, function(x) do.call(bind_rows, x)) %>% do.call(bind_rows, .)

ord<-AUCs %>% spread(key=Type, value=AUC) %>% as.data.frame() %>% column_to_rownames("Task") %>% dist(., "euclidian") %>% hclust()

AUCs %>%
  mutate(Type=factor(Type, levels=c("OTU","PhILR","FBratio","KO"))) %>%
 # mutate(Task=factor(Task, levels=rev(unique(Task)))) %>%
  mutate(Task=factor(Task, levels=ord$labels[ord$order])) %>%
  ggplot(aes(y=Task, x=Type, fill=AUC)) +
  geom_tile() +
  scale_fill_viridis_c() +
  theme_MicrobeR() +
  coord_cartesian(expand=0)
```

```
ggsave("figures/Auc_heatmap.pdf", device="pdf", height=5, width=3, useDingbats=F)
write_tsv(AUCs,"figures/AUCs_bystudy.txt")
```

---

### 7 Do Humanized Mice Improve prediction of human samples?

For the purposes of this analysis will skip the feature selection and go straight to model and prediction.

```
models<-lapply(RFdata[1:3], function(x){
     randomForest(
        x=t(x[,subset(metadata, Task=="Humanized Mice")$SampleID]),
        y=subset(metadata, Task=="Humanized Mice")$Diet_Classification)
})
models$FBratio<-glm(Diet_Classification~log2FB, data=subset(RFdata$FBratio, Task=="Humanized Mice"), family=binomial(link="logit"))
summary(models$FBratio)
```

```
## 
## Call:
## glm(formula = Diet_Classification ~ log2FB, family = binomial(link = "logit"), 
##     data = subset(RFdata$FBratio, Task == "Humanized Mice"))
## 
## Deviance Residuals: 
##      Min        1Q    Median        3Q       Max  
## -1.85397  -0.31720   0.06231   0.32639   1.87615  
## 
## Coefficients:
##             Estimate Std. Error z value Pr(>|z|)   
## (Intercept)    4.800      1.644    2.92   0.0035 **
## log2FB        -4.741      1.580   -3.00   0.0027 **
## ---
## Signif. codes:  0 '***' 0.001 '**' 0.01 '*' 0.05 '.' 0.1 ' ' 1
## 
## (Dispersion parameter for binomial family taken to be 1)
## 
##     Null deviance: 63.683  on 45  degrees of freedom
## Residual deviance: 25.728  on 44  degrees of freedom
## AIC: 29.728
## 
## Number of Fisher Scoring iterations: 7
```

```
models
```

```
## $KO
## 
## Call:
##  randomForest(x = t(x[, subset(metadata, Task == "Humanized Mice")$SampleID]),      y = subset(metadata, Task == "Humanized Mice")$Diet_Classification) 
##                Type of random forest: classification
##                      Number of trees: 500
## No. of variables tried at each split: 83
## 
##         OOB estimate of  error rate: 0%
## Confusion matrix:
##     HFD LFD class.error
## HFD  22   0           0
## LFD   0  24           0
## 
## $OTU
## 
## Call:
##  randomForest(x = t(x[, subset(metadata, Task == "Humanized Mice")$SampleID]),      y = subset(metadata, Task == "Humanized Mice")$Diet_Classification) 
##                Type of random forest: classification
##                      Number of trees: 500
## No. of variables tried at each split: 120
## 
##         OOB estimate of  error rate: 4.35%
## Confusion matrix:
##     HFD LFD class.error
## HFD  21   1  0.04545455
## LFD   1  23  0.04166667
## 
## $PhILR
## 
## Call:
##  randomForest(x = t(x[, subset(metadata, Task == "Humanized Mice")$SampleID]),      y = subset(metadata, Task == "Humanized Mice")$Diet_Classification) 
##                Type of random forest: classification
##                      Number of trees: 500
## No. of variables tried at each split: 120
## 
##         OOB estimate of  error rate: 0%
## Confusion matrix:
##     HFD LFD class.error
## HFD  22   0           0
## LFD   0  24           0
## 
## $FBratio
## 
## Call:  glm(formula = Diet_Classification ~ log2FB, family = binomial(link = "logit"), 
##     data = subset(RFdata$FBratio, Task == "Humanized Mice"))
## 
## Coefficients:
## (Intercept)       log2FB  
##       4.800       -4.741  
## 
## Degrees of Freedom: 45 Total (i.e. Null);  44 Residual
## Null Deviance:       63.68 
## Residual Deviance: 25.73     AIC: 29.73
```

```
Predictions<-list()
AUCs<-list()
for(Tasks in unique(metadata$Task)){
  Predictions[[Tasks]]<-list()
  AUCs[[Tasks]]<-list()
  for(Types in c("PhILR","KO","OTU","FBratio")){
      message(Tasks,"->",Types)
    if(Types!="FBratio"){
    tm<-predict(models[[Types]], newdata=t(RFdata[[Types]][,subset(metadata, Task==Tasks)$SampleID]), type="prob")[,2] %>%
        prediction(., subset(metadata, Task==Tasks)$Diet_Classification) %>%
        performance(., "tpr","fpr")
    Predictions[[Tasks]][[Types]]<-
        tibble(Task=Tasks, Type=Types, FPR=unlist(tm@x.values), TPR=unlist(tm@y.values))
    
    auc<-predict(models[[Types]], newdata=t(RFdata[[Types]][,subset(metadata, Task==Tasks)$SampleID]), type="prob")[,2] %>%
        prediction(., subset(metadata, Task==Tasks)$Diet_Classification) %>%
        performance(., "auc")
    
      AUCs[[Tasks]][[Types]]<-
        tibble(
          Task=Tasks,
          Type=Types,
          AUC=auc@y.values[[1]]
        )
    } else {
      #for FBratio only
      tm<-predict(object=models[[Types]], newdata=subset(RFdata[[Types]], Task==Tasks ), type="response") %>% # must select the appropriate samples
      prediction(., subset(RFdata[[Types]], Task==Tasks )$Diet_Classification) %>%
      performance(., "tpr","fpr")
    
     Predictions[[Tasks]][[Types]]<-
        tibble(Task=Tasks, Type=Types, FPR=unlist(tm@x.values), TPR=unlist(tm@y.values))
    
    auc<-predict(object=models[[Types]], newdata=subset(RFdata[[Types]], Task==Tasks ), type="response") %>% # must select the appropriate samples
      prediction(., subset(RFdata[[Types]], Task==Tasks )$Diet_Classification) %>%
      performance(., "auc")
    
      AUCs[[Tasks]][[Types]]<-
        tibble(
          Task=Tasks,
          Type=Types,
          AUC=auc@y.values[[1]]
        )
    }
  }
}


Predictions<-lapply(Predictions, function(x) do.call(bind_rows, x)) %>% do.call(bind_rows, .)
AUCs<-lapply(AUCs, function(x) do.call(bind_rows, x)) %>% do.call(bind_rows, .)
```

##### 7.0.1 AUCs

```
Nice.Table(AUCs %>% spread(key=Type, value=AUC))
```

```
write_tsv(AUCs,"figures/AUCs_hum.txt")
```

##### 7.0.2 ROCs

```
Predictions %>%
  mutate(Type=factor(Type, levels=c("OTU","PhILR","FBratio","KO"))) %>%
  mutate(Task=factor(Task, levels=c("Murine Training Set","Murine Test Set","External Murine Sample","Humanized Mice","Human"))) %>%
ggplot(aes(x=FPR, y=TPR, color=Task, group=Task)) +
        geom_abline(color="grey", linetype="dashed") +
        geom_line() +
        xlab("false positive") +
        ylab("true positive") +
        theme_MicrobeR() +
        facet_grid(~Type) +
        theme(axis.text.x=element_text(angle=45, hjust=1))
```

```
ggsave("figures/ROCs_hum.pdf", device="pdf", height=2, width=6, useDingbats=F)
```

```
AUCs %>%
  mutate(Type=factor(Type, levels=c("OTU","PhILR","FBratio","KO"))) %>%
  mutate(Task=factor(Task, levels=c("Murine Training Set","Murine Test Set","External Murine Sample","Humanized Mice","Human"))) %>%
  ggplot(aes(x=Task, y=AUC, group=Type, fill=Type)) +
  geom_bar(stat="identity", position=position_dodge(), color="black") +
  theme_MicrobeR() +
  coord_cartesian(ylim=c(0.4,1.02), expand=0) +
  scale_fill_viridis_d()
```

```
ggsave("figures/Auc_plots_hum.pdf", device="pdf", height=2, width=6, useDingbats=F)
```

Predictions are not improved, but this could also be a function of the small number of input samples for training.

### 8 Conclusions

After the removal of Lactococcus OTUs which may contribute a false signal, it is now clear that PhILR and KO offer some level of translatability to the response in human gut microbiomes.

```
save.image(paste0("RDS/Session",format(Sys.time(),"%d%b%Y_%H%M%S"),".Rdata"))
gc()
```

```
##            used  (Mb) gc trigger   (Mb) limit (Mb)  max used   (Mb)
## Ncells  7830775 418.3   13616077  727.2         NA  13616077  727.2
## Vcells 55646502 424.6  165971355 1266.3      16384 165971355 1266.3
```

---

Return to main page
