## Extended Data File 1 for "Diet induces reproducible alterations in the mouse and human gut microbiome": HFD_process.html

HFD meta-analysis: Downloading data and processing samples


Code 

- Show All Code
- Hide All Code

### HFD meta-analysis: Downloading data and processing samples

###### *Jordan E. Bisanz*

#### *2019-01-29 12:52*

### 1 Background

This document describes the methodology for obtaining raw data and closed-reference OTU picking. All code is run either in R or bash.

### 2 Environment

```
library(tidyverse)
library(readxl)
library(MicrobeR)
library(ggtern)
HFDcolor="#E69F00"
LFDcolor="#0072B2"
sessionInfo()
```

```
## R version 3.5.0 (2018-04-23)
## Platform: x86_64-apple-darwin15.6.0 (64-bit)
## Running under: macOS  10.14
## 
## Matrix products: default
## BLAS: /Library/Frameworks/R.framework/Versions/3.5/Resources/lib/libRblas.0.dylib
## LAPACK: /Library/Frameworks/R.framework/Versions/3.5/Resources/lib/libRlapack.dylib
## 
## locale:
## [1] en_US.UTF-8/en_US.UTF-8/en_US.UTF-8/C/en_US.UTF-8/en_US.UTF-8
## 
## attached base packages:
## [1] stats     graphics  grDevices utils     datasets  methods   base     
## 
## other attached packages:
##  [1] ggtern_3.0.0    MicrobeR_0.3.1  readxl_1.1.0    forcats_0.3.0  
##  [5] stringr_1.3.1   dplyr_0.7.5     purrr_0.2.5     readr_1.1.1    
##  [9] tidyr_0.8.1     tibble_1.4.2    ggplot2_3.0.0   tidyverse_1.2.1
## 
## loaded via a namespace (and not attached):
##   [1] colorspace_1.3-2    ggtree_1.12.0       philr_1.6.0        
##   [4] rprojroot_1.3-2     XVector_0.20.0      rstudioapi_0.7     
##   [7] DT_0.4              bit64_0.9-7         lubridate_1.7.4    
##  [10] xml2_1.2.0          codetools_0.2-15    splines_3.5.0      
##  [13] mnormt_1.5-5        robustbase_0.93-2   knitr_1.20         
##  [16] ade4_1.7-11         jsonlite_1.5        phyloseq_1.24.0    
##  [19] broom_0.4.4         cluster_2.0.7-1     latex2exp_0.4.0    
##  [22] compiler_3.5.0      httr_1.3.1          rvcheck_0.1.0      
##  [25] backports_1.1.2     assertthat_0.2.0    Matrix_1.2-14      
##  [28] lazyeval_0.2.1      cli_1.0.0           htmltools_0.3.6    
##  [31] tools_3.5.0         bindrcpp_0.2.2      igraph_1.2.1       
##  [34] gtable_0.2.0        glue_1.2.0          reshape2_1.4.3     
##  [37] fastmatch_1.1-0     Rcpp_0.12.19        Biobase_2.40.0     
##  [40] cellranger_1.1.0    Biostrings_2.48.0   zCompositions_1.1.1
##  [43] multtest_2.36.0     ape_5.2             nlme_3.1-137       
##  [46] DECIPHER_2.8.1      iterators_1.0.9     psych_1.8.4        
##  [49] tensorA_0.36.1      proto_1.0.0         rvest_0.3.2        
##  [52] phangorn_2.4.0      DEoptimR_1.0-8      zlibbioc_1.26.0    
##  [55] MASS_7.3-50         scales_0.5.0        hms_0.4.2          
##  [58] parallel_3.5.0      biomformat_1.8.0    rhdf5_2.24.0       
##  [61] yaml_2.2.0          memoise_1.1.0       gridExtra_2.3      
##  [64] NADA_1.6-1          stringi_1.2.3       RSQLite_2.1.1      
##  [67] S4Vectors_0.18.3    foreach_1.4.4       energy_1.7-5       
##  [70] tidytree_0.1.9      permute_0.9-4       BiocGenerics_0.26.0
##  [73] boot_1.3-20         truncnorm_1.0-8     rlang_0.2.1        
##  [76] pkgconfig_2.0.1     compositions_1.40-2 evaluate_0.10.1    
##  [79] lattice_0.20-35     Rhdf5lib_1.2.1      bindr_0.1.1        
##  [82] treeio_1.4.1        htmlwidgets_1.2     bit_1.1-14         
##  [85] tidyselect_0.2.4    plyr_1.8.4          magrittr_1.5       
##  [88] R6_2.2.2            IRanges_2.14.10     picante_1.7        
##  [91] DBI_1.0.0           pillar_1.2.3        haven_1.1.1        
##  [94] foreign_0.8-70      withr_2.1.2         mgcv_1.8-24        
##  [97] survival_2.42-3     bayesm_3.1-0.1      modelr_0.1.2       
## [100] crayon_1.3.4        plotly_4.7.1        rmarkdown_1.10     
## [103] grid_3.5.0          data.table_1.11.4   blob_1.1.1         
## [106] vegan_2.5-2         digest_0.6.15       stats4_3.5.0       
## [109] munsell_0.5.0       viridisLite_0.3.0   quadprog_1.5-5
```

### 3 Metadata

Metadata is stored in separate files on a per-study basis and all data is available from either the SRA or MG-RAST. The first step is to collate the metadata from the separate xlsx spreadsheets. The collated version is displayed below.

```
metadata<-
  list.files("/Volumes/turnbaughlab/qb3share/jbisanz/HFD_metastudy/metadata/", full.names = TRUE) %>%
  grep(".xlsx", ., value=TRUE) %>%
  lapply(., function(x) read_excel(x, skip=2, na=c("NA",""))) %>%
  do.call(bind_rows, .)
MicrobeR::Nice.Table(metadata)
```

#### 3.1 Per study summary

```
metadata %>%
  group_by(StudyID, PMID, Database) %>%
  summarize(`# Samples`=length(DB_Sample),
            `# LFD`=sum(Diet_Classification=="LFD"),
            `# HFD`=sum(Diet_Classification=="HFD"),
            Diets=paste(unique(Diet_Name), collapse=", "),
            Species=paste(unique(Species), collapse=", "),
            Background=paste(unique(Background), collapse=", "),
            Genotypes=paste(unique(Genotype), collapse=", "),
            `Author specified strains`=paste(unique(Author_Specified_Strain), collapse=", "),
            `Colonization States`=paste(unique(Colonization), collapse=", "),
            Vendors=paste(unique(Vendor), collapse=", "),
            `Sequencing Platform`=paste(unique(Sequencing_Platform), collapse=", "),
            `Sequencing Strategy`=paste(unique(Sequencing_Strategy), collapse=", "),
            `Variable regions covered`=paste(unique(Variable_Region), collapse=", ")
            ) %>%
  MicrobeR::Nice.Table()
```

#### 3.2 Correlation of other variables with StudyID

```
metadata2<-metadata %>%
  filter(!StudyID %in% c("David 2014","Wu 2011")) %>%
  mutate(Vendor=case_when(
    grepl("Taconic", Vendor) ~ "Taconic",
    grepl("CharlesRiver", Vendor) ~ "Charles River",
    TRUE~Vendor
  )) %>%
  mutate(Diet_Name=if_else(StudyID=="Ruan 2016", "Diet Unknown", Diet_Name))

tmm<-
metadata2 %>%
  select(StudyID, Diet_Name, Species, Background, Vendor, Sequencing_Platform, Sequencing_Strategy, Variable_Region, Extraction_Method, Mechanical_Lysis) %>%
  distinct() %>%
  gather(-StudyID, key=Variable, value=value) %>%
  mutate(tm=1) %>%
  reshape2::dcast(StudyID+Variable~value, value.var="tm", fill=0, fun.aggregate=function(x){return(1)}) %>%
  select(-Variable) %>%
  group_by(StudyID) %>%
  summarise_all(function(x){if_else(sum(x)>0, 1, 0)})

pdf("figures/tree.pdf", height=5, width=8)
tmm %>% column_to_rownames("StudyID") %>% dist(., "euclidean") %>% hclust(., method="average") %>% plot()
dev.off()
```

```
## quartz_off_screen 
##                 2
```

```
ord<-tmm %>% column_to_rownames("StudyID") %>% dist(., "euclidean") %>% hclust(., method="average")
ord2<-tmm %>% column_to_rownames("StudyID") %>% t() %>% dist(., "euclidean") %>% hclust(., method="average")


tmm %>%
  gather(-StudyID, key=Variable, value=Present) %>%
  left_join(
    metadata2 %>%
    select(StudyID, Diet_Name, Species, Background, Vendor, Sequencing_Platform, Sequencing_Strategy, Variable_Region, Extraction_Method, Mechanical_Lysis) %>%
    distinct() %>%
    gather(-StudyID, key=Variable, value=value) %>%
      select(Metric=Variable, Variable=value)
  ) %>%
  mutate(StudyID=factor(StudyID, levels=ord$labels[ord$order])) %>%
  mutate(Variable=factor(Variable, levels=ord2$labels[ord2$order])) %>%
  mutate(Metric=factor(Metric, levels=c("Sequencing_Platform", "Sequencing_Strategy","Variable_Region", "Species","Background","Vendor","Extraction_Method","Mechanical_Lysis", "Diet_Name"))) %>%
  #mutate(Metric="junk") %>%
  ggplot(aes(x=Variable, y=StudyID, fill=Present)) +
  geom_tile(color="grey50") +
  facet_grid(~Metric, scales="free_x", space="free") +
  theme_MicrobeR() +
  theme(axis.text.x = element_text(angle=45, hjust=1)) +
  coord_cartesian(expand=F) +
  scale_fill_gradient2(low="grey50", high="indianred") +
  theme(legend.position="none")
```

```
## Joining, by = "Variable"
```

```
ggsave("figures/metadata_hm.pdf", device="pdf", height=4, width=13)
```

#### 3.3 Ternary diagram of diet composition

```
metadata %>%
  filter(Species!="Homo sapiens") %>%
  select(StudyID, Diet_Classification, P_Protein, P_Fat, P_Carb) %>%
  distinct() %>%
  filter(!is.na(P_Protein)) %>%
  ggtern(aes(x=P_Protein, y=P_Fat, z=P_Carb, color=Diet_Classification, group=StudyID)) +
  geom_line(alpha=0.2, color="grey50") +
  geom_point(alpha=0.8, shape=16) +
  #geom_text(aes(label=StudyID)) +
  scale_color_manual(values = c(HFDcolor,LFDcolor)) +
  theme_bw() +
  theme(panel.grid.minor=element_blank(), panel.grid.major = element_blank()) +
  theme(legend.position="none")
```

```
ggsave("figures/diet_comp.pdf", device="pdf", height=3, width=3, useDingbats=FALSE)
```

#### 3.4 Summary of sample characteristics

```
metadata %>%
  filter(Species!="Homo sapiens") %>%
  select(Background, Sequencing_Platform, Sequencing_Strategy,Variable_Region) %>%
  gather(key=Metric) %>%
  group_by(Metric, value) %>%
  summarize(n=length(value)) %>%
  ungroup() %>% 
  mutate(Metric=factor(Metric, levels=c("Sequencing_Strategy", "Sequencing_Platform", "Variable_Region", "Background"))) %>%
  arrange(desc(n)) %>%
  mutate(value=factor(value, levels=unique(value))) %>%
  ggplot(aes(x=value, y=n)) +
  geom_bar(stat="identity", color="black", fill=LFDcolor) +
  facet_grid(~Metric, scales="free", space="free") +
  theme_MicrobeR() +
  theme(axis.text.x=element_text(angle=45, hjust=1)) +
  ylab("# Samples") +
  xlab("") +
  ggtitle("Sample information") +
  coord_cartesian(ylim=c(0,1000), expand=FALSE) +
  theme(panel.border = element_blank(), axis.line = element_line())
```

```
ggsave("figures/sample_meta.pdf", device="pdf", height=2.5, width=4, useDingbats=FALSE)
```

---

### 4 Downloading Raw Data

There are two different strategies depending on if the data is derived from the SRA or MGRAST. SRA data is downloaded via the SRAdb package for R to build a download list followed by a system call using wget. In this case we are using FTP rather than aspera due to firewall issues but other users would likely prefer the speed of using aspera. MG-RAST data is downloaded via curl using a system call. A different user will need to supply their own authentication token for MGRAST.

#### 4.1 MGRAST

For data from MGRAST, the original user’s uploaded file is being downloaded. Note that many files from MGRAST do not have quality encoding available. Where this is true, it has been assumed the original authors carried out appropriate quality control and filtering. For compatibility, these FASTA files are being converted to FASTQ files with all bases given a quality score of 30 (?). In some cases it would appear that data has already been filtered and trimmed but complete data about the preprocessing is not available via MGRAST.

```
library(tidyverse)
library(readxl)

metadata<-
  list.files("/labmainshare/qb3share/jbisanz/HFD_metastudy/metadata/", full.names = TRUE) %>%
  grep(".xlsx", ., value=TRUE) %>%
  lapply(., function(x) read_excel(x, skip=2, na=c("NA",""))) %>%
  do.call(bind_rows, .)
  
mgsamples<-
  metadata %>%
  filter(Database=="MGRAST")

for(i in 1:nrow(mgsamples)){
  message(mgsamples[i,]$ManuscriptID,": ", round(100*(i/nrow(mgsamples)), 3), "%")
  dir.create(paste0("Samples/", mgsamples[i,]$DB_Sample))
  system(
    paste0(
      "curl -s -X GET -H \"auth: ",
      "yourtokenhere", #your token here, this will be expired by the time another user attempts to run this, get from MG RAST profile
      "\" http://api.metagenomics.anl.gov/1/download/\"",
      mgsamples[i,]$DB_Sample,
      "\"?file=050.1 > ",
      "Samples/", mgsamples[i,]$DB_Sample,"/",mgsamples[i,]$DB_Sample,".fastq"
      )
    )
  
  system(
    paste0(
      "pigz ", "Samples/", mgsamples[i,]$DB_Sample,"/",mgsamples[i,]$DB_Sample,".fastq"
    )
  )
}


# last but not least, not all files were fastq, where this is present, search the first line and and if a fasta create a fastq using a quality score of 30 (?)
library(ShortRead)
for(i in 1:nrow(mgsamples)){
    message(mgsamples[i,]$ManuscriptID," ",mgsamples[i,]$DB_Sample,": ", round(100*(i/nrow(mgsamples)), 3), "%")
  infile<-paste0("Samples/", mgsamples[i,]$DB_Sample,"/",mgsamples[i,]$DB_Sample,".fastq.gz")
  FileType=if_else(system(paste0("zcat ", infile, " | head -n 1"), intern = TRUE) %>% substr(., 1,1)=="@", "fastq","fasta")
  if(FileType=="fasta"){
    message(mgsamples[i,]$ManuscriptID,": renaming ")
    file.rename(infile, gsub("fastq","fasta", infile))
    message(mgsamples[i,]$ManuscriptID,": reading ")
    tmp<-readFasta(gsub("fastq","fasta", infile))
    if(length(tmp)<5e5){     #process large files in multiple chunks
      message(mgsamples[i,]$ManuscriptID,": generating pseudo quality scores for less than 5e5 reads ")
      quals<-sapply(tmp@sread@ranges@width, function(x) rep("?", x) %>% paste(., collapse=""))
      message(mgsamples[i,]$ManuscriptID,": building fastq ")
      fastq<-ShortReadQ(sread=tmp@sread, quality=BStringSet(quals), id = tmp@id)
      message(mgsamples[i,]$ManuscriptID,": writting fastq ")
      writeFastq(fastq, infile)
    } else{
      chunksize=5e5
      for (chunkstart in seq(1, length(tmp), chunksize)) {
        chunkend<-chunkstart+(chunksize-1)
        if((chunkstart+(chunksize-1))>length(tmp)){chunkend=length(tmp)}
          message(mgsamples[i,]$ManuscriptID,": generating scores and writting fastq for ", round(chunkend/length(tmp)*100, 1), "% of reads")
          quals<-sapply(tmp@sread[chunkstart:chunkend]@ranges@width, function(x) rep("?", x) %>% paste(., collapse=""))
          fastq<-ShortReadQ(sread=tmp[chunkstart:chunkend]@sread, quality=BStringSet(quals), id = tmp[chunkstart:chunkend]@id)
          writeFastq(fastq, gsub("\\.gz","", infile), mode="a", compress=FALSE)
        }
      system(paste("pigz ", gsub("\\.gz","", infile)))
    }

      system(paste0("echo pseudoquality >> Samples/", mgsamples[i,]$DB_Sample, "/readme"))
    rm(fastq, tmp, quals)
    gc()
  }
}
```

---

#### 4.2 SRA

For data deposited in the Sequence Read Archive and/or European Nucleotide Archive, the sample’s accession (SRS######/ERS#######) is being queried to pull all runs belonging to the sample using the the ENA API. This is an alternative to using the SRAdb package as I found that a number of the download links were broken or did not accurately reflect PE/SE.

```
library(tidyverse)
library(readxl)

metadata<-
  list.files("/Volumes/turnbaughlab/qb3share/jbisanz/HFD_metastudy/metadata/", full.names = TRUE) %>%
  grep(".xlsx", ., value=TRUE) %>%
  lapply(., function(x) read_excel(x, skip=2, na=c("NA",""))) %>%
  do.call(bind_rows, .) %>%
  filter(Database %in% c("SRA","ENA"))

dir.create("dbs/sradownloads")

for(i in metadata$DB_Sample){
  message(i)
  query=paste0("wget -O dbs/SRAdownloads/", i,".txt \"http://www.ebi.ac.uk/ena/data/warehouse/search?query=\"secondary_sample_accession=",i,"\"&result=read_run&fields=secondary_sample_accession,fastq_ftp,secondary_study_accession&display=report&download=txt\"")
  system(query)
}

downloads<-
  list.files("dbs/SRAdownloads/", pattern="txt$", full.names=T) %>%
  lapply(., read_tsv) %>%
  do.call(bind_rows, .)

downloads<- downloads %>%
  select(DB_Study=secondary_study_accession, DB_Sample=secondary_sample_accession, DB_Run=run_accession, fastq_ftp) %>%
  filter(!is.na(DB_Sample)) %>%
  filter(!DB_Sample %in% subset(metadata, StudyID %in% c("Goodman 2011", "Wu 2011"))$DB_Sample)

write_tsv(downloads,"dbs/SRAdownloads_18dec2018.txt")
```

Now will download the SRA files. A set of samples from Goodman 2011 (n=14) and Wu 2011 (n=10) are difficult to FTP as they were run sequenced across multiple runs which themselves were uploaded in a non-demultiplexed reads which interfers with their downloading in demultiplexed state. This limited number of samples was manually downloaded using the Sequence Read Archive run browser and placed into the `Samples/` Directory with the rest of the reads.

```
library(tidyverse)

downloads<-read_tsv("dbs/SRAdownloads_18dec2018.txt")

for(i in 1:nrow(downloads)){
  sample=downloads$DB_Sample[i]
  run=downloads$DB_Run[i]
  ftp=gsub(";$","", downloads$fastq_ftp[i]) %>% str_split(., pattern=";") %>% unlist()
  
  message("Downloading ", sample, " ", run, ": ", paste(ftp, collapse=" "))
  
  dir.create(paste0("Samples/",sample,"/"))
  
  for(f in ftp){system(paste0("wget -N --quiet --directory-prefix=Samples/",sample," ", f))}
}
```

Will now do a quick check for missing files/failed downloads.

```
metadata<-
  list.files("/Volumes/turnbaughlab/qb3share/jbisanz/HFD_metastudy/metadata/", full.names = TRUE) %>%
  grep(".xlsx", ., value=TRUE) %>%
  lapply(., function(x) read_excel(x, skip=2, na=c("NA",""))) %>%
  do.call(bind_rows, .)

Samps<-list.files("/Volumes/turnbaughlab/qb3share/jbisanz/HFD_metastudy/Samples/")

print(paste("The following samples are missing:", paste(Samps[!Samps %in% metadata$DB_Sample], collapse=", "), "."))
```

```
## [1] "The following samples are missing:  ."
```

```
downloads<-
  read_tsv("/Volumes/turnbaughlab/qb3share/jbisanz/HFD_metastudy/dbs/SRAdownloads_18dec2018.txt") %>%
  separate(fastq_ftp, c("f","r"), sep=";") %>%
  select(f, r) %>%
  gather() %>%
  filter(!is.na(value)) %>%
  mutate(value=basename(value)) %>%
  pull(value) %>% as.vector()
```

```
## Parsed with column specification:
## cols(
##   DB_Study = col_character(),
##   DB_Sample = col_character(),
##   DB_Run = col_character(),
##   fastq_ftp = col_character()
## )
```

```
print(
  paste("The following runs were not downloaded successfully:",
        paste(
          downloads[!downloads %in% (list.files("/Volumes/turnbaughlab/qb3share/jbisanz/HFD_metastudy/Samples/", recursive = T, pattern="fastq\\.gz") %>% basename())]
          , collapse=", ")
        ,".")
)
```

```
## [1] "The following runs were not downloaded successfully:  ."
```

### 5 Identifying Lactococcus OTUs

For reasons that will become clear later during analysis, we wish to create a version of the data which does not contain lactococci. To accomplish this, we will find the most recent common ancestor of all named Lactococcus species in the GG tree, and remove these from the OTU table.

```
library(ape)
lactoOTUs<-read_tsv("/Volumes/turnbaughlab/qb3share/jbisanz/HFD_metastudy/dbs/gg_13_8_otus/taxonomy/97_otu_taxonomy.txt", col_names=F) %>% filter(grepl("Lactococcus", X2)) %>% pull(X1)
print(paste("The following", length(lactoOTUs),"OTUs are named as Lactococcus in the GG taxonomy:", paste(lactoOTUs, collapse=", "),"."))
tree<-read.tree("/Volumes/turnbaughlab/qb3share/jbisanz/HFD_metastudy/dbs/gg_13_8_otus/trees/97_otus.tree")
lactoMRCA<-getMRCA(phy=tree, tip=as.character(lactoOTUs))
print(paste("The MRCA for Lactococcus is", lactoMRCA,"."))
lactoOTUs<-extract.clade(tree, lactoMRCA)$tip.label
print(paste("The following", length(lactoOTUs)," OTUs will be targetted for removal:", paste(lactoOTUs, collapse=", "),"."))

write_tsv(tibble(OTU=lactoOTUs), "/Volumes/turnbaughlab/qb3share/jbisanz/HFD_metastudy/dbs/lacto.otus", col_names=F)
```

---

### 6 OTU picking

All processing and OTU picking is handled by one central SGE array job as below using USEARCH 10.0.240 (university compute cluster version),VSEARCH 2.4.4, and SortMeRNA 2.1b, and PICRUSt 1.0.0dev. Due to inconsistencies such as non-overlapping amplicon sequencing and metagenomic sequencing. Only the forward read of any given run is being used. The following pipeline is used below:

1. Build a local node-access only directory and copy reads to this directory.
2. Overlap reads (if possible), if >50% can be overlapped, will use overlapped reads, if not, use only forward read.
3. Filter reads to truncate after a quality score of 20, allowing no Ns, a max of 2 expected errors, and a minimum read length of 60bp.
4. Filter reads to only those mapping to 16S rDNA using SortMeRNA against a 90% clustered SILVA bacterial database (silva-bac-16s-id90).
5. Do closed reference OTU picking against Green Genes 13\_8.
6. Create a second version of OTU level without Lactococcus OTUs identified above.
7. Run PICRUSt to generate a correspoding table of kegg orthologs (gene content) on both versions of OTU tables.

```
#!/bin/bash
#$ -S /bin/bash
#$ -o OTUmaster.log
#$ -e OTUmaster.err
#$ -cwd
#$ -r y
#$ -j y
#$ -l mem_free=64G
#$ -l arch=linux-x64
#$ -l scratch=10G
#$ -l h_rt=144:0:0
#$ -t 1-1086

#1-1086
source /turnbaugh/qb3share/shared_resources/export_python_2.7.14.sh 
export PICRUST=/turnbaugh/qb3share/shared_resources/sftwrshare/picrust/scripts/
export PATH=$PATH:/turnbaugh/qb3share/shared_resources/sftwrshare/sortmerna-2.1b/

mkdir -p OTUs
mkdir -p nol_OTUs
mkdir -p PICRUSt
mkdir -p nol_PICRUSt
mkdir -p logs

SAMPLE=$( ls -1 Samples | sed "${SGE_TASK_ID}q;d" )

echo $(date) $SAMPLE running on $(hostname) for Job $JOB_ID and Task $SGE_TASK_ID

exec >/turnbaugh/qb3share/jbisanz/HFD_metastudy/logs/${SAMPLE}_${JOB_ID}_${SGE_TASK_ID}.log 2>/turnbaugh/qb3share/jbisanz/HFD_metastudy/logs/${SAMPLE}_${JOB_ID}_${SGE_TASK_ID}.errors

TMDIR=/scratch/${SAMPLE}_${JOB_ID}
echo $(date) $SAMPLE running on $(hostname) working at $TMDIR
mkdir $TMDIR
cd $TMDIR

cp /turnbaugh/qb3share/jbisanz/HFD_metastudy/Samples/${SAMPLE}/*fastq.gz .

ls -lhS .

RUNS=$( ls -1 *fastq.gz | sed 's/_[12]\.fastq\.gz//g' | sed 's/\.fastq\.gz//g' | sort | uniq )

echo $(date) There are $( ls -1 *fastq.gz | sed 's/_[12]\.fastq\.gz//g' | sed 's/\.fastq\.gz//g' | sort | uniq | wc -l ) echo Runs for this sample: $RUNS

read -a RA <<< $RUNS
for RUN in "${RA[@]}";do

  TYPE=$( ls -1 $RUN* | wc -l )
  
  if [ $TYPE == 1 ]; then
    echo Running as single end with $RUN
    pigz -d ${RUN}.fastq.gz
    #QC
    vsearch_v2.4.4 \
    --fastq_filter ${RUN}.fastq \
    --fastaout ${RUN}.filtered.fasta \
    --fastq_truncqual 20 \
    --fastq_maxns 0 \
    --fastq_minlen 60 \
    --fastq_maxee 2 \
    --threads 1
    
  elif [ $TYPE == 2 ]; then
  echo Running as paired end with $RUN
    pigz -d ${RUN}_*.fastq.gz
    
    vsearch_v2.4.4 \
    --fastq_mergepairs ${RUN}_1.fastq \
    --reverse ${RUN}_2.fastq \
    --threads 6 \
    --fastqout ${RUN}_ol.fastq \
    &> ${RUN}.olstats
    
    cat ${RUN}.olstats >> /turnbaugh/qb3share/jbisanz/HFD2018/logs/${SAMPLE}_${JOB_ID}_${SGE_TASK_ID}.log
    
    OVERLAPPED=$( grep 'Merged' ${RUN}.olstats | sed 's/.*(//g' | sed 's/%)//g' )
    
    if (( $(echo "$OVERLAPPED > 50" | bc -l) )); then
      echo Using the overlapped reads as $OVERLAPPED were merged
            mv ${RUN}_ol.fastq ${RUN}_tofilt.fastq
    else
      echo NOT USING the overlapped reads as $OVERLAPPED were merged
      mv ${RUN}_1.fastq ${RUN}_tofilt.fastq
    fi
    
    vsearch_v2.4.4 \
    --fastq_filter ${RUN}_tofilt.fastq \
    --fastaout ${RUN}.filtered.fasta \
    --fastq_truncqual 20 \
    --fastq_maxns 0 \
    --fastq_minlen 60 \
    --fastq_maxee 2 \
    --threads 1
  fi
  
  #Identify rRNA reads
  sortmerna \
  --ref /turnbaugh/qb3share/jbisanz/HFD_metastudy/dbs/sortmerna-2.1b/rRNA_databases/silva-bac-16s-id90.fasta,/turnbaugh/qb3share/jbisanz/HFD_metastudy/dbs/sortmerna-2.1b/index/silva-bac-16s-id90 \
  -v \
  --reads ${RUN}.filtered.fasta \
  -a 1 \
  --fastx \
  --aligned ${RUN}.aligned
    
  #closed ref picking
  usearch10.0.240 \
    -fastx_relabel ${RUN}.aligned.fasta \
    -prefix ${RUN}.read_ \
    -fastaout ${RUN}.relab.fasta
  
  usearch10.0.240 \
  -closed_ref ${RUN}.relab.fasta \
  -otutabout ${RUN}.otus \
  -db /turnbaugh/qb3share/jbisanz/HFD_metastudy/dbs/gg_13_8_otus/rep_set/97_otus.fasta \
  -strand both \
  -threads 1 \
  -id 0.97
  
  # Create a Lacto-free version of the OTU table
  cp ${RUN}.otus toclean.tsv
  module load MRO
  Rscript -e 'suppressMessages(library(tidyverse));lotu<-read_tsv("/turnbaugh/qb3share/jbisanz/HFD_metastudy/dbs/lacto.otus", col_names=F);read_tsv("toclean.tsv", comment="") %>% filter(!`#OTU ID` %in% lotu$X1) %>% write_tsv("cleaned.tsv")'
  cp cleaned.tsv ${RUN}.noL_otus
  
  cp ${RUN}.otus /turnbaugh/qb3share/jbisanz/HFD_metastudy/OTUs/${SAMPLE}_${RUN}.otus 
  cp ${RUN}.noL_otus /turnbaugh/qb3share/jbisanz/HFD_metastudy/nol_OTUs/${SAMPLE}_${RUN}.otus_nol
  
  #####PICRUST
  biom convert --to-json -i ${RUN}.otus -o ${RUN}.biom
  $PICRUST/normalize_by_copy_number.py -i ${RUN}.biom -o ${RUN}.norm.biom
  $PICRUST/predict_metagenomes.py -f \
  -i ${RUN}.norm.biom \
  -o ${RUN}.picrust
  
  cp ${RUN}.picrust /turnbaugh/qb3share/jbisanz/HFD_metastudy/PICRUSt/${SAMPLE}_${RUN}.picrust
  
  biom convert --to-json -i ${RUN}.noL_otus -o ${RUN}.biom
  $PICRUST/normalize_by_copy_number.py -i ${RUN}.biom -o ${RUN}.norm.biom
  $PICRUST/predict_metagenomes.py -f \
  -i ${RUN}.norm.biom \
  -o ${RUN}.noL_picrust
  
  cp ${RUN}.noL_picrust /turnbaugh/qb3share/jbisanz/HFD_metastudy/nol_PICRUSt/${SAMPLE}_${RUN}.picrust_nol
  
done

ls -lh
cd /turnbaugh/qb3share/jbisanz/HFD_metastudy
rm -r $TMDIR
echo COMPLETE ..... $(date) $SAMPLE Complete on $(hostname) for Job $JOB_ID and Task $SGE_TASK_ID
echo COMPLETE ..... $(date) $SAMPLE Complete on $(hostname) for Job $JOB_ID and Task $SGE_TASK_ID >> OTUmaster.log
```

---

### 7 Collating Tables

As tables were formed on a per-sample basis. We need to merge them together and then aggregate into sample. This is with the exception of samples from Voigt 2014 where each run represents a separate animal.

```
vgt<-read_excel("/Volumes/turnbaughlab/qb3share/jbisanz/HFD_metastudy/metadata/Voigt 2014.xlsx", skip=2)
```

#### 7.1 OTUs

```
OTUs<-
list.files("/Volumes/turnbaughlab/qb3share/jbisanz/HFD_metastudy/OTUs/", full.names = T, pattern="\\.otus") %>%
  lapply(., function(x) suppressMessages(read_tsv(x, skip=1, col_names=c("OTU", "Reads"))) %>% mutate(Sample=gsub("\\.otus","", basename(x)))) %>%
  do.call(bind_rows, .) %>%
  separate(Sample, c("Sample","Run"), sep="_") %>%
  mutate(Sample=case_when(
    Sample %in% vgt$DB_Sample ~ Run,
    TRUE ~ Sample
  )) %>%
  group_by(OTU, Sample) %>%
  summarize(Reads=sum(Reads)) %>%
  ungroup() %>%
  spread(key=Sample, value=Reads, fill=0)

write_tsv(OTUs, "/Volumes/turnbaughlab/qb3share/jbisanz/HFD_metastudy/CollatedData/OTUs.tsv")
```

#### 7.2 OTUs without lactococci

```
OTUs<-
list.files("/Volumes/turnbaughlab/qb3share/jbisanz/HFD_metastudy/nol_OTUs/", full.names = T, pattern="\\.otus_nol") %>%
  lapply(., function(x) suppressMessages(read_tsv(x, skip=1, col_names=c("OTU", "Reads"))) %>% mutate(Sample=gsub("\\.otus_nol","", basename(x)))) %>%
  do.call(bind_rows, .) %>%
  separate(Sample, c("Sample","Run"), sep="_") %>%
    mutate(Sample=case_when(
    Sample %in% vgt$DB_Sample ~ Run,
    TRUE ~ Sample
  )) %>%
  group_by(OTU, Sample) %>%
  summarize(Reads=sum(Reads)) %>%
  ungroup() %>%
  spread(key=Sample, value=Reads, fill=0)

write_tsv(OTUs, "/Volumes/turnbaughlab/qb3share/jbisanz/HFD_metastudy/CollatedData/OTUs_nol.tsv")
```

#### 7.3 PICRUSt

```
PICRUST<-
list.files("/Volumes/turnbaughlab/qb3share/jbisanz/HFD_metastudy/PICRUSt/", full.names = T, pattern="\\.picrust") %>%
  lapply(., function(x) suppressMessages(read_tsv(x, skip=2, col_names=c("OTU", "Reads","Junk"))) %>% select(-Junk) %>% mutate(Sample=gsub("\\.picrust","", basename(x)))) %>%
  do.call(bind_rows, .) %>%
  separate(Sample, c("Sample","Run"), sep="_") %>%
    mutate(Sample=case_when(
    Sample %in% vgt$DB_Sample ~ Run,
    TRUE ~ Sample
  )) %>%
  group_by(OTU, Sample) %>%
  summarize(Reads=sum(Reads)) %>%
  ungroup() %>%
  spread(key=Sample, value=Reads, fill=0)

write_tsv(PICRUST, "/Volumes/turnbaughlab/qb3share/jbisanz/HFD_metastudy/CollatedData/PICRUSt.tsv")
```

#### 7.4 PICRUSt without lactococci

```
PICRUST<-
list.files("/Volumes/turnbaughlab/qb3share/jbisanz/HFD_metastudy/nol_PICRUSt/", full.names = T, pattern="\\.picrust_nol") %>%
  lapply(., function(x) suppressMessages(read_tsv(x, skip=1, col_names=c("OTU", "Reads","Junk"))) %>% select(-Junk) %>% mutate(Sample=gsub("\\.picrust_nol","", basename(x)))) %>%
  do.call(bind_rows, .) %>%
  separate(Sample, c("Sample","Run"), sep="_") %>%
    mutate(Sample=case_when(
    Sample %in% vgt$DB_Sample ~ Run,
    TRUE ~ Sample
  )) %>%
  group_by(OTU, Sample) %>%
  summarize(Reads=sum(Reads)) %>%
  ungroup() %>%
  spread(key=Sample, value=Reads, fill=0)

write_tsv(PICRUST, "/Volumes/turnbaughlab/qb3share/jbisanz/HFD_metastudy/CollatedData/PICRUSt_nol.tsv")
```

#### 7.5 Metadata

Finally, will make an updated version of the aggregated metadata.

```
vgt<-read_excel("/Volumes/turnbaughlab/qb3share/jbisanz/HFD_metastudy/metadata/Voigt 2014.xlsx", skip=2) %>%
  mutate(Supplemental_Information=paste(DB_Sample, Supplemental_Information)) %>%
  mutate(DB_Sample=gsub("..+;","", Supplemental_Information) %>% gsub(" ","", .))

vgt<-
bind_rows(
lapply(1:10, function(x) subset(vgt, Diet_Classification=="LFD") %>% mutate(DB_Sample=str_split(DB_Sample, ",")[[1]][x])) %>% do.call(bind_rows, .),
lapply(1:8, function(x) subset(vgt, Diet_Classification=="HFD") %>% mutate(DB_Sample=str_split(DB_Sample, ",")[[1]][x])) %>% do.call(bind_rows, .)
)


metadata<-
  list.files("/Volumes/turnbaughlab/qb3share/jbisanz/HFD_metastudy/metadata/", full.names = TRUE, pattern="xlsx") %>%
  lapply(., function(x) read_excel(x, skip=2, na=c("NA",""))) %>%
  do.call(bind_rows, .) %>%
  filter(StudyID!="Voigt 2014") %>%
  bind_rows(., vgt)

write_tsv(metadata, "/Volumes/turnbaughlab/qb3share/jbisanz/HFD_metastudy/CollatedData/metadata.tsv")
```

---

# 8 QC

#### 8.1 Check for failed OTU picking

```
metadata<-read_tsv("/Volumes/turnbaughlab/qb3share/jbisanz/HFD_metastudy/CollatedData/metadata.tsv")
```

```
## Parsed with column specification:
## cols(
##   .default = col_character(),
##   PMID = col_integer(),
##   P_Protein = col_double(),
##   P_Fat = col_double(),
##   P_Carb = col_double()
## )
```

```
## See spec(...) for full column specifications.
```

```
otus<-read.table("/Volumes/turnbaughlab/qb3share/jbisanz/HFD_metastudy/CollatedData/OTUs.tsv", sep="\t", row.names=1, header=T) %>% colnames(.)
otus_nol<-read.table("/Volumes/turnbaughlab/qb3share/jbisanz/HFD_metastudy/CollatedData/OTUs_nol.tsv", sep="\t", row.names=1, header=T) %>% colnames(.)
picrust<-read.table("/Volumes/turnbaughlab/qb3share/jbisanz/HFD_metastudy/CollatedData/PICRUSt.tsv", sep="\t", row.names=1, header=T) %>% colnames(.)
picrust_nol<-read.table("/Volumes/turnbaughlab/qb3share/jbisanz/HFD_metastudy/CollatedData/PICRUSt_nol.tsv", sep="\t", row.names=1, header=T) %>% colnames(.)

metadata %>% filter(!DB_Sample %in% otus)
```

```
## # A tibble: 1 x 27
##   StudyID ManuscriptID Database DB_Study DB_Sample   PMID Diet_Classifica…
##   <chr>   <chr>        <chr>    <chr>    <chr>      <int> <chr>           
## 1 Moya-P… SD2_2        ENA      ERP0089… ERS622305 2.62e7 LFD             
## # ... with 20 more variables: Diet_Name <chr>, P_Protein <dbl>,
## #   P_Fat <dbl>, P_Carb <dbl>, Species <chr>, Background <chr>,
## #   Genotype <chr>, Author_Specified_Strain <chr>, Colonization <chr>,
## #   Donor_Info <chr>, Vendor <chr>, Vendor_Location <chr>,
## #   Sequencing_Platform <chr>, Sequencing_Strategy <chr>,
## #   Variable_Region <chr>, Read_Type <chr>, Extraction_Method <chr>,
## #   Mechanical_Lysis <chr>, Enzymatic_Lysis <chr>,
## #   Supplemental_Information <chr>
```

```
metadata %>% filter(!DB_Sample %in% otus_nol)
```

```
## # A tibble: 1 x 27
##   StudyID ManuscriptID Database DB_Study DB_Sample   PMID Diet_Classifica…
##   <chr>   <chr>        <chr>    <chr>    <chr>      <int> <chr>           
## 1 Moya-P… SD2_2        ENA      ERP0089… ERS622305 2.62e7 LFD             
## # ... with 20 more variables: Diet_Name <chr>, P_Protein <dbl>,
## #   P_Fat <dbl>, P_Carb <dbl>, Species <chr>, Background <chr>,
## #   Genotype <chr>, Author_Specified_Strain <chr>, Colonization <chr>,
## #   Donor_Info <chr>, Vendor <chr>, Vendor_Location <chr>,
## #   Sequencing_Platform <chr>, Sequencing_Strategy <chr>,
## #   Variable_Region <chr>, Read_Type <chr>, Extraction_Method <chr>,
## #   Mechanical_Lysis <chr>, Enzymatic_Lysis <chr>,
## #   Supplemental_Information <chr>
```

```
metadata %>% filter(!DB_Sample %in% picrust)
```

```
## # A tibble: 1 x 27
##   StudyID ManuscriptID Database DB_Study DB_Sample   PMID Diet_Classifica…
##   <chr>   <chr>        <chr>    <chr>    <chr>      <int> <chr>           
## 1 Moya-P… SD2_2        ENA      ERP0089… ERS622305 2.62e7 LFD             
## # ... with 20 more variables: Diet_Name <chr>, P_Protein <dbl>,
## #   P_Fat <dbl>, P_Carb <dbl>, Species <chr>, Background <chr>,
## #   Genotype <chr>, Author_Specified_Strain <chr>, Colonization <chr>,
## #   Donor_Info <chr>, Vendor <chr>, Vendor_Location <chr>,
## #   Sequencing_Platform <chr>, Sequencing_Strategy <chr>,
## #   Variable_Region <chr>, Read_Type <chr>, Extraction_Method <chr>,
## #   Mechanical_Lysis <chr>, Enzymatic_Lysis <chr>,
## #   Supplemental_Information <chr>
```

```
metadata %>% filter(!DB_Sample %in% picrust_nol)
```

```
## # A tibble: 1 x 27
##   StudyID ManuscriptID Database DB_Study DB_Sample   PMID Diet_Classifica…
##   <chr>   <chr>        <chr>    <chr>    <chr>      <int> <chr>           
## 1 Moya-P… SD2_2        ENA      ERP0089… ERS622305 2.62e7 LFD             
## # ... with 20 more variables: Diet_Name <chr>, P_Protein <dbl>,
## #   P_Fat <dbl>, P_Carb <dbl>, Species <chr>, Background <chr>,
## #   Genotype <chr>, Author_Specified_Strain <chr>, Colonization <chr>,
## #   Donor_Info <chr>, Vendor <chr>, Vendor_Location <chr>,
## #   Sequencing_Platform <chr>, Sequencing_Strategy <chr>,
## #   Variable_Region <chr>, Read_Type <chr>, Extraction_Method <chr>,
## #   Mechanical_Lysis <chr>, Enzymatic_Lysis <chr>,
## #   Supplemental_Information <chr>
```

A single sample (ERS622305) has been lost in this analysis because it appears that a fastq file is not available for this sample. We will remove this sample from further analysis.

```
metadata<-metadata %>% filter(DB_Sample!="ERS622305")
write_tsv(metadata, "/Volumes/turnbaughlab/qb3share/jbisanz/HFD_metastudy/CollatedData/metadata_filtered.tsv")
```

#### 8.2 Check read depth and observed OTUs

```
otus<-read.table("/Volumes/turnbaughlab/qb3share/jbisanz/HFD_metastudy/CollatedData/OTUs.tsv", sep="\t", row.names=1, header=T)

summary(colSums(otus))
```

```
##    Min. 1st Qu.  Median    Mean 3rd Qu.    Max. 
##     113    9258   31103   96663   80969 1425468
```

```
data.frame(Counts=colSums(otus)) %>%
  rownames_to_column("SampleID") %>%
  ggplot(aes(x=Counts)) +
  geom_freqpoly() +
  scale_x_log10() +
  theme_MicrobeR() +
  annotation_logticks(sides="b") +
  ylab("# Samples") +
  xlab("Counts") +
  ggtitle("Count distribution across all samples")
```

```
## `stat_bin()` using `bins = 30`. Pick better value with `binwidth`.
```

```
ggsave("figures/depth.pdf", device="pdf", height=3, width=3)
```

```
## `stat_bin()` using `bins = 30`. Pick better value with `binwidth`.
```

So there at least 100 reads in all samples so will not apply a minimum read count filter. Now will check for number of observed OTUs.

```
summary(vegan::specnumber(otus, MARGIN=2))
```

```
##    Min. 1st Qu.  Median    Mean 3rd Qu.    Max. 
##    12.0   351.0   663.0   987.3  1557.0  4463.0
```

```
data.frame(Counts=vegan::specnumber(otus, MARGIN=2)) %>%
  rownames_to_column("SampleID") %>%
  ggplot(aes(x=Counts)) +
  geom_freqpoly() +
  scale_x_log10() +
  theme_MicrobeR() +
  annotation_logticks(sides="b") +
  ylab("# Samples") +
  xlab("Observed OTUs") +
  ggtitle("OTU richness distribution across all samples")
```

```
## `stat_bin()` using `bins = 30`. Pick better value with `binwidth`.
```

```
ggsave("figures/richness.pdf", device="pdf", height=3, width=3)
```

```
## `stat_bin()` using `bins = 30`. Pick better value with `binwidth`.
```

All looks good so no additional QC will be done.

#### 8.3 Read depth per study

```
counts<-data.frame(Counts=colSums(otus)) %>%
  rownames_to_column("DB_Sample") %>%
  left_join(metadata)
```

```
## Joining, by = "DB_Sample"
```

```
ord<-counts %>% group_by(StudyID) %>% summarize(median=median(Counts)) %>% arrange(desc(median)) %>% pull(StudyID)

counts %>% mutate(StudyID=factor(StudyID, levels=ord)) %>%
  ggplot(aes(x=StudyID, y=Counts, fill=StudyID)) +
  geom_boxplot(outlier.size=0.1) +
  theme_MicrobeR() +
  scale_y_log10() +
  annotation_logticks(sides="l") +
  ylab("# Samples") +
  xlab("Counts") +
  ggtitle("Count distribution across all studies") +
  theme(legend.position="none") +
  theme(axis.text.x = element_text(angle=45, hjust=1)) +
  geom_hline(yintercept = 100, linetype="dashed", color="red")
```

```
ggsave("figures/depth_by_study.pdf", device="pdf", height=3, width=4, useDingbats=F)
```

### 9 Save Session

```
save.image(paste0("sessions/Session_",format(Sys.time(),"%d%b%Y_%H%M%S"),".Rdata"))
gc()
```

```
##            used  (Mb) gc trigger  (Mb) limit (Mb) max used  (Mb)
## Ncells  4655983 248.7    7296692 389.7         NA  7296692 389.7
## Vcells 24795693 189.2   86530936 660.2      16384 86528034 660.2
```

---

Return to main page
