## Extended Data File 1 for "Diet induces reproducible alterations in the mouse and human gut microbiome": breakpoints.min.js

```
/* breakpoints.js v1.0 | @ajlkn | MIT licensed */
var breakpoints=function(){"use strict";function e(e){t.init(e)}var t={list:null,media:{},events:[],init:function(e){t.list=e,window.addEventListener("resize",t.poll),window.addEventListener("orientationchange",t.poll),window.addEventListener("load",t.poll),window.addEventListener("fullscreenchange",t.poll)},active:function(e){var n,a,s,i,r,d,c;if(!(e in t.media)){if(">="==e.substr(0,2)?(a="gte",n=e.substr(2)):"<="==e.substr(0,2)?(a="lte",n=e.substr(2)):">"==e.substr(0,1)?(a="gt",n=e.substr(1)):"<"==e.substr(0,1)?(a="lt",n=e.substr(1)):"!"==e.substr(0,1)?(a="not",n=e.substr(1)):(a="eq",n=e),n&&n in t.list)if(i=t.list[n],Array.isArray(i)){if(r=parseInt(i[0]),d=parseInt(i[1]),isNaN(r)){if(isNaN(d))return;c=i[1].substr(String(d).length)}else c=i[0].substr(String(r).length);if(isNaN(r))switch(a){case"gte":s="screen";break;case"lte":s="screen and (max-width: "+d+c+")";break;case"gt":s="screen and (min-width: "+(d+1)+c+")";break;case"lt":s="screen and (max-width: -1px)";break;case"not":s="screen and (min-width: "+(d+1)+c+")";break;default:s="screen and (max-width: "+d+c+")"}else if(isNaN(d))switch(a){case"gte":s="screen and (min-width: "+r+c+")";break;case"lte":s="screen";break;case"gt":s="screen and (max-width: -1px)";break;case"lt":s="screen and (max-width: "+(r-1)+c+")";break;case"not":s="screen and (max-width: "+(r-1)+c+")";break;default:s="screen and (min-width: "+r+c+")"}else switch(a){case"gte":s="screen and (min-width: "+r+c+")";break;case"lte":s="screen and (max-width: "+d+c+")";break;case"gt":s="screen and (min-width: "+(d+1)+c+")";break;case"lt":s="screen and (max-width: "+(r-1)+c+")";break;case"not":s="screen and (max-width: "+(r-1)+c+"), screen and (min-width: "+(d+1)+c+")";break;default:s="screen and (min-width: "+r+c+") and (max-width: "+d+c+")"}}else s="("==i.charAt(0)?"screen and "+i:i;t.media[e]=!!s&&s}return t.media[e]!==!1&&window.matchMedia(t.media[e]).matches},on:function(e,n){t.events.push({query:e,handler:n,state:!1}),t.active(e)&&n()},poll:function(){var e,n;for(e=0;e<t.events.length;e++)n=t.events[e],t.active(n.query)?n.state||(n.state=!0,n.handler()):n.state&&(n.state=!1)}};return e._=t,e.on=function(e,n){t.on(e,n)},e.active=function(e){return t.active(e)},e}();!function(e,t){"function"==typeof define&&define.amd?define([],t):"object"==typeof exports?module.exports=t():e.breakpoints=t()}(this,function(){return breakpoints});
```
