## Extended Data File 1 for "Diet induces reproducible alterations in the mouse and human gut microbiome": browser.min.js

```
/* browser.js v1.0 | @ajlkn | MIT licensed */
var browser=function(){"use strict";var e={name:null,version:null,os:null,osVersion:null,touch:null,mobile:null,_canUse:null,canUse:function(n){e._canUse||(e._canUse=document.createElement("div"));var o=e._canUse.style,r=n.charAt(0).toUpperCase()+n.slice(1);return n in o||"Moz"+r in o||"Webkit"+r in o||"O"+r in o||"ms"+r in o},init:function(){var n,o,r,i,t=navigator.userAgent;for(n="other",o=0,r=[["firefox",/Firefox\/([0-9\.]+)/],["bb",/BlackBerry.+Version\/([0-9\.]+)/],["bb",/BB[0-9]+.+Version\/([0-9\.]+)/],["opera",/OPR\/([0-9\.]+)/],["opera",/Opera\/([0-9\.]+)/],["edge",/Edge\/([0-9\.]+)/],["safari",/Version\/([0-9\.]+).+Safari/],["chrome",/Chrome\/([0-9\.]+)/],["ie",/MSIE ([0-9]+)/],["ie",/Trident\/.+rv:([0-9]+)/]],i=0;i<r.length;i++)if(t.match(r[i][1])){n=r[i][0],o=parseFloat(RegExp.$1);break}for(e.name=n,e.version=o,n="other",o=0,r=[["ios",/([0-9_]+) like Mac OS X/,function(e){return e.replace("_",".").replace("_","")}],["ios",/CPU like Mac OS X/,function(e){return 0}],["wp",/Windows Phone ([0-9\.]+)/,null],["android",/Android ([0-9\.]+)/,null],["mac",/Macintosh.+Mac OS X ([0-9_]+)/,function(e){return e.replace("_",".").replace("_","")}],["windows",/Windows NT ([0-9\.]+)/,null],["bb",/BlackBerry.+Version\/([0-9\.]+)/,null],["bb",/BB[0-9]+.+Version\/([0-9\.]+)/,null],["linux",/Linux/,null],["bsd",/BSD/,null],["unix",/X11/,null]],i=0;i<r.length;i++)if(t.match(r[i][1])){n=r[i][0],o=parseFloat(r[i][2]?r[i][2](RegExp.$1):RegExp.$1);break}e.os=n,e.osVersion=o,e.touch="wp"==e.os?navigator.msMaxTouchPoints>0:!!("ontouchstart"in window),e.mobile="wp"==e.os||"android"==e.os||"ios"==e.os||"bb"==e.os}};return e.init(),e}();!function(e,n){"function"==typeof define&&define.amd?define([],n):"object"==typeof exports?module.exports=n():e.browser=n()}(this,function(){return browser});
```
