## Extended Data File 1 for "Diet induces reproducible alterations in the mouse and human gut microbiome": jquery.scrolly.min.js

```
/* jquery.scrolly v1.0.0-dev | (c) @ajlkn | MIT licensed */
(function(e){function u(s,o){var u,a,f;if((u=e(s))[t]==0)return n;a=u[i]()[r];switch(o.anchor){case"middle":f=a-(e(window).height()-u.outerHeight())/2;break;default:case r:f=Math.max(a,0)}return typeof o[i]=="function"?f-=o[i]():f-=o[i],f}var t="length",n=null,r="top",i="offset",s="click.scrolly",o=e(window);e.fn.scrolly=function(i){var o,a,f,l,c=e(this);if(this[t]==0)return c;if(this[t]>1){for(o=0;o<this[t];o++)e(this[o]).scrolly(i);return c}l=n,f=c.attr("href");if(f.charAt(0)!="#"||f[t]<2)return c;a=jQuery.extend({anchor:r,easing:"swing",offset:0,parent:e("body,html"),pollOnce:!1,speed:1e3},i),a.pollOnce&&(l=u(f,a)),c.off(s).on(s,function(e){var t=l!==n?l:u(f,a);t!==n&&(e.preventDefault(),a.parent.stop().animate({scrollTop:t},a.speed,a.easing))})}})(jQuery);
```
