## Extended Data File 1 for "Diet induces reproducible alterations in the mouse and human gut microbiome": main.js

```
/*
	Photon by HTML5 UP
	html5up.net | @ajlkn
	Free for personal and commercial use under the CCA 3.0 license (html5up.net/license)
*/

(function($) {

	var	$window = $(window),
		$body = $('body');

	// Breakpoints.
		breakpoints({
			xlarge:   [ '1141px',  '1680px' ],
			large:    [ '981px',   '1140px' ],
			medium:   [ '737px',   '980px'  ],
			small:    [ '481px',   '736px'  ],
			xsmall:   [ '321px',   '480px'  ],
			xxsmall:  [ null,      '320px'  ]
		});

	// Play initial animations on page load.
		$window.on('load', function() {
			window.setTimeout(function() {
				$body.removeClass('is-preload');
			}, 100);
		});

	// Scrolly.
		$('.scrolly').scrolly();

})(jQuery);
```
