## Extended Data File 1 for "Diet induces reproducible alterations in the mouse and human gut microbiome": util.js

```
(function($) {

	/**
	 * Generate an indented list of links from a nav. Meant for use with panel().
	 * @return {jQuery} jQuery object.
	 */
	$.fn.navList = function() {

		var	$this = $(this);
			$a = $this.find('a'),
			b = [];

		$a.each(function() {

			var	$this = $(this),
				indent = Math.max(0, $this.parents('li').length - 1),
				href = $this.attr('href'),
				target = $this.attr('target');

			b.push(
				'<a ' +
					'class="link depth-' + indent + '"' +
					( (typeof target !== 'undefined' && target != '') ? ' target="' + target + '"' : '') +
					( (typeof href !== 'undefined' && href != '') ? ' href="' + href + '"' : '') +
				'>' +
					'<span class="indent-' + indent + '"></span>' +
					$this.text() +
				'</a>'
			);

		});

		return b.join('');

	};

	/**
	 * Panel-ify an element.
	 * @param {object} userConfig User config.
	 * @return {jQuery} jQuery object.
	 */
	$.fn.panel = function(userConfig) {

		// No elements?
			if (this.length == 0)
				return $this;

		// Multiple elements?
			if (this.length > 1) {

				for (var i=0; i < this.length; i++)
					$(this[i]).panel(userConfig);

				return $this;

			}

		// Vars.
			var	$this = $(this),
				$body = $('body'),
				$window = $(window),
				id = $this.attr('id'),
				config;

		// Config.
			config = $.extend({

				// Delay.
					delay: 0,

				// Hide panel on link click.
					hideOnClick: false,

				// Hide panel on escape keypress.
					hideOnEscape: false,

				// Hide panel on swipe.
					hideOnSwipe: false,

				// Reset scroll position on hide.
					resetScroll: false,

				// Reset forms on hide.
					resetForms: false,

				// Side of viewport the panel will appear.
					side: null,

				// Target element for "class".
					target: $this,

				// Class to toggle.
					visibleClass: 'visible'

			}, userConfig);

			// Expand "target" if it's not a jQuery object already.
				if (typeof config.target != 'jQuery')
					config.target = $(config.target);

		// Panel.

			// Methods.
				$this._hide = function(event) {

					// Already hidden? Bail.
						if (!config.target.hasClass(config.visibleClass))
							return;

					// If an event was provided, cancel it.
						if (event) {

							event.preventDefault();
							event.stopPropagation();

						}

					// Hide.
						config.target.removeClass(config.visibleClass);

					// Post-hide stuff.
						window.setTimeout(function() {

							// Reset scroll position.
								if (config.resetScroll)
									$this.scrollTop(0);

							// Reset forms.
								if (config.resetForms)
									$this.find('form').each(function() {
										this.reset();
									});

						}, config.delay);

				};

			// Vendor fixes.
				$this
					.css('-ms-overflow-style', '-ms-autohiding-scrollbar')
					.css('-webkit-overflow-scrolling', 'touch');

			// Hide on click.
				if (config.hideOnClick) {

					$this.find('a')
						.css('-webkit-tap-highlight-color', 'rgba(0,0,0,0)');

					$this
						.on('click', 'a', function(event) {

							var $a = $(this),
								href = $a.attr('href'),
								target = $a.attr('target');

							if (!href || href == '#' || href == '' || href == '#' + id)
								return;

							// Cancel original event.
								event.preventDefault();
								event.stopPropagation();

							// Hide panel.
								$this._hide();

							// Redirect to href.
								window.setTimeout(function() {

									if (target == '_blank')
										window.open(href);
									else
										window.location.href = href;

								}, config.delay + 10);

						});

				}

			// Event: Touch stuff.
				$this.on('touchstart', function(event) {

					$this.touchPosX = event.originalEvent.touches[0].pageX;
					$this.touchPosY = event.originalEvent.touches[0].pageY;

				})

				$this.on('touchmove', function(event) {

					if ($this.touchPosX === null
					||	$this.touchPosY === null)
						return;

					var	diffX = $this.touchPosX - event.originalEvent.touches[0].pageX,
						diffY = $this.touchPosY - event.originalEvent.touches[0].pageY,
						th = $this.outerHeight(),
						ts = ($this.get(0).scrollHeight - $this.scrollTop());

					// Hide on swipe?
						if (config.hideOnSwipe) {

							var result = false,
								boundary = 20,
								delta = 50;

							switch (config.side) {

								case 'left':
									result = (diffY < boundary && diffY > (-1 * boundary)) && (diffX > delta);
									break;

								case 'right':
									result = (diffY < boundary && diffY > (-1 * boundary)) && (diffX < (-1 * delta));
									break;

								case 'top':
									result = (diffX < boundary && diffX > (-1 * boundary)) && (diffY > delta);
									break;

								case 'bottom':
									result = (diffX < boundary && diffX > (-1 * boundary)) && (diffY < (-1 * delta));
									break;

								default:
									break;

							}

							if (result) {

								$this.touchPosX = null;
								$this.touchPosY = null;
								$this._hide();

								return false;

							}

						}

					// Prevent vertical scrolling past the top or bottom.
						if (($this.scrollTop() < 0 && diffY < 0)
						|| (ts > (th - 2) && ts < (th + 2) && diffY > 0)) {

							event.preventDefault();
							event.stopPropagation();

						}

				});

			// Event: Prevent certain events inside the panel from bubbling.
				$this.on('click touchend touchstart touchmove', function(event) {
					event.stopPropagation();
				});

			// Event: Hide panel if a child anchor tag pointing to its ID is clicked.
				$this.on('click', 'a[href="#' + id + '"]', function(event) {

					event.preventDefault();
					event.stopPropagation();

					config.target.removeClass(config.visibleClass);

				});

		// Body.

			// Event: Hide panel on body click/tap.
				$body.on('click touchend', function(event) {
					$this._hide(event);
				});

			// Event: Toggle.
				$body.on('click', 'a[href="#' + id + '"]', function(event) {

					event.preventDefault();
					event.stopPropagation();

					config.target.toggleClass(config.visibleClass);

				});

		// Window.

			// Event: Hide on ESC.
				if (config.hideOnEscape)
					$window.on('keydown', function(event) {

						if (event.keyCode == 27)
							$this._hide(event);

					});

		return $this;

	};

	/**
	 * Apply "placeholder" attribute polyfill to one or more forms.
	 * @return {jQuery} jQuery object.
	 */
	$.fn.placeholder = function() {

		// Browser natively supports placeholders? Bail.
			if (typeof (document.createElement('input')).placeholder != 'undefined')
				return $(this);

		// No elements?
			if (this.length == 0)
				return $this;

		// Multiple elements?
			if (this.length > 1) {

				for (var i=0; i < this.length; i++)
					$(this[i]).placeholder();

				return $this;

			}

		// Vars.
			var $this = $(this);

		// Text, TextArea.
			$this.find('input[type=text],textarea')
				.each(function() {

					var i = $(this);

					if (i.val() == ''
					||  i.val() == i.attr('placeholder'))
						i
							.addClass('polyfill-placeholder')
							.val(i.attr('placeholder'));

				})
				.on('blur', function() {

					var i = $(this);

					if (i.attr('name').match(/-polyfill-field$/))
						return;

					if (i.val() == '')
						i
							.addClass('polyfill-placeholder')
							.val(i.attr('placeholder'));

				})
				.on('focus', function() {

					var i = $(this);

					if (i.attr('name').match(/-polyfill-field$/))
						return;

					if (i.val() == i.attr('placeholder'))
						i
							.removeClass('polyfill-placeholder')
							.val('');

				});

		// Password.
			$this.find('input[type=password]')
				.each(function() {

					var i = $(this);
					var x = $(
								$('<div>')
									.append(i.clone())
									.remove()
									.html()
									.replace(/type="password"/i, 'type="text"')
									.replace(/type=password/i, 'type=text')
					);

					if (i.attr('id') != '')
						x.attr('id', i.attr('id') + '-polyfill-field');

					if (i.attr('name') != '')
						x.attr('name', i.attr('name') + '-polyfill-field');

					x.addClass('polyfill-placeholder')
						.val(x.attr('placeholder')).insertAfter(i);

					if (i.val() == '')
						i.hide();
					else
						x.hide();

					i
						.on('blur', function(event) {

							event.preventDefault();

							var x = i.parent().find('input[name=' + i.attr('name') + '-polyfill-field]');

							if (i.val() == '') {

								i.hide();
								x.show();

							}

						});

					x
						.on('focus', function(event) {

							event.preventDefault();

							var i = x.parent().find('input[name=' + x.attr('name').replace('-polyfill-field', '') + ']');

							x.hide();

							i
								.show()
								.focus();

						})
						.on('keypress', function(event) {

							event.preventDefault();
							x.val('');

						});

				});

		// Events.
			$this
				.on('submit', function() {

					$this.find('input[type=text],input[type=password],textarea')
						.each(function(event) {

							var i = $(this);

							if (i.attr('name').match(/-polyfill-field$/))
								i.attr('name', '');

							if (i.val() == i.attr('placeholder')) {

								i.removeClass('polyfill-placeholder');
								i.val('');

							}

						});

				})
				.on('reset', function(event) {

					event.preventDefault();

					$this.find('select')
						.val($('option:first').val());

					$this.find('input,textarea')
						.each(function() {

							var i = $(this),
								x;

							i.removeClass('polyfill-placeholder');

							switch (this.type) {

								case 'submit':
								case 'reset':
									break;

								case 'password':
									i.val(i.attr('defaultValue'));

									x = i.parent().find('input[name=' + i.attr('name') + '-polyfill-field]');

									if (i.val() == '') {
										i.hide();
										x.show();
									}
									else {
										i.show();
										x.hide();
									}

									break;

								case 'checkbox':
								case 'radio':
									i.attr('checked', i.attr('defaultValue'));
									break;

								case 'text':
								case 'textarea':
									i.val(i.attr('defaultValue'));

									if (i.val() == '') {
										i.addClass('polyfill-placeholder');
										i.val(i.attr('placeholder'));
									}

									break;

								default:
									i.val(i.attr('defaultValue'));
									break;

							}
						});

				});

		return $this;

	};

	/**
	 * Moves elements to/from the first positions of their respective parents.
	 * @param {jQuery} $elements Elements (or selector) to move.
	 * @param {bool} condition If true, moves elements to the top. Otherwise, moves elements back to their original locations.
	 */
	$.prioritize = function($elements, condition) {

		var key = '__prioritize';

		// Expand $elements if it's not already a jQuery object.
			if (typeof $elements != 'jQuery')
				$elements = $($elements);

		// Step through elements.
			$elements.each(function() {

				var	$e = $(this), $p,
					$parent = $e.parent();

				// No parent? Bail.
					if ($parent.length == 0)
						return;

				// Not moved? Move it.
					if (!$e.data(key)) {

						// Condition is false? Bail.
							if (!condition)
								return;

						// Get placeholder (which will serve as our point of reference for when this element needs to move back).
							$p = $e.prev();

							// Couldn't find anything? Means this element's already at the top, so bail.
								if ($p.length == 0)
									return;

						// Move element to top of parent.
							$e.prependTo($parent);

						// Mark element as moved.
							$e.data(key, $p);

					}

				// Moved already?
					else {

						// Condition is true? Bail.
							if (condition)
								return;

						$p = $e.data(key);

						// Move element back to its original location (using our placeholder).
							$e.insertAfter($p);

						// Unmark element as moved.
							$e.removeData(key);

					}

			});

	};

})(jQuery);
```
