## Extended Data File 1 for "Diet induces reproducible alterations in the mouse and human gut microbiome": LICENSE.txt

Creative Commons Attribution 3.0 Unported
http://creativecommons.org/licenses/by/3.0/

License

THE WORK (AS DEFINED BELOW) IS PROVIDED UNDER THE TERMS OF THIS CREATIVE COMMONS PUBLIC LICENSE ("CCPL" OR "LICENSE"). THE WORK IS PROTECTED BY COPYRIGHT AND/OR OTHER APPLICABLE LAW. ANY USE OF THE WORK OTHER THAN AS AUTHORIZED UNDER THIS LICENSE OR COPYRIGHT LAW IS PROHIBITED.

BY EXERCISING ANY RIGHTS TO THE WORK PROVIDED HERE, YOU ACCEPT AND AGREE TO BE BOUND BY THE TERMS OF THIS LICENSE. TO THE EXTENT THIS LICENSE MAY BE CONSIDERED TO BE A CONTRACT, THE LICENSOR GRANTS YOU THE RIGHTS CONTAINED HERE IN CONSIDERATION OF YOUR ACCEPTANCE OF SUCH TERMS AND CONDITIONS.

1. Definitions

   1. "Adaptation" means a work based upon the Work, or upon the Work and other pre-existing works, such as a translation, adaptation, derivative work, arrangement of music or other alterations of a literary or artistic work, or phonogram or performance and includes cinematographic adaptations or any other form in which the Work may be recast, transformed, or adapted including in any form recognizably derived from the original, except that a work that constitutes a Collection will not be considered an Adaptation for the purpose of this License. For the avoidance of doubt, where the Work is a musical work, performance or phonogram, the synchronization of the Work in timed-relation with a moving image ("synching") will be considered an Adaptation for the purpose of this License.
   2. "Collection" means a collection of literary or artistic works, such as encyclopedias and anthologies, or performances, phonograms or broadcasts, or other works or subject matter other than works listed in Section 1(f) below, which, by reason of the selection and arrangement of their contents, constitute intellectual creations, in which the Work is included in its entirety in unmodified form along with one or more other contributions, each constituting separate and independent works in themselves, which together are assembled into a collective whole. A work that constitutes a Collection will not be considered an Adaptation (as defined above) for the purposes of this License.
   3. "Distribute" means to make available to the public the original and copies of the Work or Adaptation, as appropriate, through sale or other transfer of ownership.
   4. "Licensor" means the individual, individuals, entity or entities that offer(s) the Work under the terms of this License.
   5. "Original Author" means, in the case of a literary or artistic work, the individual, individuals, entity or entities who created the Work or if no individual or entity can be identified, the publisher; and in addition (i) in the case of a performance the actors, singers, musicians, dancers, and other persons who act, sing, deliver, declaim, play in, interpret or otherwise perform literary or artistic works or expressions of folklore; (ii) in the case of a phonogram the producer being the person or legal entity who first fixes the sounds of a performance or other sounds; and, (iii) in the case of broadcasts, the organization that transmits the broadcast.
   6. "Work" means the literary and/or artistic work offered under the terms of this License including without limitation any production in the literary, scientific and artistic domain, whatever may be the mode or form of its expression including digital form, such as a book, pamphlet and other writing; a lecture, address, sermon or other work of the same nature; a dramatic or dramatico-musical work; a choreographic work or entertainment in dumb show; a musical composition with or without words; a cinematographic work to which are assimilated works expressed by a process analogous to cinematography; a work of drawing, painting, architecture, sculpture, engraving or lithography; a photographic work to which are assimilated works expressed by a process analogous to photography; a work of applied art; an illustration, map, plan, sketch or three-dimensional work relative to geography, topography, architecture or science; a performance; a broadcast; a phonogram; a compilation of data to the extent it is protected as a copyrightable work; or a work performed by a variety or circus performer to the extent it is not otherwise considered a literary or artistic work.
   7. "You" means an individual or entity exercising rights under this License who has not previously violated the terms of this License with respect to the Work, or who has received express permission from the Licensor to exercise rights under this License despite a previous violation.
   8. "Publicly Perform" means to perform public recitations of the Work and to communicate to the public those public recitations, by any means or process, including by wire or wireless means or public digital performances; to make available to the public Works in such a way that members of the public may access these Works from a place and at a place individually chosen by them; to perform the Work to the public by any means or process and the communication to the public of the performances of the Work, including by public digital performance; to broadcast and rebroadcast the Work by any means including signs, sounds or images.
   9. "Reproduce" means to make copies of the Work by any means including without limitation by sound or visual recordings and the right of fixation and reproducing fixations of the Work, including storage of a protected performance or phonogram in digital form or other electronic medium.

2. Fair Dealing Rights. Nothing in this License is intended to reduce, limit, or restrict any uses free from copyright or rights arising from limitations or exceptions that are provided for in connection with the copyright protection under copyright law or other applicable laws.

3. License Grant. Subject to the terms and conditions of this License, Licensor hereby grants You a worldwide, royalty-free, non-exclusive, perpetual (for the duration of the applicable copyright) license to exercise the rights in the Work as stated below:

   1. to Reproduce the Work, to incorporate the Work into one or more Collections, and to Reproduce the Work as incorporated in the Collections;
   2. to create and Reproduce Adaptations provided that any such Adaptation, including any translation in any medium, takes reasonable steps to clearly label, demarcate or otherwise identify that changes were made to the original Work. For example, a translation could be marked "The original work was translated from English to Spanish," or a modification could indicate "The original work has been modified.";
   3. to Distribute and Publicly Perform the Work including as incorporated in Collections; and,
   4. to Distribute and Publicly Perform Adaptations.
   5.

      For the avoidance of doubt:
         1. Non-waivable Compulsory License Schemes. In those jurisdictions in which the right to collect royalties through any statutory or compulsory licensing scheme cannot be waived, the Licensor reserves the exclusive right to collect such royalties for any exercise by You of the rights granted under this License;
         2. Waivable Compulsory License Schemes. In those jurisdictions in which the right to collect royalties through any statutory or compulsory licensing scheme can be waived, the Licensor waives the exclusive right to collect such royalties for any exercise by You of the rights granted under this License; and,
         3. Voluntary License Schemes. The Licensor waives the right to collect royalties, whether individually or, in the event that the Licensor is a member of a collecting society that administers voluntary licensing schemes, via that society, from any exercise by You of the rights granted under this License.

The above rights may be exercised in all media and formats whether now known or hereafter devised. The above rights include the right to make such modifications as are technically necessary to exercise the rights in other media and formats. Subject to Section 8(f), all rights not expressly granted by Licensor are hereby reserved.

4. Restrictions. The license granted in Section 3 above is expressly made subject to and limited by the following restrictions:

   1. You may Distribute or Publicly Perform the Work only under the terms of this License. You must include a copy of, or the Uniform Resource Identifier (URI) for, this License with every copy of the Work You Distribute or Publicly Perform. You may not offer or impose any terms on the Work that restrict the terms of this License or the ability of the recipient of the Work to exercise the rights granted to that recipient under the terms of the License. You may not sublicense the Work. You must keep intact all notices that refer to this License and to the disclaimer of warranties with every copy of the Work You Distribute or Publicly Perform. When You Distribute or Publicly Perform the Work, You may not impose any effective technological measures on the Work that restrict the ability of a recipient of the Work from You to exercise the rights granted to that recipient under the terms of the License. This Section 4(a) applies to the Work as incorporated in a Collection, but this does not require the Collection apart from the Work itself to be made subject to the terms of this License. If You create a Collection, upon notice from any Licensor You must, to the extent practicable, remove from the Collection any credit as required by Section 4(b), as requested. If You create an Adaptation, upon notice from any Licensor You must, to the extent practicable, remove from the Adaptation any credit as required by Section 4(b), as requested.
   2. If You Distribute, or Publicly Perform the Work or any Adaptations or Collections, You must, unless a request has been made pursuant to Section 4(a), keep intact all copyright notices for the Work and provide, reasonable to the medium or means You are utilizing: (i) the name of the Original Author (or pseudonym, if applicable) if supplied, and/or if the Original Author and/or Licensor designate another party or parties (e.g., a sponsor institute, publishing entity, journal) for attribution ("Attribution Parties") in Licensor's copyright notice, terms of service or by other reasonable means, the name of such party or parties; (ii) the title of the Work if supplied; (iii) to the extent reasonably practicable, the URI, if any, that Licensor specifies to be associated with the Work, unless such URI does not refer to the copyright notice or licensing information for the Work; and (iv) , consistent with Section 3(b), in the case of an Adaptation, a credit identifying the use of the Work in the Adaptation (e.g., "French translation of the Work by Original Author," or "Screenplay based on original Work by Original Author"). The credit required by this Section 4 (b) may be implemented in any reasonable manner; provided, however, that in the case of a Adaptation or Collection, at a minimum such credit will appear, if a credit for all contributing authors of the Adaptation or Collection appears, then as part of these credits and in a manner at least as prominent as the credits for the other contributing authors. For the avoidance of doubt, You may only use the credit required by this Section for the purpose of attribution in the manner set out above and, by exercising Your rights under this License, You may not implicitly or explicitly assert or imply any connection with, sponsorship or endorsement by the Original Author, Licensor and/or Attribution Parties, as appropriate, of You or Your use of the Work, without the separate, express prior written permission of the Original Author, Licensor and/or Attribution Parties.
   3. Except as otherwise agreed in writing by the Licensor or as may be otherwise permitted by applicable law, if You Reproduce, Distribute or Publicly Perform the Work either by itself or as part of any Adaptations or Collections, You must not distort, mutilate, modify or take other derogatory action in relation to the Work which would be prejudicial to the Original Author's honor or reputation. Licensor agrees that in those jurisdictions (e.g. Japan), in which any exercise of the right granted in Section 3(b) of this License (the right to make Adaptations) would be deemed to be a distortion, mutilation, modification or other derogatory action prejudicial to the Original Author's honor and reputation, the Licensor will waive or not assert, as appropriate, this Section, to the fullest extent permitted by the applicable national law, to enable You to reasonably exercise Your right under Section 3(b) of this License (right to make Adaptations) but not otherwise.

5. Representations, Warranties and Disclaimer

UNLESS OTHERWISE MUTUALLY AGREED TO BY THE PARTIES IN WRITING, LICENSOR OFFERS THE WORK AS-IS AND MAKES NO REPRESENTATIONS OR WARRANTIES OF ANY KIND CONCERNING THE WORK, EXPRESS, IMPLIED, STATUTORY OR OTHERWISE, INCLUDING, WITHOUT LIMITATION, WARRANTIES OF TITLE, MERCHANTIBILITY, FITNESS FOR A PARTICULAR PURPOSE, NONINFRINGEMENT, OR THE ABSENCE OF LATENT OR OTHER DEFECTS, ACCURACY, OR THE PRESENCE OF ABSENCE OF ERRORS, WHETHER OR NOT DISCOVERABLE. SOME JURISDICTIONS DO NOT ALLOW THE EXCLUSION OF IMPLIED WARRANTIES, SO SUCH EXCLUSION MAY NOT APPLY TO YOU.

6. Limitation on Liability. EXCEPT TO THE EXTENT REQUIRED BY APPLICABLE LAW, IN NO EVENT WILL LICENSOR BE LIABLE TO YOU ON ANY LEGAL THEORY FOR ANY SPECIAL, INCIDENTAL, CONSEQUENTIAL, PUNITIVE OR EXEMPLARY DAMAGES ARISING OUT OF THIS LICENSE OR THE USE OF THE WORK, EVEN IF LICENSOR HAS BEEN ADVISED OF THE POSSIBILITY OF SUCH DAMAGES.

7. Termination

   1. This License and the rights granted hereunder will terminate automatically upon any breach by You of the terms of this License. Individuals or entities who have received Adaptations or Collections from You under this License, however, will not have their licenses terminated provided such individuals or entities remain in full compliance with those licenses. Sections 1, 2, 5, 6, 7, and 8 will survive any termination of this License.
   2. Subject to the above terms and conditions, the license granted here is perpetual (for the duration of the applicable copyright in the Work). Notwithstanding the above, Licensor reserves the right to release the Work under different license terms or to stop distributing the Work at any time; provided, however that any such election will not serve to withdraw this License (or any other license that has been, or is required to be, granted under the terms of this License), and this License will continue in full force and effect unless terminated as stated above.

8. Miscellaneous

   1. Each time You Distribute or Publicly Perform the Work or a Collection, the Licensor offers to the recipient a license to the Work on the same terms and conditions as the license granted to You under this License.
   2. Each time You Distribute or Publicly Perform an Adaptation, Licensor offers to the recipient a license to the original Work on the same terms and conditions as the license granted to You under this License.
   3. If any provision of this License is invalid or unenforceable under applicable law, it shall not affect the validity or enforceability of the remainder of the terms of this License, and without further action by the parties to this agreement, such provision shall be reformed to the minimum extent necessary to make such provision valid and enforceable.
   4. No term or provision of this License shall be deemed waived and no breach consented to unless such waiver or consent shall be in writing and signed by the party to be charged with such waiver or consent.
   5. This License constitutes the entire agreement between the parties with respect to the Work licensed here. There are no understandings, agreements or representations with respect to the Work not specified here. Licensor shall not be bound by any additional provisions that may appear in any communication from You. This License may not be modified without the mutual written agreement of the Licensor and You.
   6. The rights granted under, and the subject matter referenced, in this License were drafted utilizing the terminology of the Berne Convention for the Protection of Literary and Artistic Works (as amended on September 28, 1979), the Rome Convention of 1961, the WIPO Copyright Treaty of 1996, the WIPO Performances and Phonograms Treaty of 1996 and the Universal Copyright Convention (as revised on July 24, 1971). These rights and subject matter take effect in the relevant jurisdiction in which the License terms are sought to be enforced according to the corresponding provisions of the implementation of those treaty provisions in the applicable national law. If the standard suite of rights granted under applicable copyright law includes additional rights not granted under this License, such additional rights are deemed to be included in the License; this License is not intended to restrict the license of any rights under applicable law.
