## Extended Data File 1 for "Diet induces reproducible alterations in the mouse and human gut microbiome": README.txt

The following directory contains the data and code necessary to recapitulate the analysis in our manuscript. Please start with index.html if you have downloaded this directory.


##############################################

Please see below for index template information:

Photon by HTML5 UP
html5up.net | @ajlkn
Free for personal and commercial use under the CCA 3.0 license (html5up.net/license)


A simple (gradient-heavy) single pager that revisits a style I messed with on two
previous designs (Tessellate and Telephasic). Fully responsive, built on Sass,
and, as usual, loaded with an assortment of pre-styled elements. Have fun! :)

Demo images* courtesy of Unsplash, a radtastic collection of CC0 (public domain) images
you can use for pretty much whatever.

(* = Not included)

Feedback, bug reports, and comments are not only welcome, but strongly encouraged :)

AJ
 | @ajlkn


Credits:

	Demo Images:
		Unsplash (unsplash.com)

	Icons:
		Font Awesome (fontawesome.io)

	Other:
		jQuery (jquery.com)
		Responsive Tools (github.com/ajlkn/responsive-tools)
