## Supplementary figures and images for "Diet induces reproducible alterations in the mouse and human gut microbiome"

### cml.png

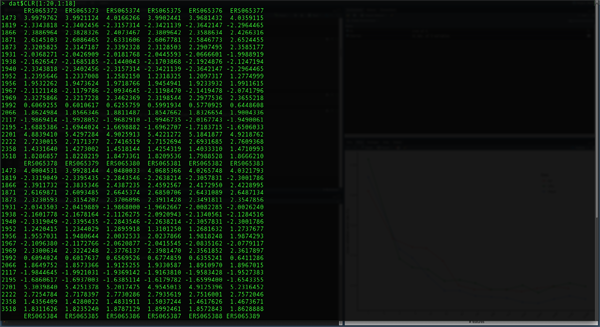

### header.jpg

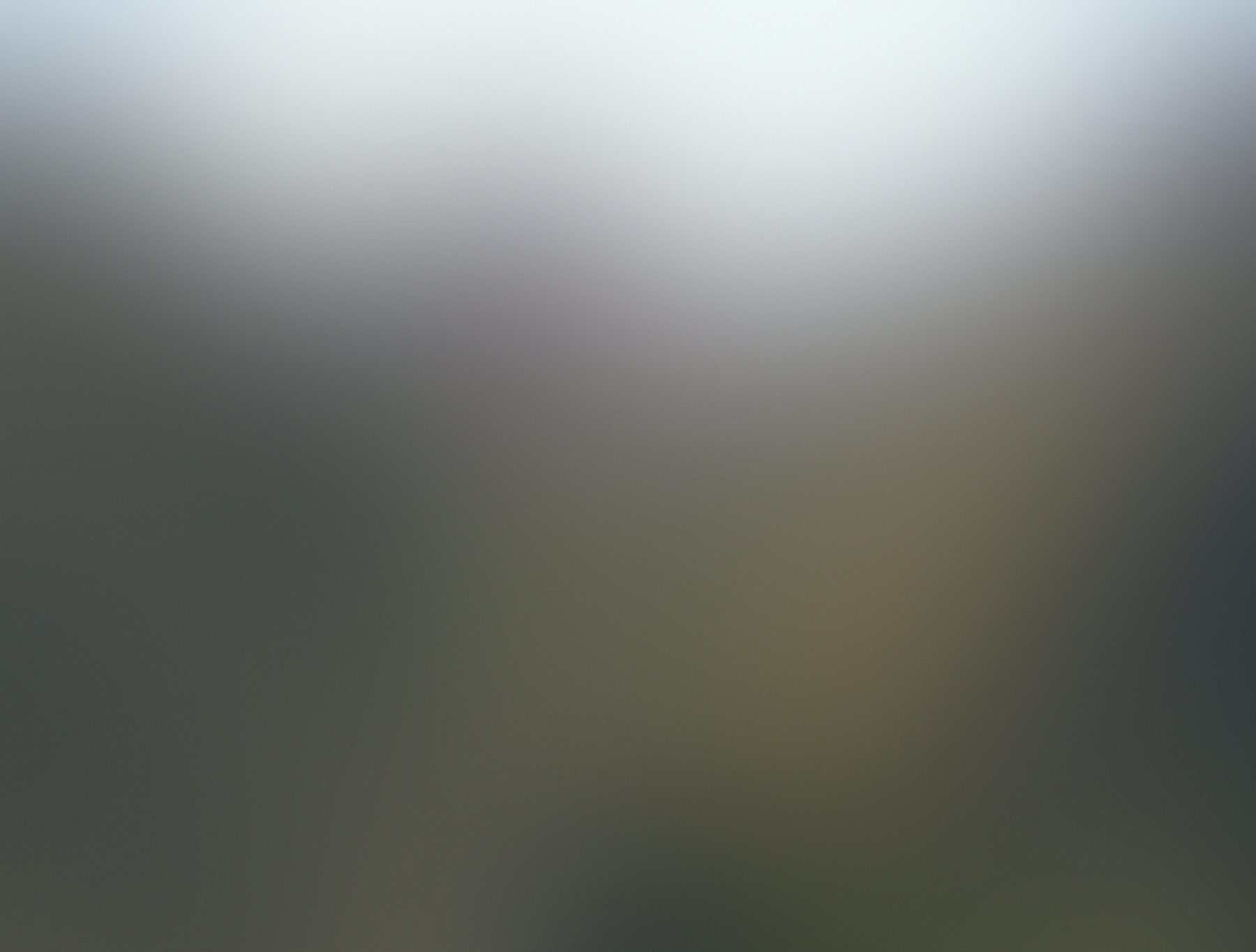

### overlay1.png

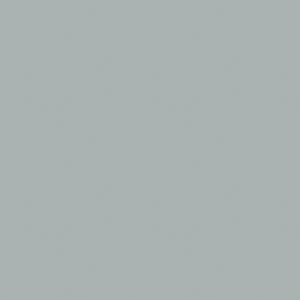

### overlay2.png

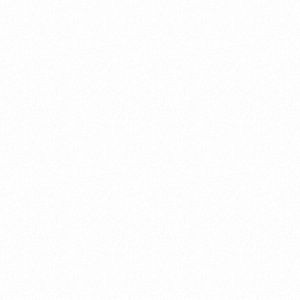

### rs.png

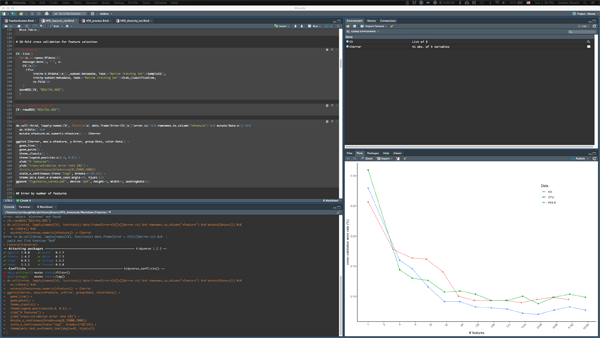
